# Single-deletion-mutant, third-generation rabies viral vectors allow nontoxic retrograde targeting of projection neurons with greatly increased efficiency

**DOI:** 10.1101/2022.02.23.481706

**Authors:** Lei Jin, Heather A. Sullivan, Mulangma Zhu, Nicholas E. Lea, Thomas K. Lavin, Xin Fu, Makoto Matsuyama, YuanYuan Hou, Guoping Feng, Ian R. Wickersham

## Abstract

Rabies viral vectors have become important components of the systems neuroscience toolkit, allowing both direct retrograde targeting of projection neurons and monosynaptic tracing of inputs to defined postsynaptic populations, but the rapid cytotoxicity of first-generation (ΔG) vectors limits their use to short-term experiments. We recently introduced second-generation, double-deletion-mutant (ΔGL) rabies viral vectors, showing that they efficiently retrogradely infect projection neurons and express recombinases effectively but with little to no detectable toxicity; more recently, we have shown that ΔGL viruses can be used for monosynaptic tracing with far lower cytotoxicity than the first-generation system. Here we introduce third-generation (ΔL) rabies viral vectors, which, like first-generation vectors, have only a single gene deleted from their genomes (in this case the viral polymerase gene L) but which appear to be as nontoxic as second-generation ones, based on *in vivo* longitudinal structural and functional two-photon imaging and *ex vivo* electrophysiology. Although third-generation vectors are therefore phenotypically very similar to second-generation ones, we show that they have the major advantage of growing to much higher titers, and this key difference results in significantly increased numbers of retrogradely labeled neurons *in vivo*. These ΔL rabies viral vectors therefore constitute a new state of the art for minimally perturbative, pathway-specific expression of recombinases and transactivators in mammalian neurons selected on the basis of their axonal projections. Because replication of deletion-mutant rabies viruses within complementing cells is precisely the process that underlies monosynaptic tracing, the higher replication efficiency of this new class of rabies viral vectors furthermore suggests the potential to provide the foundation of an improved nontoxic monosynaptic tracing system.

## INTRODUCTION

Since their introduction to neuroscience in 2007 (Wickersham et al., 2007a; Wickersham et al., 2007b), recombinant rabies viral vectors have become widely-adopted tools in neuroscience, allowing “monosynaptic tracing” of direct inputs to genetically-targeted starting postsynaptic neuronal populations (Jin et al., 2021; Wall et al., 2010; Wickersham *et al*., 2007b) as well as simple retrograde targeting of projection neurons when injected at the sites of these projection neurons’ axonal arborizations (Chatterjee et al., 2018; Wickersham *et al*., 2007a). These vectors are now used in a large number of laboratories worldwide and have contributed to many high-impact studies of a wide variety of neural systems (Foster et al., 2021; Miyamichi et al., 2011; Reardon et al., 2016; Schwarz et al., 2015; Siu et al., 2021; Smith et al., 2021; Stephenson-Jones et al., 2016; Wu et al., 2021; Yao et al., 2021).

Because of the cytotoxicity of first-generation (“ΔG”) rabies viral vectors, which have only the glycoprotein gene G deleted from their genomes (Chatterjee *et al*., 2018; Jin *et al*., 2021; Wickersham *et al*., 2007a), we recently introduced second-generation, “ΔGL” rabies viral vectors, which have both G and the viral polymerase gene L (for “large” protein) deleted from their genomes (Chatterjee *et al*., 2018). The viral polymerase is absolutely required for transcription of all genes from the rabies viral genome as well as for replication of the viral genome itself (Albertini et al., 2011; Finke and Conzelmann, 2005; Horwitz et al., 2020; Morin et al., 2013; Ogino and Green, 2019; Te Velthuis et al., 2021). This additional deletion, by design, therefore reduces gene expression to a minimal level (provided by the few starting copies of the polymerase protein that are copackaged in each viral particle) that appears to be harmless to the “infected” cells. Because transgene expression is reduced by the same degree, we inserted the genes for Cre and Flpo recombinases, expression of which even at low levels is sufficient to cause neuroscientifically-useful downstream effects such as expression of fluorophores or calcium indicators in labeled cells (Chatterjee *et al*., 2018). We originally showed that these ΔGL vectors are useful tools for retrograde targeting of projection neurons (Chatterjee *et al*., 2018), and they have since been used as such for applications including optogenetics and transcriptomic profiling (Ren et al., 2021; Roy et al., 2021; Tasic et al., 2018). More recently, we have also shown that ΔGL vectors can be complemented *in vivo* by expression of both G and L *in trans*, yielding a second-generation monosynaptic tracing system with far lower cytotoxicity than the first-generation version (Jin *et al*., 2021).

Here we show that deletion of L alone appears to make rabies viral vectors as nontoxic as ΔGL ones, with labeled neurons surviving for at least months with apparently unperturbed visual response properties. We find that these ΔL vectors have a major growth advantage over ΔGL ones in cell culture, attaining much higher titers in complementing cells in culture. This higher replication efficiency translates into the practical advantage of retrogradely labeling many more projection neurons when injected into these neurons’ target sites *in vivo*.

## RESULTS

### Construction and characterization of ΔL rabies virus

We began by constructing rabies viral vectors with only the polymerase gene deleted and characterizing their gene expression levels and growth dynamics in cell culture (Fig. 1). Beginning with the genome plasmid of a ΔGL virus (Chatterjee *et al*., 2018), we reinserted the native glycoprotein gene into its original location, followed by the gene for Cre recombinase (codon-optimized for mouse (Koresawa et al., 2000)) in an additional transcriptional unit, then produced infectious virus by standard techniques (see Methods). We then compared the gene expression levels of the resulting virus, RVΔL-Cre, to those of first- and second-generation versions (RVΔG-Cre and RVΔGL-Cre, respectively) in cell culture (HEK 293T/17 cells) using immunostaining for Cre as well as for the viral nucleoprotein, the highest-expressed rabies viral protein.

**Figure 1.**
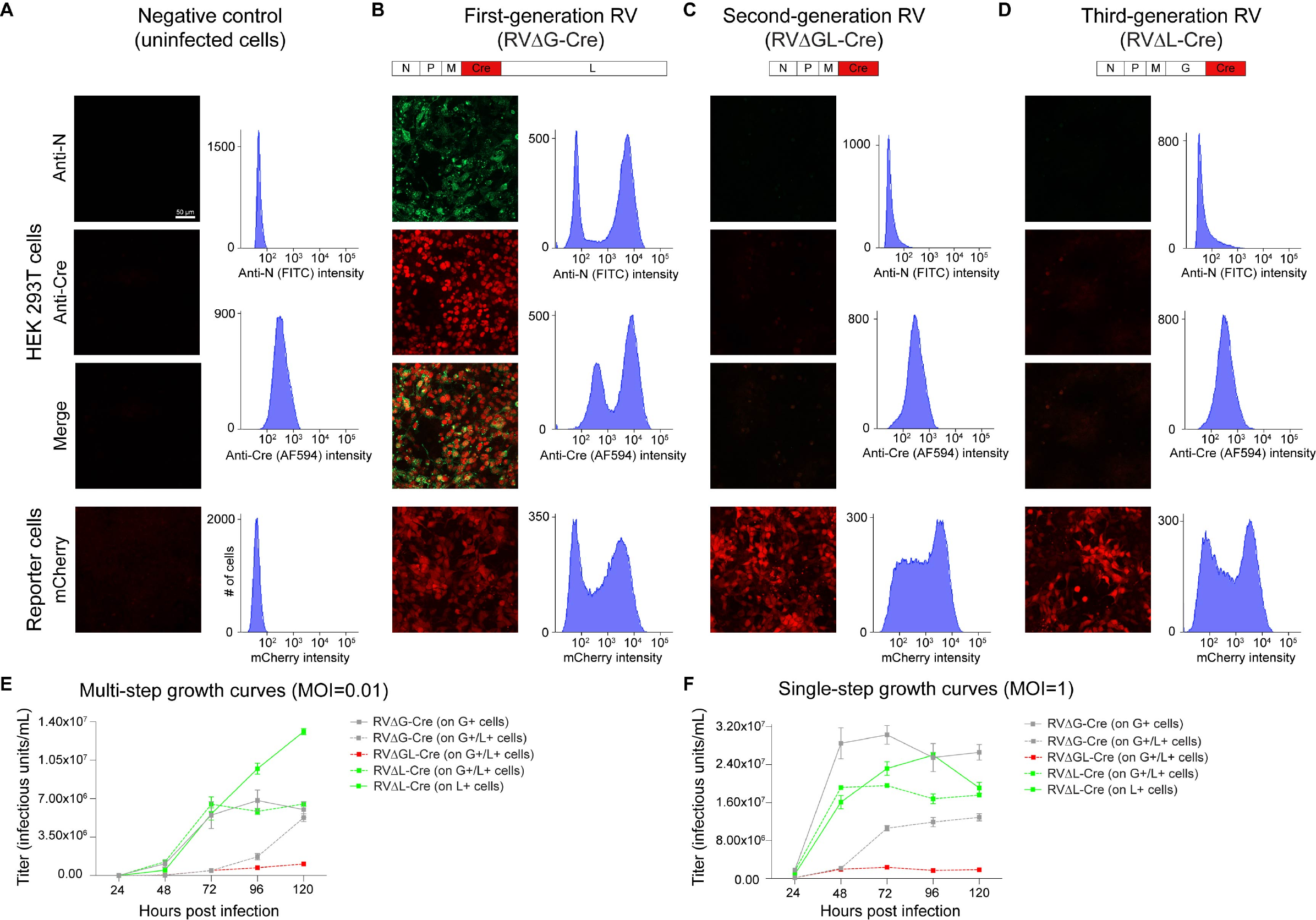
Rabies virus with just the polymerase gene deleted (ΔL) is phenotypically similar to double-deletion-mutant (ΔGL) virus but replicates to much higher titers within complementing cells. (A-D) Deletion of just the polymerase gene L reduces transgene expression to levels that are very low but still sufficient to support reporter allele recombination in Cre reporter cells. (A) Negative controls (uninfected cells). Top: Uninfected HEK 293T cells stained for rabies virus nucleoprotein (green) and for Cre (red). Histograms to right of panels show flow cytometric quantification of baseline fluorescence of uninfected cells in these channels. Bottom: Uninfected reporter cells which express mCherry following Cre recombination. Little signal is seen in these negative controls. (B) Cells infected with a first-generation (ΔG) vector expressing Cre. Both Cre and N are expressed at very high levels, and infected Cre reporter cells brightly express mCherry (note that dilutions at which roughly half of cells were infected were chosen for this figure). (C) Consistent with our previous findings (Chatterjee *et al*., 2018), expression of both nucleoprotein and Cre from a second-generation (ΔGL) vector is drastically reduced with respect to the first-generation vector, with expression levels comparable to those seen in negative controls. Despite this, the low Cre levels are still high enough to activate mCherry expression in reporter cells. (D) A third-generation (ΔL) vector expresses nucleoprotein and Cre at similarly very low levels, but again Cre expression is nonetheless high enough to successfully activate mCherry expression in reporter cells. (E-F) Third-generation (ΔL) vectors grow to much high titers in cultured cells than second-generation (ΔGL) ones do. (E) Viral titers in supernatants of complementing cells (expressing L, G, or both) infected with ΔL, ΔGL, or ΔG viruses at a multiplicity of infection (MOI) of 0.01 (“multi-step growth curves”), with supernatants collected every 24 hours for five days. Whereas a ΔGL virus only achieves 1.05E+06 infectious units (i.u.)/ml over the duration of the experiment, the ΔL virus grows to 6.2-fold higher on the same cell line, and 12.5-fold higher on a line expressing L alone. The highest ΔL titers obtained in this experiment were significantly higher than the highest obtained with a first-generation (ΔG) virus (single-factor ANOVA, p = 3.24E-03, n = 3 replicates per condition). (F) Similarly, at a MOI of 1 (“single-step” growth curves), the ΔGL virus titer peaks at 2.37E+06 i.u./ml, whereas the peak titer of the ΔL virus is 2.60E+07 i.u./ml, 11.0-fold higher than that of the ΔGL virus and not significantly different from that of the ΔG virus (single-factor ANOVA, p = 0.105, n = 3 replicates per condition). Graphs in (E-F) show means ± s.e.m. See Supplementary File S1 for titers and statistical comparisons.

As shown in Fig. 1A-D, whereas the first-generation (ΔG) virus expressed high levels of nucleoprotein (which accumulated in cytoplasmic inclusions) and Cre (which localized to the nuclei), the ΔGL and ΔL viruses had very low expression levels of both Cre and nucleoprotein, with the amount of label for these proteins appearing much more similar to that seen in uninfected control cells than in cells infected with the ΔG virus. We also found similarly low transgene expression levels for ΔL and ΔGL viruses expressing EGFP (Supplementary Fig. S1). However, just as we found previously for ΔGL viruses (Chatterjee *et al*., 2018), the Cre expressed by RVΔL-Cre was sufficient to result in bright labeling of Cre reporter cells (bottom row in panels A-D).

These results led us to predict that ΔL viruses would be as nontoxic as ΔGL ones, because of their similarly low expression levels, and also that they would be similarly able to recombine reporter alleles *in vivo* in order to allow downstream expression of useful transgene products such as fluorophores, activity indicators, or opsins.

It remained to be seen, however, whether ΔL viruses would have any particular advantage over ΔGL ones for purposes of retrogradely targeting neurons. Specifically, if they could not be produced at significantly higher titers, they could be expected to label similar numbers of projection neurons, making ΔL vectors a mere curiosity with no immediate relevance to neuroscientists. However, if they could be grown to much higher titers than ΔGL vectors, that could be expected to translate to the ability to retrogradely label many more projection neurons, a desirable characteristic indeed for a tool for retrograde targeting.

To examine this, we directly compared the ability of ΔL virus to replicate in complementing cells with that of ΔGL and ΔG viruses (Fig. 1E-F). We infected cell lines expressing L, G, or both with the three different generations of virus, at two different multiplicities of infection (MOI, measured in infectious units per cell): either very low (MOI = 0.01, “multi-step growth curves” (Gomme et al., 2010; Wang and Bushman, 2006)) or high (MOI = 1, “single-step growth curves”. Following a one-hour incubation in the presence of the viruses, we washed the cells twice with DPBS and applied fresh medium, then collected supernatant samples every 24 hours for five days after infection, then titered the samples on reporter cells.

As seen in Fig. 1E-F, the results were clear: whereas the ΔGL virus (on cells expressing both G and L) never accumulated to titers higher than 2.37e6 iu/mL in either experiment, the ΔL virus grew to maximal titers of 6.51e6 iu/mL (at MOI of 0.01) and 1.96e7 iu/mL (at MOI of 1) on the same cell line (expressing both G and L) and considerably higher (maximal titers of 1.31e7 iu/mL at MOI=0.01 and 2.60e7 iu/mL at MOI=1) on cells expressing L alone. The ΔG virus grew to similarly high (or slightly higher, in the MOI=1 case) titers to the ΔL one: 6.83e6 iu/mL at MOI=0.01 and 3.03e7 iu/mL at MOI=1, suggesting that single-deletion-mutant rabies viruses may in general be easier to make at high titers than viruses with multiple deleted genes. In non-complementing cells, by contrast, no such replication of any of these viruses (ΔL, ΔGL, or ΔG) occurred (Supplementary Fig. S2).

We furthermore found that ΔL viruses grew to significantly higher titers than comparable ΔGL ones regardless of whether they expressed Cre or Flpo (mouse-codon-optimized Flp recombinase (Raymond and Soriano, 2007)) and regardless of the titering method: see Supplementary Fig. S3. We also found that the titers of ΔGL and ΔL viruses expressing Cre were significantly higher than those of their corresponding Flpo-expressing versions when measured using a method that depends on the activity of the recombinase, presumably due to the higher recombination efficiency of Cre vs Flpo (see Supplementary Fig. S3 for details).

These findings that a ΔL rabies virus could be grown to much higher titers than a corresponding ΔGL one led us to predict that ΔL viruses would be superior tools for retrograde targeting *in vivo*, because their higher titers would result in retrograde infection of many more projection neurons.

### Retrograde targeting *in vivo*

To test this prediction, we made side-by-side preparations (see Methods) of ΔGL and ΔL viruses expressing either Cre or Flpo, then injected each of the four viruses in the somatosensory thalami of reporter mice (Ai14 (Madisen et al., 2010) for the Cre viruses, Ai65F (Daigle et al., 2018) for the Flpo ones; both lines express tdTomato following recombination by the respective recombinase). We sacrificed the mice at either 7 days or 4 weeks after injection, sectioned and imaged the brains by confocal microscopy, and counted the numbers of retrogradely labeled cells in cortex. Fig. 2 shows the results. Note that each data point in the graphs is the total number in one series consisting of every sixth 50 µm section from a given brain – see Methods – so that the total number of labeled S1 neurons in each brain would be approximately six times the corresponding number shown here.

**Figure 2.**
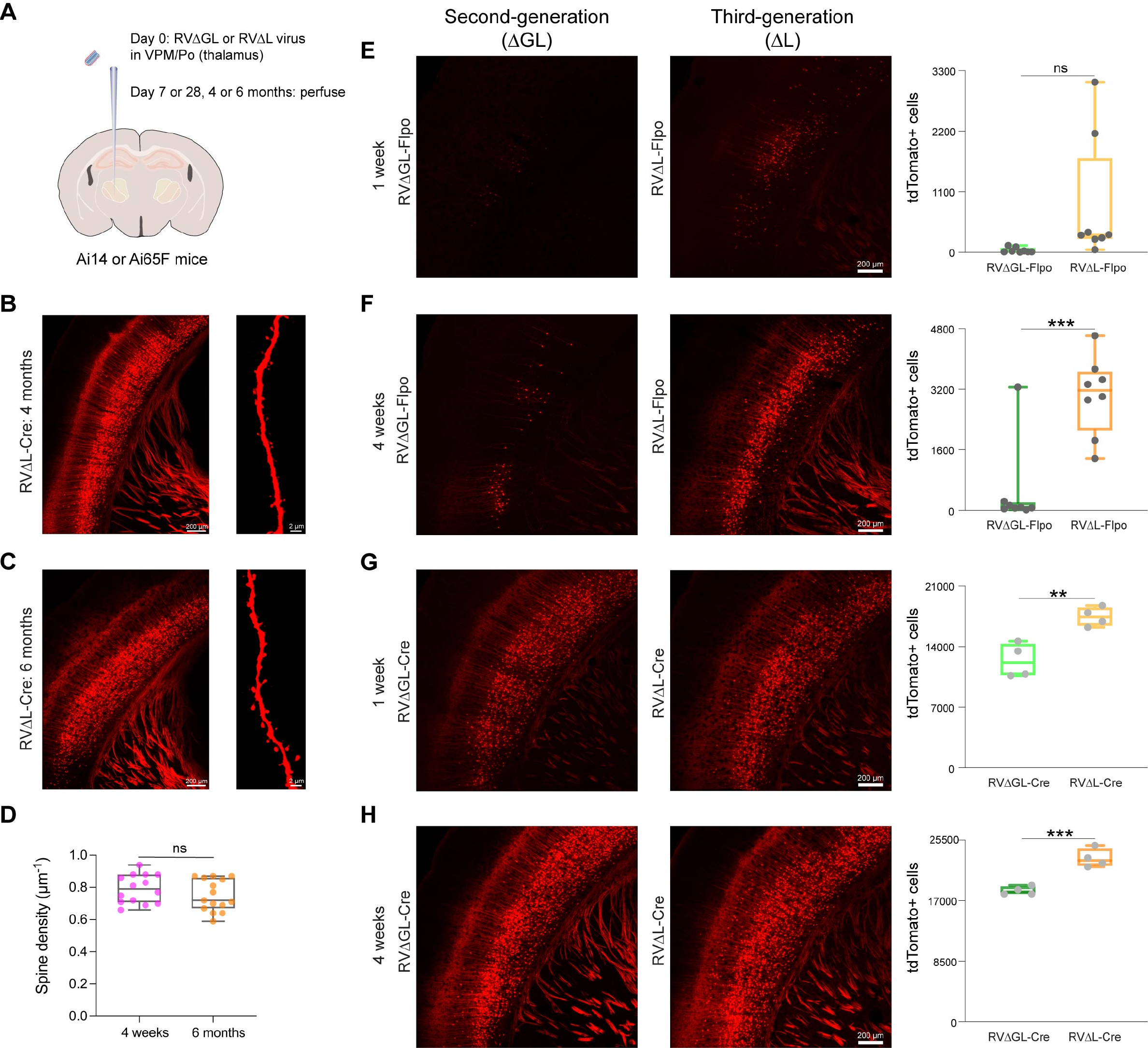
Third-generation (ΔL) rabies viral vectors retrogradely label many more projection neurons *in vivo* than do second-generation (ΔGL) ones and leave cells morphologically normal for at least six months. (A) Design of experiments retrogradely targeting corticothalamic cells in reporter mice. Either second-generation vector RVΔGL-Flpo or RVΔGL-Cre, or third-generation vector RVΔL-Flpo or RVΔL-Cre, was injected into somatosensory thalamus (VPM/Po) of either Ai65F (Flpo reporter) or Ai14 (Cre reporter). Mice were perfused 1 week (E, G), 4 weeks (F, H), 4 months (B), or 6 months later (C). (B) Corticothalamic neurons in S1 of Ai14 mice labeled with RVΔL-Cre at 4 months postinjection. Cells appear morphologically completely normal, with no blebbing or decomposition of processes. Scale bars: 200 µm (left image) and 2 µm (right image). (C) Corticothalamic neurons in S1 of Ai14 mice labeled with RVΔL-Cre at 6 months postinjection. Cells still appear morphologically completely normal. Scale bars: 200 µm (left image) and 2 µm (right image). (D) Quantification of the total number of basal dendritic spines per µm in RVΔL-Cre infected corticothalamic neurons between 4 weeks and 6 months after rabies infection. There was no significant difference between the spine densitites at the two different survival times (single-factor ANOVA, p = 0.26312, 4 weeks: n = 14 FOVs of 2 Ai14 mice; 6 months: n = 15 FOVs of 2 Ai14 mice. See Supplementary File S3 for counts and statistics). (E-H) Efficacy comparison of Flpo- and Cre-expressing ΔGL and ΔL vectors. (E) Corticothalamic neurons in S1 of Ai65F mice labeled with RVΔGL-Flpo (left) or RVΔL-Flpo (center) at 1 week postinjection. Scale bar: 200 µm, applies to both images. Counts of labeled cortical neurons are shown at right (each data point is the total number in one series consisting of every sixth 50 µm section from a given brain – see Methods – so that the total number of labeled S1 neurons in each brain would be approximately six times the corresponding number shown here). The ΔL virus labeled 24 times as many cortical neurons than the ΔGL virus did, although the difference in this case is not significant due to high variance (single-factor ANOVA, p = 0.0608, n = 8 mice per group). (F) Corticothalamic neurons in S1 of Ai65F mice labeled with RVΔGL-Flpo (left) or RVΔL-Flpo (center) at 4 weeks postinjection. Scale bar: 200 µm, applies to both images. Counts of labeled cortical neurons are shown at right. The ΔL virus labeled 6.25 times as many cortical neurons than the ΔGL virus did, an extremely significant difference (single-factor ANOVA, p = 0.000321, n = 8 mice per group). (G) Corticothalamic neurons in S1 of Ai14 mice labeled with RVΔGL-Cre (left) or RVΔL-Cre (center) at 1 week postinjection. Scale bar: 200 µm, applies to both images. Counts of labeled cortical neurons are shown at right. The ΔL virus labeled 1.4 times as many cortical neurons than the ΔGL virus did, a highly significant difference (single-factor ANOVA, p = 0.00420, n = 4 mice per group). (H) Corticothalamic neurons in S1 of Ai14 mice labeled with RVΔGL-Cre (left) or RVΔL-Cre (center) at 4 weeks postinjection. Scale bar: 200 µm, applies to both images. Counts of labeled cortical neurons are shown at right. The ΔL virus labeled 1.25 times as many cortical neurons than the ΔGL virus did, an extremely significant difference (single-factor ANOVA, p = 0.000738, n = 4 mice each group).

As seen in panels 2E-H, for both recombinases, and at both timepoints, the ΔL viruses outperformed the ΔGL ones. For the Flpo viruses, at the 1-week timepoint, the ΔL virus labeled 24 times as many cells as the ΔGL one (although this difference was not statistically significant due to high variance in the ΔL cohort: single factor ANOVA, p=0.275, n = 8 mice each group); by the 4-week timepoint, the ΔL-Flpo virus had labeled 6.25 times as many cells as the ΔGL counterpart, a difference that was significant (single factor ANOVA, p=3.21E-04, n = 8 mice each group). For the Cre viruses, the difference was smaller, presumably due a ceiling effect (see Discussion), but still highly significant: at 1 week, the ΔL-Cre virus had labeled 1.40 times as many cells as ΔGL-Cre (single factor ANOVA, p=4.20E-03; n = 4 mice each group); at the 4-week timepoint, the ΔL-Cre virus had labeled 1.25 times as many cells as ΔGL-Cre (single factor ANOVA, p=7.38E-04, n = 4 mice per group). See Supplementary File S3 for counts and statistical comparisons.

We also made some injections of RVΔL-Cre, in thalami of Ai14 mice, with the much longer survival times of 4 or 6 months (Fig. 2B-C). The results at both of these longer survival times appeared very similar to those at the shorter ones. Consistent with extensive prior literature on corticothalamic neurons (Alitto and Usrey, 2003; Rockland, 2021; Rouiller and Welker, 2000) and with our previous results with corticothalamic injections of ΔG and ΔGL viruses (Chatterjee *et al*., 2018; Wickersham *et al*., 2007a), the cells labeled in cortex by both viruses at all timepoints were pyramidal neurons in layer 6, with a few in layer 5. Furthermore, labeled neurons all appeared morphologically normal even months after injection, with the fine processes of axons and dendrites, including individual spines (rightmost images in 2B-C) clearly visible and without blebbing or other obvious abnormalities. We found no significant difference in spine density between the 4-week and 6-month survival times (Fig 2D; single-factor ANOVA, p = 0.26312, 4 weeks: n = 14 FOVs of 2 Ai14 mice; 6 months: n = 15 FOVs of 2 Ai14 mice. See Supplementary File S4 for counts and statistical comparison).

As a further test of the flexibility of ΔL vectors, we made a version expressing the tetracycline transactivator (tTA) and injected it in the thalamus of Ai63 reporter mice, in which TRE-tight drives tdTomato expression (Daigle *et al*., 2018). As seen in Supplementary Fig. S4, many corticothalamic cells were found retrogradely labeled at both 1-week and 4-week survival times. Unlike with the Cre and Flpo versions, in this case there was no significant difference between the numbers of labeled cells at earlier versus later timepoints (single factor ANOVA, p = 0.772, n = 4 mice per group. See Supplementary File S3 for counts and statistical comparisons); also the numbers of labeled neurons were lower using this virus and reporter mouse line than with the Cre and Flpo versions. These differences could be due either to the transgene *per se* or to the integrative mechanism of readout inherent to a recombinase as opposed to a transactivator (that is, recombination of the CAG-LSL/FSF alleles in the Ai14/Ai65F mouse lines is a binary and permanent event, allowing stochastic accumulation of labeled neurons even with low recombinase expression levels, whereas expression from the TRE-tight-tdTomato allele in the Ai63 mouse line requires ongoing suprathreshold expression of tTA for visibly labeling neurons).

### Comparison of tropism to that of rAAV2-retro and CAV-2

We next compared the efficacy of RVΔL to that of two other types of viral vectors commonly used for retrograde transduction in the nervous system, rAAV2-retro (Tervo et al., 2016) and canine adenoviral vectors (CAV-2) (Soudais et al., 2001) (Fig. 3, Supplementary Fig. S5, and Supplementary Figs. S6 – S11). We ordered samples of these other two vector types from reputable production facilities (rAAV2-retro-hSyn-Cre from Addgene, CAV-Cre from the Plateforme de Vectorologie de Montpellier; see Methods) and injected each of the three viruses undiluted into either the anterior cingulate (ACA) (Fig. 3) or anteromedial (AM) (Supplementary Fig. S5) areas of Ai14 reporter mice, keeping the injectate volume the same (200 nl) for each injection, then perfused the mice at four weeks postinjection.

**Figure 3.**
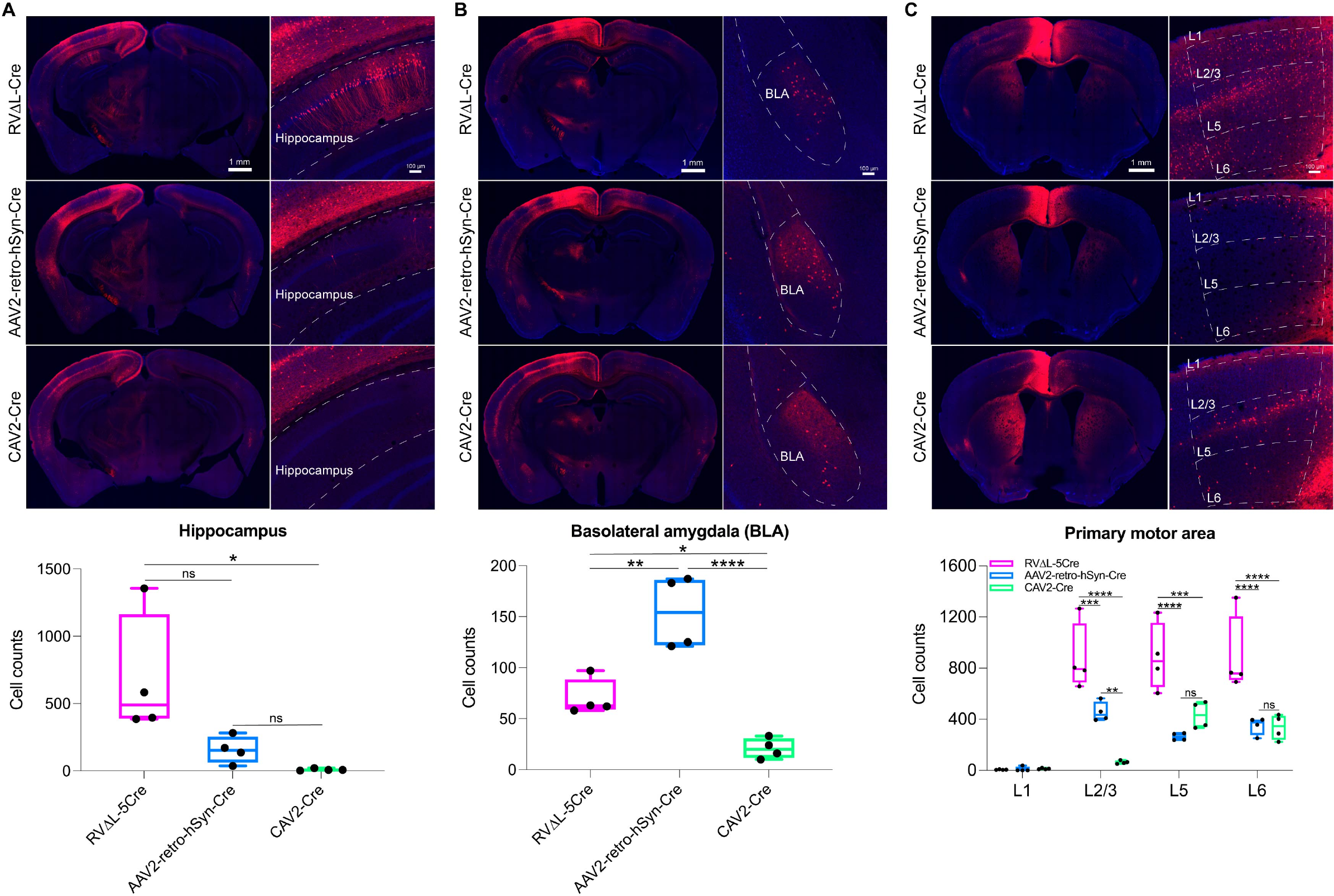
Differential tropism of ΔL rabies virus, rAAV2-retro, and CAV-2. RVΔL-Cre, rAAV2-retro-hSyn-Cre (from Addgene), or CAV-Cre (from the Plateforme de Vectorologie de Montpellier) was injected undiluted into the cortical anterior cingulate area (ACA) of reporter mice, with injections being of equal volumes (200 µl); after a 4-week survival time, brain sections were imaged and labeled neurons in three brain regions were counted. Note that each data point is the total number in one series consisting of every sixth 50 µm section from a given brain (see Methods), so that the total number of labeled neurons in the given region of each brain would be approximately six times the corresponding number shown here. (A) In hippocampus, more neurons were labeled by RVΔL-Cre than by either of the other viruses, although the difference with rAAV2-retro was not quite statistically significant (single factor ANOVA with Tukey’s multiple comparison test, p = 0.05345). CAV-Cre labeled almost no hippocampal cells. (B) In basolateral amygdala, RVΔL labeled more cells than CAV-2 but fewer cells than rAAV2-retro. (C) In ipsilateral primary motor cortex, RVΔL labeled significantly more cells overall, and significantly more cells in individual layers 2/3, 5, and 6, than either of the other two viruses. RVΔL labeled 2.46 times as many cells as rAAV2-retro, and 3.14 times as many cells as CAV-2 (means of 2661, 1080, and 846.25 cells, respectively; again note that each count is of labeled neurons found in a series containing every sixth 50 µm section (see Methods)). See Supplementary Figure S5 for results from similar experiments with the injections in anteromedial visual area; see also Supplementary Figures S6 – S11 for sets of high-resolution confocal images of series of coronal sections from mice labeled with each of the three viruses for each of the two injection sites. See Supplementary File S5 for all counts and statistical comparisons.

We found that the tropism depended on the pathway, to an extent. Following ACA injection (Fig. 3), RVΔL labeled more hippocampal neurons (A) than either rAAV2-retro or CAV-2, although the difference between RVΔL and rAAV2-retro did not quite achieve statistical significance (single factor ANOVA with Tukey’s multiple comparison test, p = 0.05345, n = 4 mice per group; see Supplementary File S5 for counts and statistical comparisons). In basolateral amygdala (B), however, RVΔL labeled more cells than CAV-2 but fewer than rAAV2-retro. In ipsilateral primary motor cortex (C), RVΔL labeled more neurons than either of the other viruses, both overall as well as in individual layers 2/3, 5, and 6 (in layer 1, few neurons were labeled with any of the viruses and differences were not significant). Following AM injection (Supplementary Fig. S5), RVΔL labeled more cells on average in layers 2/3, 5, and 6 in contralateral primary somatosensory cortex than either of the other two viruses, although in each layer the difference between RVΔL and the “runner up” (rAAV2-retro in layers 2/3 and 6, CAV-2 in layer 5) was not statistically significant. Note that many of these comparisons were underpowered due to high variance and low n (= 4 mice per group): for example, for the comparison of ACA-projecting hippocampal neurons labeled by RVΔL vs rAAV2-retro, the sample size needed to achieve 80% power, given the variance we obtained, would have been 13 mice per group (https://www.stat.ubc.ca/~rollin/stats/ssize/n2.html), far more than the 4 mice per group that we used in this study. Few neurons were retrogradely labeled in layers 1 and 4 by any virus, and the differences in those layers were not significant. See Supplementary File S5 for all counts and statistical comparisons.

See also Supplementary Figs. S6 – S11 for sets of high-resolution confocal images of series of coronal sections from mice labeled with each of the three viruses for each of the two injection sites.

### Longitudinal structural two-photon imaging *in vivo*

Because examining only postmortem tissue can be misleading when attempting to determine whether a virus is nontoxic (see (Jin et al., 2023) for a detailed case study), we conducted longitudinal two-photon imaging of RVΔL-labeled neurons *in vivo* (Fig. 4). We injected either RVΔGL-Cre (Chatterjee *et al*., 2018) or RVΔL-Cre in primary visual cortex (V1) of Ai14 reporter mice, then imaged the resulting tdTomato-expressing neurons at or near the injection site beginning 7 days after injection and continuing every 7 or 14 days until 16 weeks postinjection. As seen in Fig. 4, the results using the two viruses were very similar. For both ΔGL and ΔL viruses, the numbers of visibly labeled neurons increased significantly between 1 week and 4 weeks postinjection (Fig. 4D, G), by 56.27% for ΔGL and by 67.77% for ΔL (paired t-tests, p = 1.319E-04 for ΔGL, p = 1.003E-05 for ΔL, n = 8 FOVs for each virus). Also for both viruses, the numbers of visibly labeled neurons remained nearly completely constant from the 4-week timepoint onward through all remaining imaging sessions (Fig. 4E, H) (with the number of labeled cells at the 4-week timepoint being not significantly different than that at the 12-week timepoint for the ΔL virus (paired t-test, p = 0.1327, n = 8 FOVs) and slightly (0.5%) lower for the ΔGL virus (paired t-test, p = 0.0056, n = 8 FOVs). See Supplementary File S6 for all counts and statistical comparisons; see also Supplementary Video S1 for a rendering of a group of ΔL-labeled neurons at 2 weeks and again at 10 weeks.

**Figure 4.**
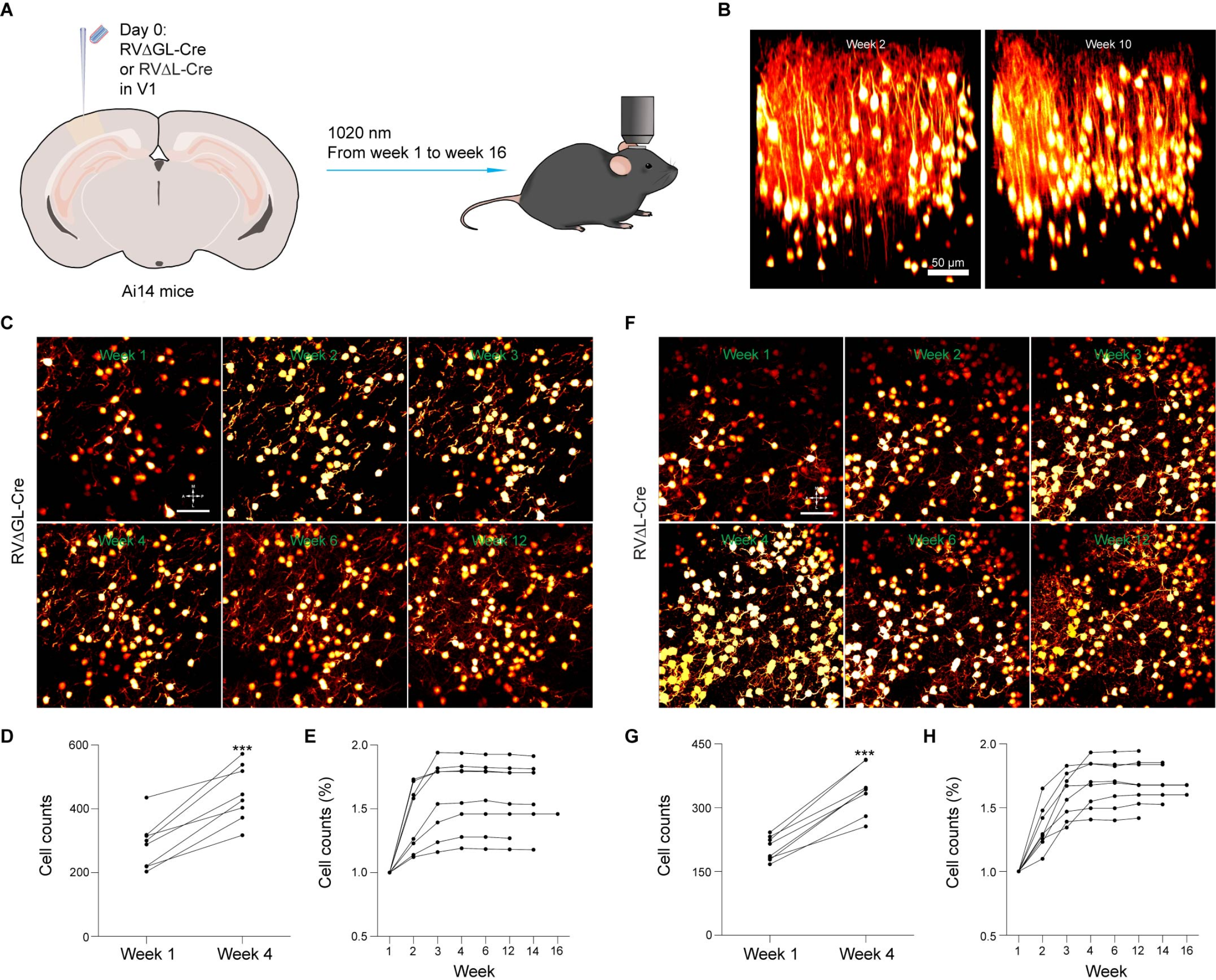
Neurons labeled by ΔL rabies virus survive for at least 16 weeks. (A) Experimental design for longitudinal structural two-photon imaging *in vivo*. Second-generation (ΔGL) or third-generation (ΔL) virus expressing Cre was injected in primary visual cortex of reporter mice, then fields of view near the injection sites were imaged repeatedly for the following 16 weeks (see Methods). (B) Example renderings of the same volume of cortex labeled by RVΔL-Cre and imaged with a two-photon microscope at two different timepoints, 2 weeks (left) and 10 weeks (right). Every labeled neuron visible at 2 weeks is still present at 10 weeks. Scale bar: 50 µm. See also Video S1. (C) & (F), Example two-photon images of single fields of view (FOV) of cortex labeled by either the second-generation vector RVΔGL-Cre (C) or the third-generation vector RVΔL-Cre (F), imaged at different timepoints, from 1 week (top left) to 12 weeks (bottom right). All labeled neurons visible at earlier timepoints are still present at later ones, for both viruses. Scale bars: 50 µm, apply to all images. (D) & (G), Absolute numbers of cells visibly labeled by RVΔGL-Cre (D) or RVΔL-Cre (G) for all structural FOVs in the study, at the 1-week and 4-week timepoints. Numbers of visibly labeled cells increased by 56.27% for ΔGL and by 67.77% for ΔL, as we found previously for second-generation vectors (Chatterjee *et al*., 2018)), suggesting accumulation and persistent activity of recombinase on this timescale. These increases were both extremely significant (one-tailed paired t-tests, p = 0.000132 (ΔGL) and 0.00001003 (ΔL), n = 8 FOVs each virus), but there was no significant difference between the increases seen for the two viruses (two-tailed unpaired t-test, p = 0.5187, n = 8 FOVs per group). Note that the higher numbers of labeled neurons in the ΔGL case is not meaningful and simply reflects the numbers that happened to be present in the fields of view selected quasi-randomly on the basis of sufficient sparsity to allow resolution of individual neurons. (E) & (H), Percentages of cells visibly labeled by RVΔGL-Cre (E) and RVΔL-Cre (H) over time, relative to the numbers visible at 1 week after rabies injection; each connected set of dots represents numbers seen in a given FOV at the different time points. For both viruses, the numbers of labeled neurons remain nearly constant from the 4-week timepoint onward, as we found previously (Chatterjee *et al*., 2018) for ΔGL virus. Imaging was discontinued for some mice at weeks 12 or 14 due to cloudiness of the optical windows.

### *Ex vivo* whole-cell recordings from retrogradely labeled neurons

We conducted an *ex vivo* electrophysiological experiment in order to further assess the health of neurons labeled by ΔL rabies virus (Fig. 5). We injected either RVΔL-Cre or rAAV2-retro-hSyn-Cre into the nucleus accumbens of Ai14 reporter mice, then, at either 4 weeks or 12 weeks postinjection, we made acute brain slices and performed *ex vivo* whole-cell electrophysiological recordings from retrogradely labeled neurons in the basolateral amygdala. We measured 5 membrane properties – resting membrane potential, capacitance, rheobase, input resistance, and action potential number versus current injected – and found no differences between those of the neurons labeled with either virus at either survival time (see Methods). These results suggest that neither RVΔL nor rAAV2-retro perturbed the neurons significantly between the 4-week and 12-week timepoints.

**Figure 5.**
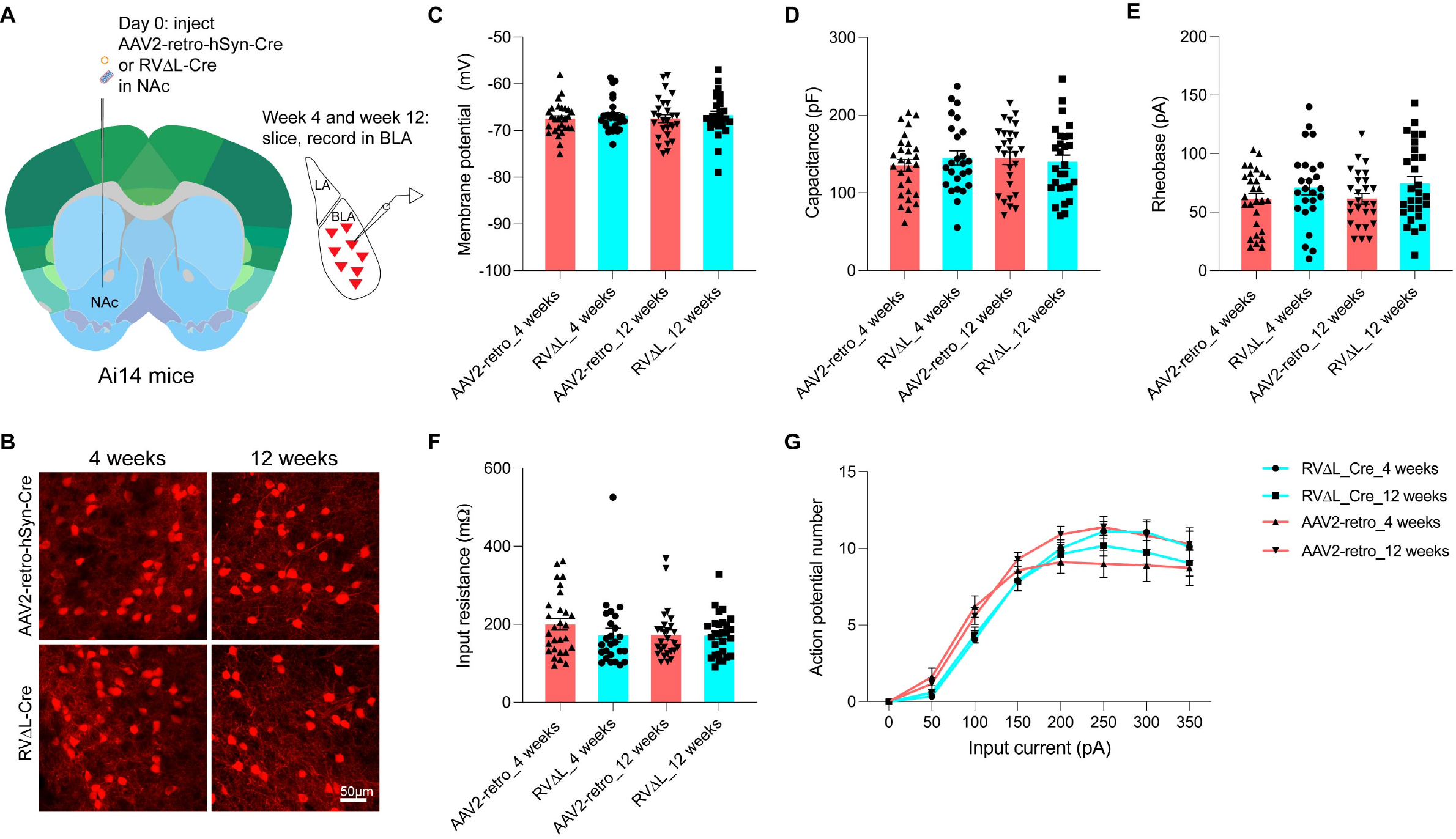
Membrane properties of basolateral amygdalar neurons retrogradely labeled by rAAV2-retro and ΔL rabies virus. (A) Experimental design. rAAV2-retro-hSyn-Cre or RVΔL-5Cre was injected into the nucleus accumbens of reporter mice. 4 weeks or 12 weeks after injection, brain slices were prepared for whole cell patch-clamp recordings from labeled cells in basolateral amygdala (BLA). (B) Confocal images of BLA neurons labeled by the two viruses following the two survival times. (C-G) None of the four groups (two viruses, two survival times) differed significantly from the others in any of the measured membrane properties: resting membrane potential (C), membrane capacitance (D), rheobase (E), input resistance (F), or action potential number versus input current (G). See Methods - Quantification and Statistical Analysis for details of comparisons.

### Longitudinal functional two-photon imaging *in vivo*

We went on to examine the functional properties of RVΔL-labeled neurons *in vivo*. As for the structural imaging (see above), we injected RVΔL-Cre into the primary visual cortices of reporter mice, in this case mice that express the calcium indicator GCaMP6s (Chen et al., 2013) after Cre recombination (Fig. 6). Beginning one week later, we imaged the calcium signals in the labeled neurons as the awake mice viewed visual stimuli consisting of drifting gratings of different orientations and frequencies, in a series of imaging sessions that continued until 16 weeks postinjection. Just as we found previously for ΔGL viruses (Chatterjee *et al*., 2018; Jin *et al*., 2021), we found no signs of dysfunction in cells labeled by the third-generation vector even at 16 weeks postinjection, the longest we followed them (Fig. 6; see also Supplementary Figs. S12 and S13. See Supplementary File S7 for all counts and statistical comparisons and Supplementary Video S2 for an example of calcium responses in a group of cortical neurons 16 weeks after injection of RVΔL-Cre). Unlike for the structural imaging (Fig. 4), for this functional imaging we were not in general able to track individual neurons over multiple imaging sessions, because we imaged GCaMP6s (rather than the clearer tdTomato) signal only in a single plane in each field of view (rather than z-stacks as for the structural imaging), and in awake mice in which motion was more of a factor (whereas for the structural imaging the mice were anesthetized). Nevertheless, in some cases we were able to track individual identified neurons across imaging sessions (Fig. 6C-D and Supplementary Fig. S13). Fig. 6G shows the numbers of cells that were tuned at 4 weeks that were identifiable at 14 weeks: 94% of these neurons were still tuned at 14 weeks.

**Figure 6.**
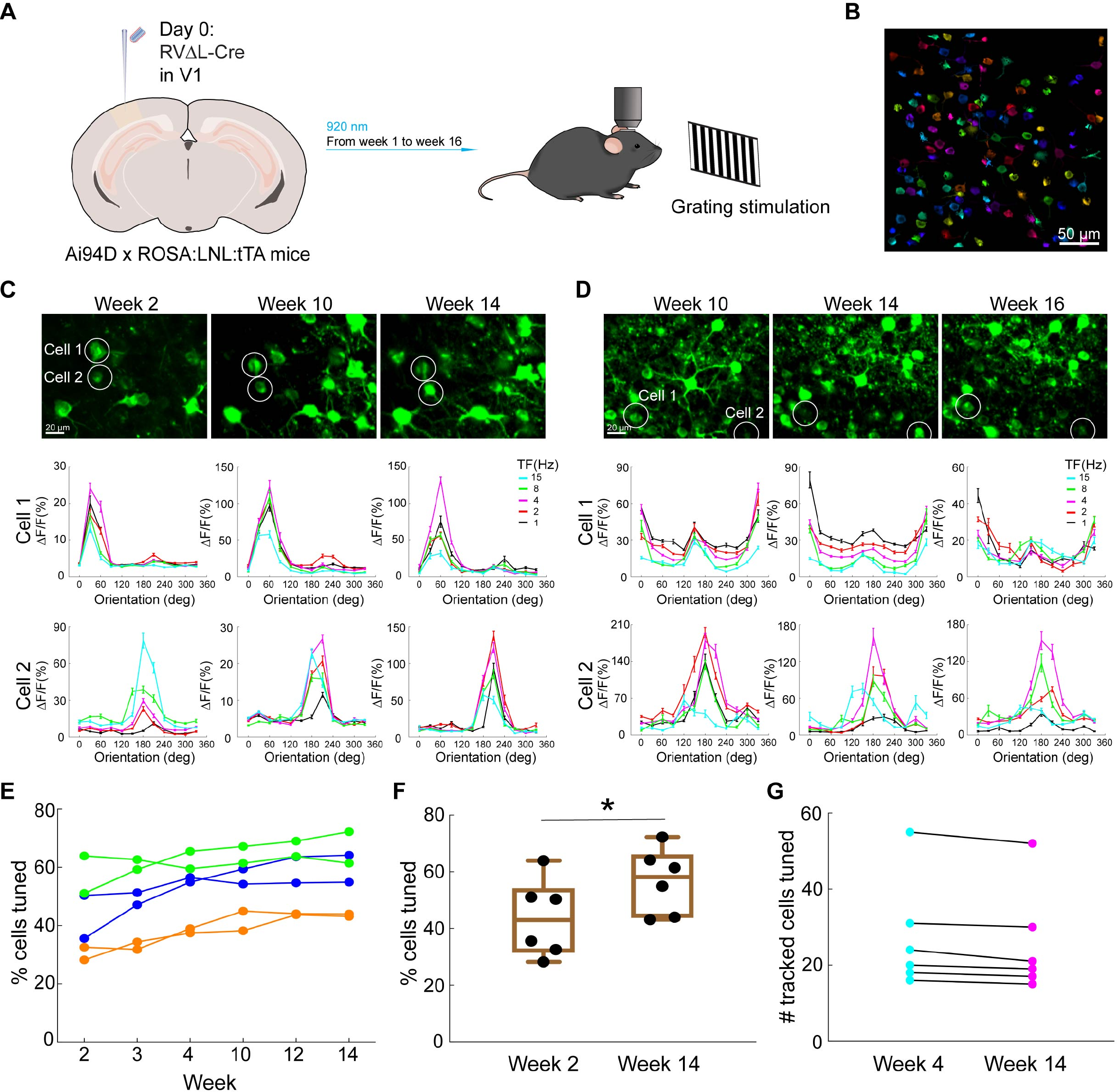
Imaging of ΔL-labeled neurons’ visual response properties over 16 weeks. (A) Experimental design for longitudinal functional two-photon imaging *in vivo*. ΔL virus expressing Cre was injected in primary visual cortex of reporter mice expressing GCaMP6s (Chen *et al*., 2013) after Cre recombination, then the injection sites were imaged while the awake mice were presented with drifting grating stimuli of different orientations and temporal frequencies, repeatedly for 16 weeks following virus injection. (B) Example FOV from a GCaMP6s imaging session 16 weeks after RV injection. Individual analyzed cells are randomly pseudocolored. This is the same FOV as shown in Supplementary Video S2. Scale bar: 50 μm. (C-D) Long-term stability of orientation and temporal frequency tuning in RVΔL-Cre-labeled neurons. The top rows show maximum intensity projections of the imaged GCaMP6s signal in two different FOVs at three different timepoints for each FOV. Scale bars: 20 µm, apply to all images. Visual response tuning curves of the two circled cells in each FOV at the corresponding timepoint, obtained with drifting gratings presented at 12 directions of motion and 5 temporal frequencies (TF) (mean ΔF/F ± s.e.m., averaged over 10 repeats), are shown under each image. More examples from the same FOV are shown in Supplementary Figure S13. (E) Percentages of labeled cells that were visually tuned (see Methods), from 6 different FOVs in 3 mice imaged over 14 weeks. Connected sets of dots in a given color indicate data from a single mouse (data from 2 FOVs are shown per mouse). (F) Comparison of the percentages of labeled cells that were visually tuned at 2 weeks and 14 weeks. The percentages increased moderately but significantly between the two timepoints, from 60% to 68% (paired two-sample t-test, p = 0.0178, n = 6). (G) Although in general we were not able to track individual neurons over multiple imaging sessions due to limitations of our imaging capabilities, in some cases we were able to do so. This graph shows the numbers of cells that were tuned at 4 weeks that were identifiable at 14 weeks and that were still tuned at that later timepoint: 94% of these neurons were still tuned at 14 weeks.

## DISCUSSION

Here we have shown that deletion of only the polymerase gene renders rabies viral vectors nontoxic, like second-generation (ΔGL) vectors, but leaves them able to replicate much more efficiently within complementing cells. This ability to be grown to much higher titers results in significantly increased transduction of projection neurons within a given pathway. This more comprehensive access to projection neurons will increase the yield and efficacy of systems neuroscience experiments that depend on the retrograde targeting approach.

In the corticothalamic pathway that we have examined here, the advantage of a ΔL vector over the ΔGL equivalent was clearest in the case of the Flpo-expressing versions, with the ΔL vector labeling 6.25 times as many neurons as the ΔGL one did at four weeks postinjection. This ratio is of the same order of magnitude as the ratio of the titers of the injected Flpo viruses (14.4; see Methods). By contrast, for the Cre-expressing versions, the advantage of the ΔL vector over the ΔGL one was more modest, labeling 1.25 times as many cells, even though the ratio of the titers of these Cre vectors was even higher (20.5). Because the absolute numbers of retrogradely labeled neurons, as well as the titers, were much higher for the Cre viruses than for the corresponding Flpo ones, we presume that the smaller advantage of the ΔL version seen in this case was because of a ceiling effect, with the ΔGL-Cre virus already labeling most of the available projection neurons in this pathway, which may itself be due to a higher efficacy of Cre versus Flpo (see Supplementary Fig. S3 for data supporting this interpretation). Although we only quantified the efficiency of ΔL vs ΔGL labeling in the densely-projecting somatosensory corticothalamic pathway, in which the advantage of ΔL over ΔGL was modest (25% higher) in the case of the Cre-expressing vectors, it seems likely that the much higher titer of ΔL-Cre vs ΔGL-Cre (11-fold higher in the growth curve experiments of Fig. 1, 20-fold higher in the concentrated stocks used for Fig. 2) would result in a more dramatic advantage in other pathways that are labeled less thoroughly (perhaps because of sparser projections) by ΔGL-Cre. We also note that, although most users are likely to require the Cre-expressing version, the major advantage of ΔL that we observed in the case of the Flpo-expressing vectors could be very helpful for many applications in which Cre is needed for some other purpose or otherwise in which two recombinases are required (such as intersectional targeting, retrograde targeting in mice with floxed genomic alleles, or independently labeling two different neuronal populations).

One could argue that the much higher titers that we are easily able to obtain with ΔL vectors could also, in theory, potentially be achieved with ΔGL vectors, if enough effort were put into, for example, generating and testing producer cell lines expressing both G and L in order to find one that expressed the two genes at just the right ratio and levels. In practice, however, this hypothetical future research effort does not detract from the fact that the best currently-existing preparations of ΔL rabies viral vectors label many more cells than do ΔGL ones, making them the better choice for retrograde targeting applications.

We note that, although here we have only demonstrated the use of ΔL rabies viral vectors in mice, they are also highly likely to work in a wide variety of mammalian species, because, apart from their shorter RNA genomes, the structural properties of second- and third-generation rabies viral particles are identical to those of first-generation ones, which have been successfully used in diverse mammalian species including rats (Cruz et al., 2021), cats (Connolly et al., 2012; Liu et al., 2013), ferrets (Hasse et al., 2019), and macaques (Bragg et al., 2017; Briggs et al., 2016; Lyon et al., 2010; Nassi and Callaway, 2006; 2007; Nassi et al., 2006; Siu *et al*., 2021; Yarch et al., 2017) (and even in frogs (Faulkner et al., 2021) and fish (Dohaku et al., 2019; Satou et al., 2021; Zhu et al., 2009)).

Our findings here that ΔL rabies viruses have extremely low expression levels and do not replicate within (or spread beyond, *in vivo*) non-complementing cells are entirely consistent with similar findings in cell culture in a recent report on an L-deficient rabies virus encoding firefly luciferase (Nakagawa et al., 2017).

A note about safety: our results strongly suggest that ΔL rabies viruses are unable to replicate in the absence of complementation and moreover are harmless to any cells that they transduce. However, a mixture of ΔL and ΔG viruses could pose a safety risk, because such viruses will be mutually complementary. Care must therefore be taken to avoid contamination between ΔL and ΔG constructs – either packaged viruses or the genome plasmids used to make them – which would have the potential to create a self-complementing replication-competent mixture (see Hidaka et al. (Hidaka et al., 2018) for an example of such a self-complementing mixture).

Finally, we have recently shown (Jin *et al*., 2021) that second-generation (ΔGL) rabies viral vectors can spread transsynaptically when complemented by provision of both G and L *in trans*. That is, complementation of an L-deficient rabies virus (in that case, a G- and L-deficient virus that is also complemented by G) allows it to spread beyond initially infected cells *in vivo*. It is therefore reasonable to infer that provision of L *in trans* should allow third-generation, ΔL rabies viral vectors to spread beyond initially infected cells, especially given that we have shown here that such complementation in cell culture allows ΔL viruses to replicate very efficiently. We have also shown here, with the longitudinal two-photon imaging of labeled neurons, that ΔL viruses do not spread beyond initially infected cells *in vivo* in the absence of complementation. Collectively, our results therefore suggest the outlines of a third-generation monosynaptic tracing system based on ΔL vectors complemented with L expression *in trans*. However, genetic targeting of a ΔL vector to specific starting cell types might appear elusive: in the first- and second-generation systems (Jin *et al*., 2021; Wickersham *et al*., 2007b), this targeting is achieved by packaging the rabies viral particles with an avian retroviral envelope protein (EnvA) instead of its own envelope glycoprotein, so that they can only infect cells that have been engineered to express EnvA’s cognate receptor. On the face of it, this pseudotyping strategy requires that G be deleted from the rabies viral genome, because expression of G by the virus within the EnvA-expressing producer cells would result in the production of virions with membranes populated by a mixture of EnvA and the rabies viral glycoprotein. If this challenge could be overcome, our present findings that ΔL viruses are more efficient at replication in complementing cells, which is the fundamental process central to monosynaptic tracing (Wickersham *et al*., 2007b), suggest that a third-generation monosynaptic tracing system could be more efficient than the second-generation one.

## SUPPLEMENTAL INFORMATION

Supplemental information can be found online.

A preprint version of this paper is available on bioRxiv.

## ACKNOWLEDGMENTS

We thank Tanya Daigle and Hongkui Zeng for sharing the Ai63 mouse line, Jacque Ip, Chloe Delepine, and Mriganka Sur for sharing mice, Chang Liu for assistance with optimizing MATLAB code for analysis of the functional imaging data, and Sara Beach for helpful suggestions on the manuscript. We acknowledge E.J. Kremer as the source of CAV-Cre, via Plateforme de Vectorologie de Montpellier. Research reported in this publication was supported by BRAIN Initiative awards RF1MH120017, U01MH106018, U01MH114829, and U19MH114830 from the National Institute of Mental Health.

## AUTHOR CONTRIBUTIONS

L.J., N.E.L., M.M., Y.H., and M.Z. cloned constructs; H.A.S. produced viruses with assistance from L.J. and M.Z.; H.A.S. made cell lines and conducted cell culture assays and immunocytochemistry; L.J., N.E.L., and T.K.L. performed surgeries; L.J. and M.Z. performed histology and confocal imaging; L.J. quantified spine densities and performed two-photon imaging; L.J., M.Z., T.K.L., and N.E.L. managed mouse breeding; X.F and G.F. planned slice electrophysiology experiments; X.F. conducted slice electrophysiology experiments; I.W. planned and supervised all work; I.W. and L.J. wrote the manuscript with input from the other authors.

## STAR METHODS

### KEY RESOURCES TABLE

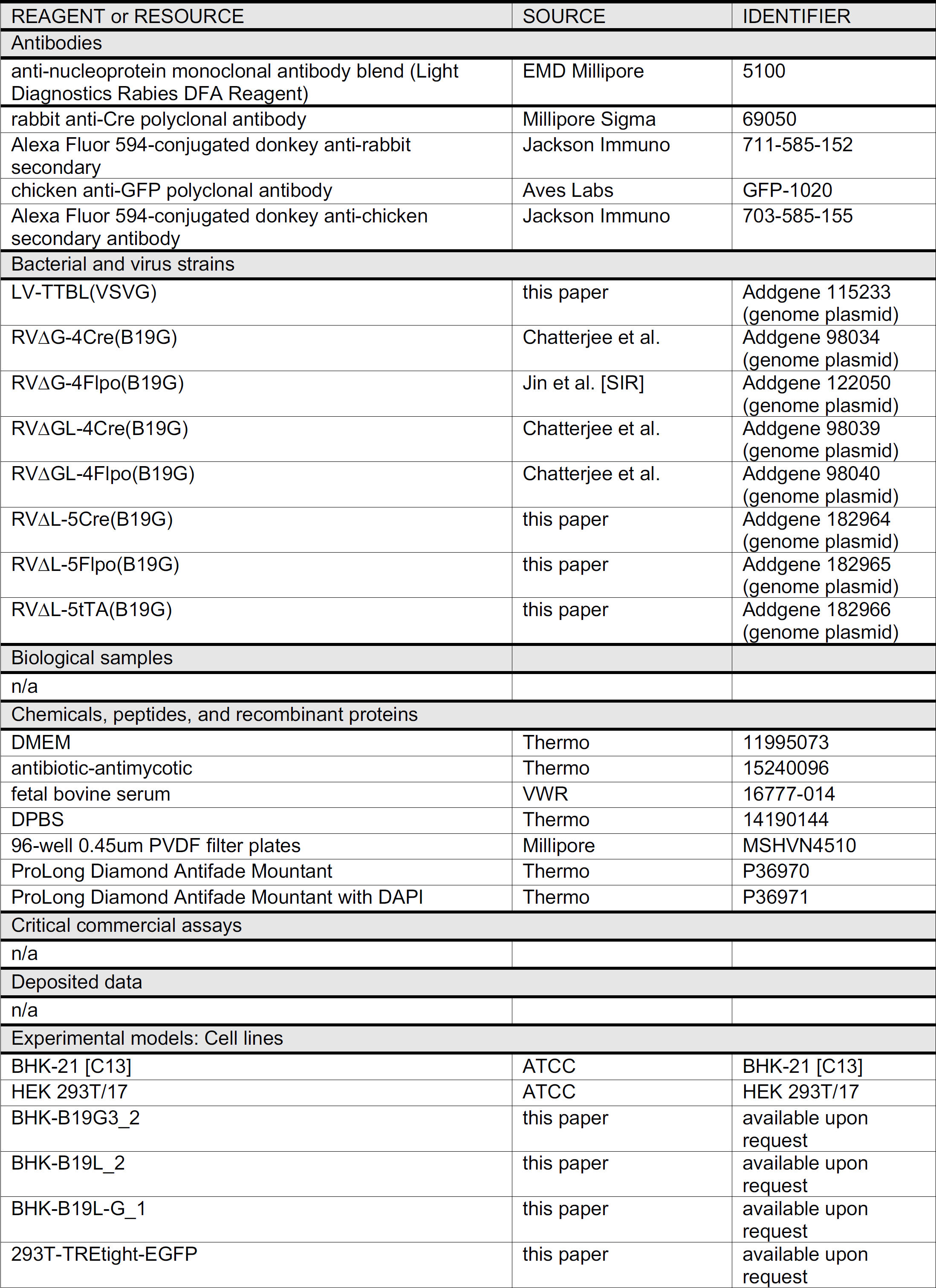
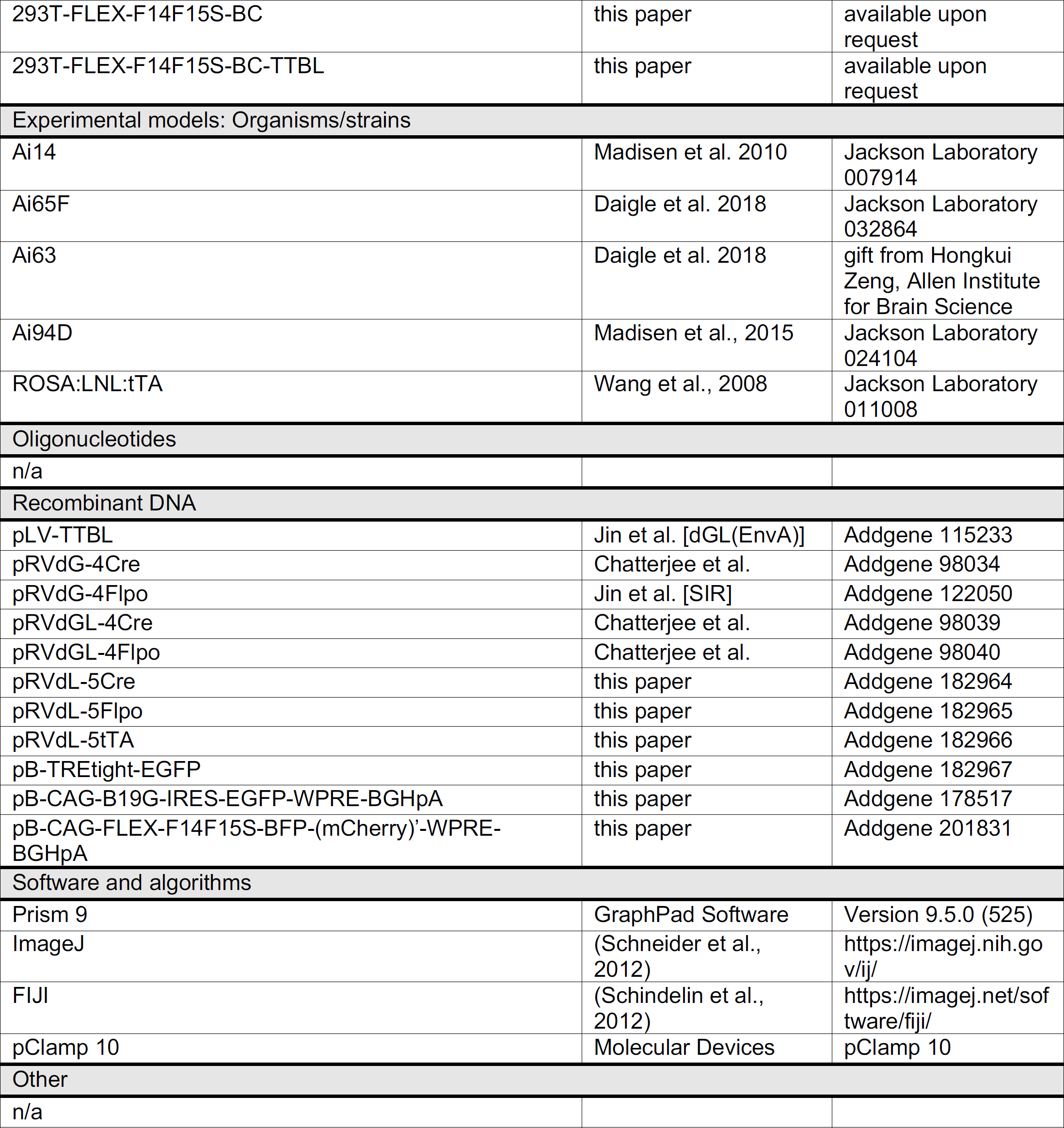

| REAGENT or RESOURCE | SOURCE | IDENTIFIER |
| --- | --- | --- |
| Antibodies |  |  |
| anti-nucleoprotein monoclonal antibody blend (Light Diagnostics Rabies DFA Reagent) | EMD Millipore | 5100 |
| rabbit anti-Cre polyclonal antibody | Millipore Sigma | 69050 |
| Alexa Fluor 594-conjugated donkey anti-rabbit secondary | Jackson Immuno | 711-585-152 |
| chicken anti-GFP polyclonal antibody | Aves Labs | GFP-1020 |
| Alexa Fluor 594-conjugated donkey anti-chicken secondary antibody | Jackson Immuno | 703-585-155 |
| Bacterial and virus strains |  |  |
| LV-TTBL(VSVG) | this paper | Addgene 115233 (genome plasmid) |
| RVΔG-4Cre(B19G) | Chatterjee et al. | Addgene 98034 (genome plasmid) |
| RVΔG-4Flpo(B19G) | Jin et al. [SIR] | Addgene 122050 (genome plasmid) |
| RVΔGL-4Cre(B19G) | Chatterjee et al. | Addgene 98039 (genome plasmid) |
| RVΔGL-4Flpo(B19G) | Chatterjee et al. | Addgene 98040 (genome plasmid) |
| RVΔL-5Cre(B19G) | this paper | Addgene 182964 (genome plasmid) |
| RVΔL-5Flpo(B19G) | this paper | Addgene 182965 (genome plasmid) |
| RVΔL-5tTA(B19G) | this paper | Addgene 182966 (genome plasmid) |
| Biological samples |  |  |
| n/a |  |  |
| Chemicals, peptides, and recombinant proteins |  |  |
| DMEM | Thermo | 11995073 |
| antibiotic-antimycotic | Thermo | 15240096 |
| fetal bovine serum | VWR | 16777-014 |
| DPBS | Thermo | 14190144 |
| 96-well 0.45um PVDF filter plates | Millipore | MSHVN4510 |
| ProLong Diamond Antifade Mountant | Thermo | P36970 |
| ProLong Diamond Antifade Mountant with DAPI | Thermo | P36971 |
| Critical commercial assays |  |  |
| n/a |  |  |
| Deposited data |  |  |
| n/a |  |  |
| Experimental models: Cell lines |  |  |
| BHK-21 [C13] | ATCC | BHK-21 [C13] |
| HEK 293T/17 | ATCC | HEK 293T/17 |
| BHK-B19G3_2 | this paper | available upon request |
| BHK-B19L_2 | this paper | available upon request |
| BHK-B19L-G_1 | this paper | available upon request |
| 293T-TREtight-EGFP | this paper | available upon request |
| 293T-FLEX-F14F15S-BC | this paper | available upon request |
| 293T-FLEX-F14F15S-BC-TTBL | this paper | available upon request |
| Experimental models: Organisms/strains |  |  |
| Ai14 | Madisen et al. 2010 | Jackson Laboratory 007914 |
| Ai65F | Daigle et al. 2018 | Jackson Laboratory 032864 |
| Ai63 | Daigle et al. 2018 | gift from Hongkui Zeng, Allen Institute for Brain Science |
| Ai94D | Madisen et al., 2015 | Jackson Laboratory 024104 |
| ROSA:LNL:tTA | Wang et al., 2008 | Jackson Laboratory 011008 |
| Oligonucleotides |  |  |
| n/a |  |  |
| Recombinant DNA |  |  |
| pLV-TTBL | Jin et al. [dGL(EnvA)] | Addgene 115233 |
| pRVdG-4Cre | Chatterjee et al. | Addgene 98034 |
| pRVdG-4Flpo | Jin et al. [SIR] | Addgene 122050 |
| pRVdGL-4Cre | Chatterjee et al. | Addgene 98039 |
| pRVdGL-4Flpo | Chatterjee et al. | Addgene 98040 |
| pRVdL-5Cre | this paper | Addgene 182964 |
| pRVdL-5Flpo | this paper | Addgene 182965 |
| pRVdL-5tTA | this paper | Addgene 182966 |
| pB-TREtight-EGFP | this paper | Addgene 182967 |
| pB-CAG-B19G-IRES-EGFP-WPRE-BGHpA | this paper | Addgene 178517 |
| pB-CAG-FLEX-F14F15S-BFP-(mCherry)'-WPRE-BGHpA | this paper | Addgene 201831 |
| Software and algorithms |  |  |
| Prism 9 | GraphPad Software | Version 9.5.0 (525) |
| ImageJ | (Schneider et al., 2012) | https://imagej.nih.gov/ij/ |
| FIJI | (Schindelin et al., 2012) | https://imagej.net/software/fiji/ |
| pClamp 10 | Molecular Devices | pClamp 10 |
| Other |  |  |
| n/a |  |  |

## RESOURCE AVAILABILITY

### Lead contact

Requests for resources and reagents should be directed to the lead contact, Ian Wickersham.

### Materials availability

The novel plasmids described in this paper have all been deposited with Addgene with the accession numbers given below. The novel cell lines are available from the lead contact upon request.

### Data and code availability

- All cell and spine counts and statistical analyses are provided in Supplemental Information. All other data reported in this paper will be shared by the lead contact upon request.
- This paper does not report original code.
- Any additional information required to reanalyze the data reported in this paper is available from the lead contact upon request.

## EXPERIMENTAL MODEL AND STUDY PARTICIPANT DETAILS

All experiments involving animals were conducted according to NIH guidelines and approved by the MIT Committee for Animal Care.

### Mouse strains

The Cre-dependent tdTomato reporter line Ai14 (Madisen *et al*., 2010) was purchased from Jackson Laboratory (catalog # 007914). The Flp-dependent tdTomato reporter line Ai65F was obtained by crossing the Cre- and Flp-dependent tdTomato double-reporter line Ai65D (Madisen et al., 2015) (Jackson Laboratory 021875) to the Cre deleter line Meox2-Cre (Tallquist and Soriano, 2000) (Jackson Laboratory 003755), then breeding out the Meox2-Cre allele. An equivalent Ai65F line, made using a different Cre deleter line, was described in Daigle et al. ’18 (Daigle *et al*., 2018) and is now available from Jackson Laboratory (catalog # 032864). The tTA-dependent tdTomato reporter line Ai63 (Daigle *et al*., 2018) was a generous gift from Hongkui Zeng and Tanya Daigle. Mice used for the functional two-photon imaging experiments were crosses of the Cre- and tTA-dependent GCaMP6s line Ai94D (Madisen *et al*., 2015) (Jackson Laboratory 024104) with the Cre-dependent tTA line ROSA:LNL:tTA (Wang et al., 2008) (Jackson Laboratory 011008). All mice were maintained in a C57BL/6J (Jackson Laboratory 000664) background.

For experiments, adult mice (>6 weeks old) of both sexes were used. We did not analyze the influence of sex on the results of the study due to the lack of sexual dimorphism in the projections examined; some neural pathways, particularly in regions involved in sexually dimorphic behaviors (Kim et al., 2019) could presumably exhibit sex differences in the degree of retrograde labeling by viral vectors.

Mice were housed 1-5 per cage under a normal light/dark cycle for all experiments. Strains used were as follows. For retrograde targeting using Cre-expressing viruses (Fig. 2) and structural two-photon imaging (Fig. 4): Ai14 heterozygotes. For retrograde targeting using Flpo-expressing viruses (Fig. 2): Ai65F heterozygotes. For retrograde targeting using RVΔL-tTA (Supplementary Figure S4): Ai63 heterozygotes. For functional two-photon imaging (Figs. 6, S12, and S13): Ai94D x ROSA:LNL:tTA double homozygotes.

Information on all individual mice used in this study (ID #, strain, genotype, age, weight, health status, whether involved in previous procedures, housing) is provided in Supplementary File S8.

### Cell line sourcing and culture

Cell lines used for virus production and analysis were obtained from ATCC (BHK-21 [C13] cells, cat. # CCL-10, unsexed); HEK 293T/17, cat. # 11268, female) without further analysis or validation and either used directly or modified as described below. Cells were cultured in standard cell culture incubators at 37°C and 5% CO_2_ as described (Wickersham and Sullivan, 2015).

## METHOD DETAILS

### Cloning

The third-generation rabies viral vector genome plasmids pRVΔL-5Cre, pRVΔL-5Flpo, and pRVΔL-5tTA (Addgene 182964, 182965, and 182966) (the “5” denoting the position of the transgene relative to the other genes in the viral genome) was made by replacing the mCre gene in pRVΔGL-4Cre (Chatterjee *et al*., 2018) (Addgene 98039) with the SAD B19 glycoprotein gene from pCAG-B19G (Chatterjee *et al*., 2018) (Addgene 59921) and either the mCre, Flpo (from pRVΔG-4Flpo (Addgene 98040)), or tTA (from pAAV-syn-FLEX-splitTVA-EGFP-tTA (Liu et al., 2017) (Addgene 100798)) gene, separated by endogenous rabies viral transcriptional stop and start signals, using seamless cloning (InFusion (Takara) or HiFi (NEB)).

The piggyBac plasmid pB-TREtight-EGFP (Addgene 182967) was made by cloning the TRE-tight element from pAAV-TREtight-mTagBFP2-B19G (Liu *et al*., 2017) and the EGFP gene into pB-CAG-TEVp-IRES-mCherry (Addgene 174377) in place of the CAG-TEVp-IRES-mCherry sequences using HiFi seamless cloning (NEB).

The piggyBac plasmid pB-CAG-B19G-IRES-EGFP-WPRE-BGHpA (Addgene 178517) was made by cloning the CAG promoter from pCAG-B19G (Addgene 59921), the SAD B19 L gene, the EMCV IRES (Gallardo et al., 1997), the EGFP gene, and the woodchuck post-transcriptional regulatory element and bovine growth hormone polyadenylation signal from pCSC-SP-PW-GFP (Addgene 12337), into PB-CMV-MCS-EF1-Puro (System Biosciences #PB510B-1).

The piggyBac plasmid pB-CAG-FLEX-F14F15S-BFP-(mCherry)’-WPRE-BGHpA (Addgene 201831) was made by cloning the CAG promoter from pCAG-B19G (Addgene 59921), the mTagBFP2 (Subach et al., 2011) and reverse-complemented mCherry (Shaner et al., 2004) genes flanked by orthogonal lox and FRT sites, and the woodchuck post-transcriptional regulatory element and bovine growth hormone polyadenylation signal from pCSC-SP-PW-GFP (Addgene 12337), into PB-CMV-MCS-EF1-Puro (System Biosciences #PB510B-1).

### Lentivirus production and titering

The lentiviral vector LV-TTBL(VSVG) was made as described (Wickersham et al., 2015) using the genome plasmid pLV-TTBL (Jin *et al*., 2021) (Addgene 115233) and using the VSV envelope expression plasmid pMD2.G (Addgene 12259) instead of pCAG-B19GVSVGCD, then purified and titered as described in (Wickersham et al., 2010).

### Cell line production

The BHK-B19G3 cell line, expressing the SAD B19 strain rabies virus glycoprotein gene, was made by resorting BHK-B19G2 cells (Wickersham *et al*., 2010) on a BD Facs Aria cell sorter and retaining the brightest 2% of EGFP-positive cells as well as the next-brightest 18%. Following the sort, both populations were expanded and refrozen, then thawed and tested for their efficacy at supporting replication of ΔG virus; the second-brightest population (“BHK-B19G3_2”) was found to result in higher titers and is referred to here as BHK-B19G3.

The BHK-B19L cell line, expressing the SAD B19 strain rabies virus polymerase gene, was made by transfecting BHK-21 cells (ATCC CCL-10) with pCAG-hypBase (Jin *et al*., 2021) and pB-CAG-B19L-IRES-mCherry-WPRE-BGHpA (Jin *et al*., 2021) using Lipofectamine 2000 (Thermo Fisher 11668019), then expanding the cells and sorting on a FACS Aria sorter (BD) to collect the brightest 5%, as well as the next brightest 5%, of mCherry-expressing cells. The two collected populations were expanded and refrozen, then thawed and tested for their efficacy at supporting replication of ΔL virus; the second-brightest population (“BHK-B19L_2”) was found to result in higher titers and is referred to here as BHK-B19L.

The BHK-B19L-G cell line, expressing the SAD B19 strain rabies virus polymerase and glycoprotein genes, was made by transfecting BHK-B19L cells (see above) with pCAG-hypBase and pB-CAG-B19G-IRES-EGFP-WPRE-BGHpA (see above), then expanding and sorting on a BD FACS Aria, keeping the brightest 5%, as well as the next brightest 5%, of EGFP-expressing cells which also expressed mCherry. The sorted cells were expanded and refrozen, then thawed and tested for their efficacy at supporting replication of ΔGL virus; the brightest population (“BHK-B19L-G_1”) was found to result in higher titers and is referred to here as BHK-B19L-G.

The 293T-TREtight-EGFP cell line for titering tTA-expressing viruses was made by transfecting HEK 293T/17 cells with pCAG-hypBase and pB-TREtight-EGFP (described above), then expanded and sorted on a BD FACS Aria, excluding the brightest 2% of EGFP cells, and keeping four of the next brightest EGFP cell populations. The sorted cells were expanded, frozen, and then thawed for testing their efficacy at titering ΔL-tTA virus. The fourth-brightest tranche of cells was used for subsequent titering of ΔL-tTA virus.

The 293T-FLEX-F14F15S-BC cell line for titering viruses expressing either Cre or Flpo was made by transfecting HEK-293T/17 cells with pCAG-hypBase and pB-CAG-FLEX-F14F15S-BFP-(mCherry)’-WPRE-BGHpA (described above), then expanded and sorted twice on a BD FAC Aria.

The 293T-FLEX-F14F15S-BC-TTBL titering cell line expressing SAD B19 L was made by infecting 293T-FLEX-F14F15S-BC cells (described above) with LV-TTBL(VSVG) (described above) at a multiplicity of infection of 18.6 infectious units per cell, then expanding, aliquoting, and freezing the cells.

### Rabies virus production and titering

The first-generation vectors RVΔG-4Cre(B19G) and RVΔG-4Flpo(B19G), the second-generation vectors RVΔGL-4Cre(B19G) and RVΔGL-4Flpo(B19G), and the third-generation vectors RVΔL-5Cre(B19G), RVΔL-5Flpo(B19G), and RVΔL-5tTA(B19G) were rescued as described previously (Chatterjee *et al*., 2018) using genome plasmids pRVΔGL-4Cre, pRVΔGL-4Flpo, pRVΔL-5Cre, pRVΔL-5Flpo, and pRVΔL-5tTA, respectively. For simplicity, these viruses are referred to in this manuscript as RVΔG-Cre, RVΔGL-Cre, RVΔGL-Flpo, RVΔL-Cre, RVΔL-Flpo, and RVΔL-tTA, omitting the numbers denoting the positions of the transgenes within the viral genomes as well as the “(B19G)” suffix denoting the SAD B19 rabies virus glycoprotein used to coat the viral particles. Rescue supernatants were collected and filtered as described (Wickersham and Sullivan, 2015), titered on the reporter cell lines 293T-FLEX BC (for Cre viruses) or 293T-F14F15S-BC (for Flpo viruses) (Jin *et al*., 2021) as described (Wickersham *et al*., 2010), then used to infect BHK-B19G3, BHK-B19L-G, or BHK-B19L cells (see above) at multiplicities of infection ranging from 0.1 to 1. Supernatants from these “P1” plates were collected and titered as described (Wickersham and Sullivan, 2015); in some cases, these were used for a similar second passage (“P2”). Purification and concentration of either P1 or P2 supernatants was as described (Wickersham *et al*., 2010), with supernatants treated with benzonase (Sigma 71206) (25 minute incubation at 37°C with 30 units/ml at) before ultracentrifugation. Concentrated viruses were aliquoted and frozen at -80°C. Rabies viruses were titered on reporter cells (293T-FLEX-BC for Cre viruses, 293T-F14F15S-BC for Flpo viruses, 293T-TREtight-EGFP (see above) for RVΔL-tTA) as described (Wickersham *et al*., 2010), using a LUNA-II cell counter (Logos Biosystems) instead of a hemocytometer for counting cells, and in some cases using two-fold (as opposed to ten-fold) dilution series for more precise comparisons of titers.

For direct comparison of the retrograde targeting efficacies of ΔGL and ΔL viruses expressing Cre and Flpo (Fig. 2), four rabies viruses – RVΔGL-4Cre(B19G), RVΔL-5Cre(B19G), RVΔGL-Flpo(B19G), and RVΔL-5Flpo(B19G) – were rescued in parallel in 15cm plates as described (Wickersham *et al*., 2010). Rescue supernatants were collected and filtered as described (Wickersham and Sullivan, 2015), titered on the reporter cell lines 293T-FLEX BC (for Cre viruses) or 293T-F14F15S-BC (for Flpo viruses) (Jin *et al*., 2021) as described (Wickersham *et al*., 2010), then passaged in parallel by application of rescue supernatant to one 15cm plate each of either BHK-B19L-G cells (described above) for the ΔGL viruses and BHK-B19L cells (described above) for the ΔL viruses at a multiplicity of 0.1. Medium was replaced with 13 ml fresh medium the following day, then supernatants from these P1 plates were collected for the three following days, with each supernatant filtered, treated with benzonase, concentrated by ultracentrifugation as described (Wickersham *et al*., 2010), then aliquoted and stored at -80°C. The concentrated third supernatant of each virus was used for injections (see below).

For direct comparison of the titers of ΔG, ΔGL, and ΔL viruses expressing Cre and Flpo (Supplementary Fig. S3), all six rabies viruses – RVΔG-4Cre(B19G), RVΔGL-4Cre(B19G), RVΔL-5Cre(B19G), RVΔG-4Flpo(B19G), RVΔGL-4Flpo(B19G), and RVΔL-5Flpo(B19G) – were rescued in parallel in 15cm plates as described (Wickersham *et al*., 2010), then passaged once in parallel by application of an equal volume of rescue supernatant (2.1 ml of the 5^th^ day’s collection from each virus’s rescue plate) to one 15cm plate of BHK-B19L-G (described above) for each virus. The day after infection, medium was removed and replaced with 13 ml fresh medium per plate; one day later, supernatants were collected, filtered, and replaced with 13 ml fresh medium per plate; one day after that, a second supernatant was collected from each plate. This second set of supernatants was used for titering, as follows:

293T-FLEX-F14F15S-BC and 293T -FLEX-F14F15S-BC-TTBL (described above) were infected with RVΔG-4Cre(B19G), RVΔGL-4Cre(B19G), RVΔL-5Cre(B19G), RVΔG-4Flpo(B19G), RVΔGL-4Flpo(B19G), and RVΔL-5Flpo(B19G) in three-fold dilution series, with each condition in triplicate, then fixed three days later, immunostained for rabies virus nucleoprotein (see below), and analyzed by flow cytometry to calculate titers from the percentage of cells in each well labeled by either mCherry or FITC as described (Wickersham *et al*., 2010).

### Immunostaining, imaging, and flow cytometry of cultured cells

For anti-nucleoprotein and anti-Cre staining (for Fig. 1): HEK 293T/17 (ATCC 11268) cells were plated on poly-L-lysine-coated coverslips in 24-well plates, then infected the following day with serial dilutions of RVΔG-4Cre(B19G) (Chatterjee *et al*., 2018), RVΔGL-4Cre(B19G) (Chatterjee *et al*., 2018), or RVΔL-5Cre(B19G). Three days after infection, cells were fixed with 2% paraformaldehyde, washed repeatedly with blocking/permeabilization buffer (0.1% Triton-X (Sigma) and 1% bovine serum albumin (Sigma) in PBS), then labeled with a 1:100 dilution of anti-nucleoprotein monoclonal antibody blend (Light Diagnostics Rabies DFA Reagent, Millipore Sigma 5100) as well as a 1:250 dilution of rabbit anti-Cre polyclonal antibody (Millipore Sigma 69050) followed by a 1:200 dilution of Alexa Fluor 594-conjugated donkey anti-rabbit secondary (Jackson Immuno 711-585-152).

For anti-EGFP staining (for Supplementary Fig. S1), HEK cells were plated as above, then infected the following day with serial dilutions of RVΔG-4EGFP(B19G) (Wickersham *et al*., 2010), RVΔGL-4EGFP(B19G) (Chatterjee *et al*., 2018), or RVΔL-5EGFP(B19G), with immunostaining three days postinfection, using a 1:1000 dilution of chicken anti-GFP polyclonal antibody (Aves Labs, GFP-1020) and a 1:500 dilution of Alexa Fluor 594-conjugated donkey anti-chicken secondary antibody (Jackson Immuno 703-585-155).

Immunostained cells on coverslips were mounted on slides using ProLong Diamond Antifade mounting medium (Thermo P36970) and imaged on a Zeiss LSM 900 confocal microscope using a 20× objective.

For matched flow cytometric analysis of immunostained cells, cells were plated in 24-well plates without poly-L-lysine-coated coverslips but otherwise immunostained as described above, then analyzed on an LSR II flow cytometer (BD) using FACS Diva software (BD). Histograms displayed in Fig. 1 were smoothed using the FACS Diva “Smooth histogram” setting.

### Viral growth analysis

For determining growth curves, BHK-B19G3, BHK-B19L-G, and BHK-B19L cells (see above) were plated in 10 cm plates coated in poly-L-lysine in normal medium (10% fetal bovine serum (VWR 16777-014) and antibiotic-antimycotic (Thermo 15240096) in DMEM (Thermo 11995073)) (Wickersham *et al*., 2010). The following day, cells were infected with RVΔG-4Cre(B19G), RVΔGL-4Cre(B19G), or RVΔL-5Cre(B19G) at an MOI of either 1 (for single-step growth curves) or 0.01 (for multi-step growth curves), with viruses diluted in normal medium at a total volume of 2 ml per plate, with each condition in triplicate. Following a one-hour incubation, the virus-containing medium was aspirated, plates were washed twice in DPBS (Thermo 14190144), and 12 ml fresh medium was added to each plate before they were returned to the incubator. Every 24 hours for the following five days, 200 µl of supernatant was collected from each plate; these supernatant samples were filter-sterilized using a 96-well 0.45um PVDF filter plate (Millipore MSHVN4510), then frozen at -80°C before all samples were thawed and titered on 293T-FLEX-BC cells as described above.

### Stereotaxic injections

200 nl of rabies virus was injected into either somatosensory thalamus (VPM/Po, for Fig. 2) or primary visual cortex (for two-photon experiments) of anesthetized adult mice using a stereotaxic instrument (Stoelting Co., 51925) and a custom injection apparatus consisting of a hydraulic manipulator (Narishige, MO-10) with headstage coupled via custom adaptors to a wire plunger advanced through pulled glass capillaries (Drummond, Wiretrol II) back-filled with mineral oil and front-filled with viral vector solution (Lavin et al., 2019). We have described this injection system in detail previously. Injection coordinates for VPM/Po were: anteroposterior (AP) = -1.82 mm with respect to (w.r.t.) bregma, lateromedial (LM) = +1.54 mm w.r.t bregma, dorsoventral (DV) = -3.15 mm w.r.t the brain surface; injection coordinates for V1 cortex were: AP = -2.70 mm w.r.t. bregma, LM = 2.50 mm w.r.t. bregma, DV = -0.26 mm w.r.t the brain surface.

For mice to be used for two-photon imaging, a 3 mm craniotomy was opened over primary visual cortex (V1). Glass windows composed of a 3mm-diameter glass coverslip (Warner Instruments CS-3R) glued (Optical Adhesive 61, Norland Products) to a 5mm-diameter glass coverslip (Warner Instruments CS-5R) were affixed over the craniotomy with Metabond (Parkell) after virus injection.

For the ΔGL vs. ΔL injections (Fig. 2), the four viruses were produced in parallel for direct comparison (see above), and RVΔL-5Cre(B19G) (6.16E+10 i.u./ml) or RVΔGL-4Cre(B19G) (3.01E+09 i.u./ml) was injected into Ai14 (het) mice, and RVΔL-5Flpo(B19G) (1.61E+09 i.u./ml) or RVΔGL-4Flpo(B19G) (1.12E+08 i.u./ml) was injected into Ai65F (het) mice. For the 4-month and 6-month experiments for Fig. 2, RVΔL-Cre (1.66E+10 i.u./ml) was injected into Ai14 (het) mice. For Supplementary Fig. S4, RVΔL-tTA (3.63E+10 iu/ml) was injected into Ai63 (het) mice.

For two-photon structural experiments (Fig. 4), RVΔGL-4Cre(B19G) (1.19E+10 iu/ml) or RVΔL-5Cre(B19G) (1.66E+10 iu/ml diluted to 1.19E+10 iu/ml for matching to RVΔGL-4Cre(B19G)) was injected into Ai14 (het) mice. For two-photon functional experiments in Fig. 6, RVΔL-5Cre(B19G) (2.61E+10 iu/ml) was injected into homo/homo Ai94D x ROSA:LNL:tTA mice.

For comparisons of RVΔL to other viral species (Fig. 3), 200 nl of each the following viruses was injected into either anterior cingulate area (ACA) or anteromedial visual cortex (AM) of heterozygous Ai14 mice: RVΔL-5Cre(B19G) (1.73E+10 iu/ml); CAV-2-Cre (“CAV-Cre” from Plateforme de Vectorologie de Montpellier, 6.9E+12 physical particles/ml); AAV2-retro-hSyn-Cre (Addgene 105553-AAVrg, lot v13700, 2.39E+13 gc/ml). Injection coordinates for ACA were: AP = 0.50 mm w.r.t. bregma, LM = -0.25 w.r.t. bregma, DV = 0.90 w.r.t. the brain surface; injection coordinates for AM were: AP = -2.18 w.r.t. bregma, LM = -1.60 w.r.t. bregma, DV = 0.55 w.r.t the brain surface. 4 mice were used for each virus at each injection site (24 mice total).

For *ex vivo* electrophysiological experiments, 200 nl of AAV2-retro-hSyn-Cre (2.39E+13 gc/ml from Addgene) or RVΔL-5Cre(B19G) (6.90E+10 iu/ml) viruses were injected into nucleus accumbens (AP=1.30 w.r.t. bregma, LM = 0.80 w.r.t. bregma, DV = -4.00 w.r.t. the brain surface) of 16 Ai14(het) mice. 4 weeks and 12 weeks after virus injection, 3 animals for each virus were used for slice electrophysiology experiments, and 1 animal per group was used for confocal imaging.

### Perfusions, histology, and confocal imaging

1 week to 6 months (see main text) after injection of rabies virus, anesthetized mice were transcardially perfused with 4% paraformaldehyde. Brains were postfixed overnight in 4% paraformaldehyde in PBS on a shaker at 4°C and cut into 50 μm coronal sections on a vibrating microtome (Leica, VT-1000S). Sections were collected sequentially into 6 tubes containing cryoprotectant, so that each tube contained every sixth section, then frozen at -20°C. Sections to be imaged were washed to remove cryoprotectant, then mounted with ProLong Diamond Antifade mounting medium (Thermo Fisher P36970) and imaged on a confocal microscope (Zeiss, LSM 900). For comparisons of RVΔL to other viral species, perfusion was 4 weeks after virus injection, and brain slices were mounted with ProLong Diamond Antifade Mountant with DAPI (Catalog number: P36971, ThermoFisher).

To ensure that the confocal images included in the figures are representative of each group, the images were taken after the counts were conducted, and the mouse with the next higher number of labeled neurons than the average number for its group was selected for confocal imaging.

### Quantification of retrograde targeting

Coronal sections between 0.43mm anterior and 4.07mm posterior to bregma were imaged with an epifluorescence microscope for cell counting (Zeiss, Imager.Z2). Due to the high density of retrogradely labeled tdTomato neurons in the cortex at the injection site (VPM/Po), cells were counted using the Analyze Particle function in ImageJ (size in micron^2: 20-400; circularity: 0.20-1.00). Only one of the six series of sections (i.e., every sixth section: see above) was counted for each mouse.

### Quantification of spine density

Sparsely-labeled basal dendritic spines were imaged on a Zeiss LSM 900 confocal microscope using a 63X oil-immersion objective to take 17-to 47-section z-stacks with 0.5 µm distance between each optical section. Images were viewed in ImageJ for analysis, with spines counted manually and process length measured using the Fiji segmented line tool.

### Structural two-photon imaging and image analysis

Beginning seven days after injection of each rabies virus and recurring at the subsequent indicated timepoints (see main text) up to a maximum of 16 weeks following rabies virus injection, fields of view (FOVs) were imaged on a Prairie/Bruker Ultima IV In Vivo two-photon microscope driven by a Spectra Physics Mai-Tai Deep See laser with a mode locked Ti:sapphire laser emitting at a wavelength of 1020 nm for excitation of tdTomato. In order to distinguish individual labeled neurons, FOVs were chosen some distance away from the area of brightest tdTomato labeling. Two well-separated areas were chosen in each mouse. For each imaging session, mice were reanesthetized and mounted via their headplates to a custom frame, with ointment applied to protect their eyes and with a handwarmer maintaining body temperature. Imaging parameters were as follows: image size 512 X 512 pixels (282.6 μm × 282.6 μm), 0.782 Hz frame rate, dwell time 4.0 μs, 2× optical zoom, Z-stack step size 1 μm. Image acquisition was controlled with Prairie View 5.4 software. Laser power exiting the 20× water-immersion objective (Zeiss, W plan-apochromat, NA 1.0) varied between 20 and 65 mW depending on focal plane depth (Pockels cell value was automatically increased from 450 at the top section of each stack to 750 at the bottom section). For the example images of labeled cells, maximum intensity projections (stacks of 100-200 μm) were made with ImageJ software. Cell counting was automated using the “Analyze Particles” function in ImageJ.

### Brain slice electrophysiology

4 weeks and 12 weeks after injection of RVΔL-5Cre(B19G) or AAV2-retro-hSyn-Cre (see above), coronal brain slices containing the BLA were collected from 3 mice per virus for *ex vivo* patch clamp recordings. Mice were anesthetized with isoflurane, perfused with ice cold cutting solution, and decapitated using scissors. Brains were extracted and immersed in ice-cold (0–4 °C) sucrose cutting solution containing (in mM) 252 sucrose, 26 NaHCO3, 2.5 KCl, 1.25 NaH2PO4, 1 CaCl2, 5 MgCl2 and 10 glucose, which was oxygenated with 95% O2 and 5% CO2. The brains were trimmed and coronal brain slices (300 µm) were sectioned using a vibrating microtome (VT1200, Leica). After sectioning, slices for patch clamp recordings were transferred to a holding chamber containing oxygenated patch clamp recording medium (artificial cerebrospinal fluid, aCSF) containing (in mM): 126 NaCl, 2.5 KCl, 1.25 NaH2PO4, 1.3 MgCl2, 2.5 CaCl2, 26 NaHCO3, and 10 glucose, where they were maintained at 32 °C for 30 min before the chamber temperature was decreased to ∼20 °C. Slices were transferred one at a time from the holding chamber to a submerged recording chamber mounted on the fixed stage of an Olympus BX51WI fluorescence microscope equipped with differential interference contrast (DIC) illumination. The slices in the recording chamber were continuously perfused at a rate of 2 ml/min with recording aCSF at room temperature and continuously aerated with 95% O2/5% CO2. Glass pipettes with a resistance of 4-6 MΩ were pulled from borosilicate glass (ID 1.2 mm, OD 1.65 mm) on a horizontal puller (Sutter P-97) and filled with an intracellular patch solution containing (in mM): 130 potassium gluconate, 10 HEPES, 10 phosphocreatine Na2, 4 Mg-ATP, 0.4 Na-GTP, 5 KCl, 0.6 EGTA; pH was adjusted to 7.25 with KOH and the solution had a final osmolarity of 293 mOsm. Fluorescently labeled neurons in basolateral amygdala were selected for whole-cell patch clamp recordings. Series resistance was continuously monitored, and cells were discarded when the series resistance changed more than 20%. Data were acquired using a Multiclamp 700B amplifier, a Digidata 1440 A analog/digital interface, and pClamp 10 software (Molecular Devices).

### Functional two-photon imaging and image analysis

Functional two-photon imaging of RVΔL-5Cre(B19G)-labeled cells began at 2 weeks after injection of each rabies virus and recurred at timepoints of 3, 4, 5, 6, 8, 10, 12, 14, and 16 weeks following rabies virus injection (the FOVs were inspected using the two-photon microscope at 7 days postinfection, but images were not collected at this timepoint because few cells were found; at the 16 week timepoint, only one mouse had window clarity sufficient for imaging). FOVs were slightly offset from the regions of brightest GCaMP6s label in left-hemisphere V1 in order to allow separate identification of individual cells. This imaging was performed using the same microscope (5.356-Hz frame rate, 1024X 128 pixels, 565.1 μm × 565.1 μm, dwell time 0.8 μs, 1× optical zoom, scan angle 45 degree) with the same objective and laser (at 920 nm) as for the structural imaging experiments. Laser power at the objective ranged from 10 to 65 mW. Calcium imaging data were acquired in supragranular layers (100 to 200 μm deep). Surface vasculature provided coarse fiducial markers for finding the same FOVs in different imaging sessions. For these experiments, mice were awake and head-fixed. No behavioral training or reward was given. Visual stimuli were generated in Matlab (R2015R version) with custom software based on Psychtoolbox (http://psychtoolbox.org) and shown on the same LCD screen as in the widefield mapping experiments. Each condition consisted of 2 s of a full-field sine wave grating drifting in one direction, presented at 80% contrast with spatial frequency of 0.04 cycles/ degree, followed by 2 s of uniform mean luminance (gray). All permutations of 12 directions (30° steps) and 5 temporal frequencies (1, 2, 4, 8 and 15 Hz) were shown, in randomized order. The complete set was repeated 10 times, for a total stimulation period of 40 min per FOV per session. Cells were then manually segmented, and single-cell fluorescence traces were extracted by averaging the fluorescence of all pixels masking the soma, using ImageJ (version 2-0-0-rc-69) software. The mean ΔF/F over the full 2 s of each stimulus condition was used to calculate orientation tuning curves, with background fluorescence (F) in Δ F/F taken as the value of the trace immediately preceding a condition, averaged over all conditions. The raw calcium traces from cells within individual FOVs (not across FOVs, given different imaging conditions across animals and time points) were sorted by mean fluorescence. Randomly colored ROI view images were created by suite2p (https://www.suite2p.org). For ‘tuned’ cells in Fig. 6 panels E – G, the counts are based on all imaged neurons’ individual tuning curves, plotted in MATLAB; any cell showing response to a preferred orientation (including narrowly tuned neurons and broadly tuned neurons) at any temporal frequency (1Hz, 2Hz, 4Hz, 8Hz, or 15Hz) was counted manually as a tuned cell.

## QUANTIFICATION AND STATISTICAL ANALYSIS

All cell and spine counts and statistical analyses, including exact values of n, what n represents in each case, etc. are provided in Supplemental Information. Statistical analyses and plots of cell and spine counts were made with Prism 9 (GraphPad Software, San Diego, California). P-values for all comparisons were obtained using single-factor ANOVAs. For the comparisons between RVΔL, rAAV2-retro, and CAV-2, either one-way (for hippocampus, BLA, and cortex) or two-way (for cortical layers in primary motor and somatosensory cortex) ANOVAs with Tukey’s multiple comparison test were used. Power analyses were conducted using the online calculator at https://www.stat.ubc.ca/~rollin/stats/ssize/n2.html.

Slice electrophysiology data were analyzed with Clampfit 10 (Molecular Devices). The membrane capacitance (Cm) and input resistance (Rm) were calculated from a membrane seal test conducted in voltage-clamp mode, in which 100-ms, 5-mV voltage steps were delivered at a frequency of 5 Hz. In order to evaluate the action potential rheobase, a 1-s, positive current was delivered in current-clamp mode at 10pA steps. Rheobase was defined as the minimum current required to depolarize the membrane potential for action potential firing. Finally, to evaluate the relationship between injected current and firing frequency, we delivered a series of 0.5s current pulses in current-clamp in 50pA steps from 50 to 350 pA. The numbers of recorded cells in the four groups were as follows: rAAV2-retro-hSyn-Cre: n=28 cells at 4 weeks, n=27 cells at 12 weeks; RVΔL-5Cre(B19G): n=25 cells at 4 weeks, n=28 cells at 12 weeks. Statistical comparisons were conducted with one- or two-way ANOVAs, as follows: resting membrane potential: one-way ANOVA, F(3, 104) = 0.2957, p = 0.8284; membrane capacitance (Cm): one-way ANOVA, F(3, 104) = 0.2876, p = 0.8343; rheobase: one-way ANOVA, F(3, 104) = 1.491, p = 0.2214; input resistance (Rm): one-way ANOVA, F(3, 104) = 1.039, p = 0.3785; action potential number vs. input current: two-way ANOVA, F(3, 104) = 1.034, p = 0.3809.

