## Supplementary Information for "Single-deletion-mutant, third-generation rabies viral vectors allow nontoxic retrograde targeting of projection neurons with greatly increased efficiency"

### SUPPLEMENTAL INFORMATION

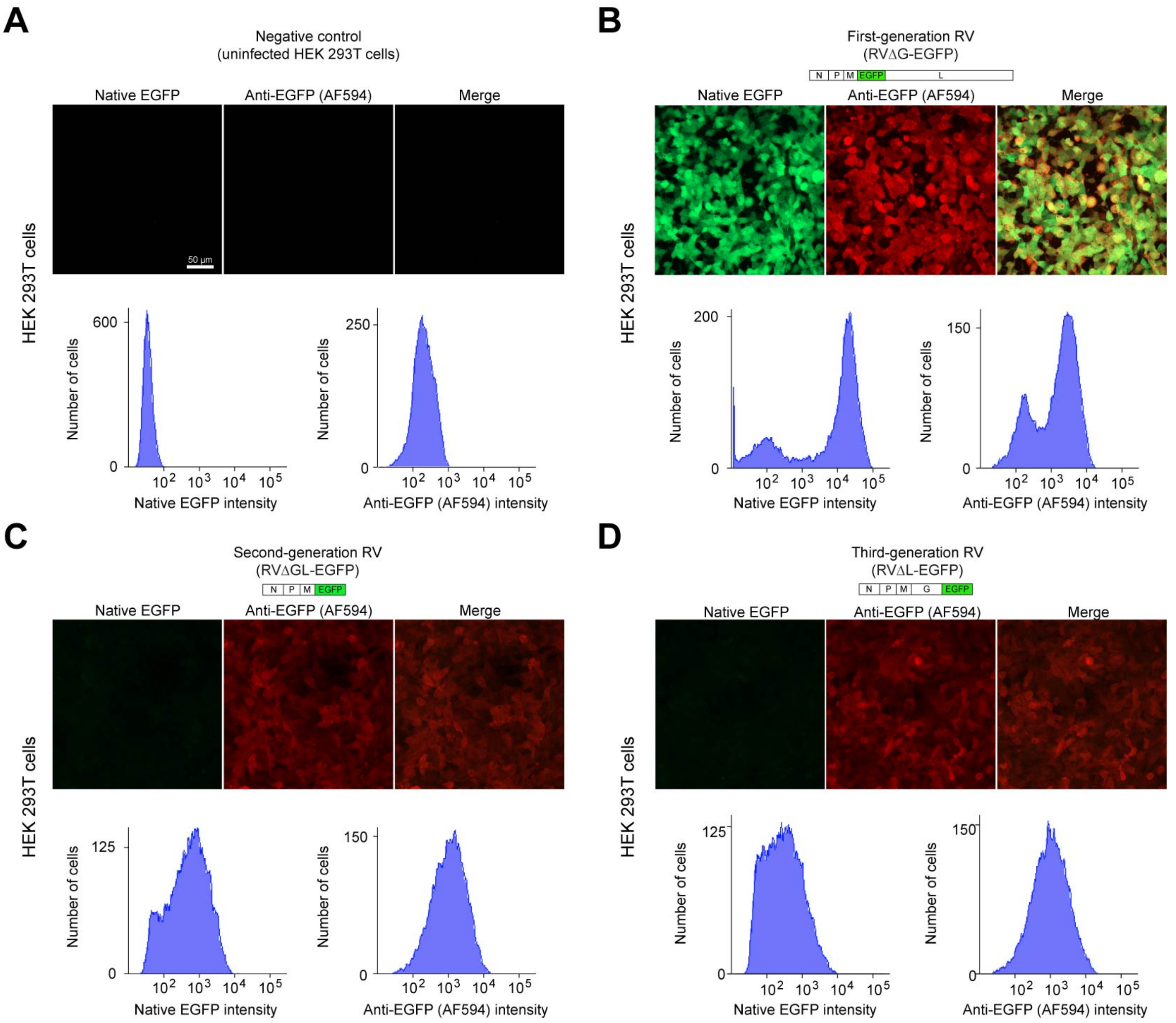

**Supplementary Figure S1. ΔL and ΔGL viruses express EGFP at similarly low levels, Related to Figure 1.**
Confocal images and flow cytometric histograms showing native and immunostained EGFP signal in uninfected cells (A) and cells infected with first-generation (ΔG) virus (B), second-generation (ΔGL) virus (C), or third-generation virus (D) expressing EGFP. Scale bar: 50 µm, applies to all images.

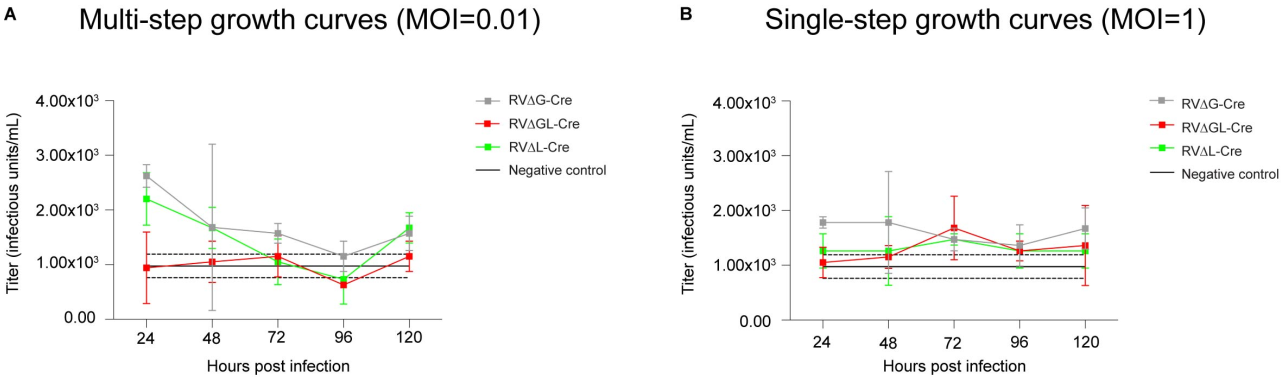

**Supplementary Figure S2. ΔG, ΔGL, and ΔL viruses do not propagate in non-complementing cells, Related to Figure 1.**

Viral titers in supernatants of BHK-21 cells not expressing any rabies viral genes, infected with ΔL, ΔGL, or ΔG viruses at a multiplicity of infection (MOI) of 0.01 ("multi-step growth curves", panel A) or 1 ("single-step" growth curves, panel B), with supernatants collected every 24 hours for five days. Graphs show mean ± s.e.m. .Black lines show negative control "titers" calculated from uninfected reporter cells (mean ± s.e.m. of 10 samples). Note that the titers in these graphs are 3-4 orders of magnitude lower than those obtained on complementing cells (Figure 1).

**Supplementary File S1. Titers and statistics for growth dynamics experiments, Related to Figure 1.** Fields highlighted in green are for timepoints with the highest mean titer for each condition (a given virus grown on a given cell line); these highest titers for each condition were used for the statistical comparisons with the other conditions.

*See following pages.*

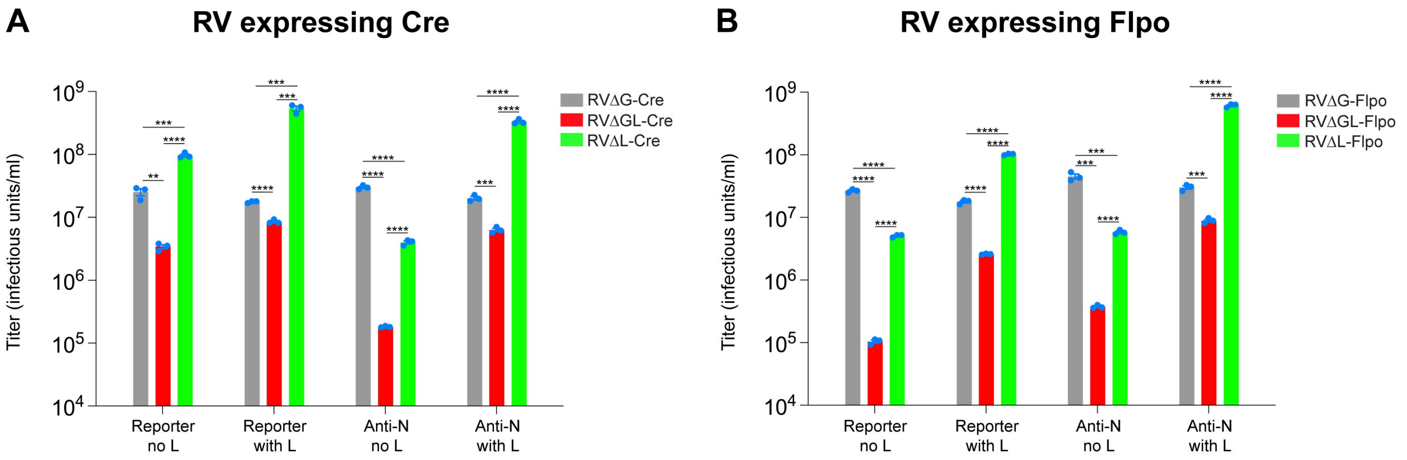

**Supplementary Figure S3. Direct comparisons of titers of ∆G, ∆GL, and ∆L viruses expressing Cre and Flpo, each titered in four different ways, Related to Figure 1.**
In order to directly compare the titers of the Cre- and Flpo-expressing versions of each of the three generations of rabies viral vectors, we made a fluorescent reporter cell line (293T-FLEX-F14F15S-BC) in which either Cre or Flpo activity causes mCherry expression (see Methods). We further made a version of this reporter line that expresses the rabies viral polymerase gene (293T-FLEX-F14F15S-BC-TTBL, see Methods), in order to amplify the low recombinase expression from ∆GL and ∆L vectors and thereby unmask subthreshold expression. In the case of ∆L viruses, but not ∆GL ones, this L expression would also allow replication and spread between cells, confounding interpretation; despite this, we titered the vectors of all three generations with and without L for completeness. After making side-by-side preparations of six viruses, RV∆G-Cre, RV∆GL-Cre, RV∆L-Cre, RV∆G-Flpo, RV∆GL-Flpo, and RV∆L-Flpo (see Methods), we titered them in triplicate by serial dilution on the above two dual reporter cell lines. For an additional readout that was independent of recombinase efficacy, we also immunostained for the rabies virus nucleoprotein, giving a total of four titering methods per virus (mCherry expression versus anti-nucleoprotein signal, with and without L). Noteworthy findings include the following:

1) The titer of each of the two ∆L viruses significantly exceeds that of the corresponding ∆GL virus using any of the four titering methods: i.e., the titer of RV∆L-Cre is significantly greater than that of RV∆GL-Cre, and the titer of RV∆L-Flpo is significantly greater than that of RV∆GL-Flpo, regardless of titering method.
2) For both ∆GL and ∆L vectors (but not ∆G vectors), the titer of the Cre-expressing version was significantly higher than that of the Flpo-expressing versions, when assessed by fluorescent reporter expression: i.e., the measured titer of RV∆L-Cre is significantly greater than that of RV∆L-Flpo, and the titer of RV∆GL-Cre is significantly greater than that of RV∆GL-Flpo, when titering based on reporter expression (and regardless of whether L was expressed *in trans*). However, when nucleoprotein staining was used to determine the presence of virus, the titer of each of the Flpo-expressing ∆GL and ∆L viruses were actually higher than that of the corresponding Cre-expressing one: that is, the titer of RV∆L-Flpo is significantly greater than that of RV∆L-Cre, and the titer of RV∆GL-Flpo is significantly greater than that of RV∆GL-Flpo, when titering using nucleoprotein staining, regardless of whether L was expressed *in trans*. These findings are consistent with an interpretation that the lower efficacy of Flpo versus that of Cre, coupled with the intentionally low expression levels of ∆GL and ∆L vectors, result in lower *de facto* titers (as judged by the percentage of mCherry-labeled reporter cells) and lower numbers of retrogradely labeled neurons *in vivo* (See Figure 2).
3) Expression of L *in trans* significantly increases the measured titers of all four ∆GL and ∆L viruses, whether measured by fluorescent reporter expression or nucleoprotein staining. This is also consistent with the above hypothesis that the intentionally low expression levels characteristic of these vectors may result in artificially low measured titers because of subthreshold expression of recombinase or nucleoprotein.

See Supplementary File S2 for titers and statistical comparisons.

**Supplementary File S2. Titers and statistical comparisons of titers of ΔG, ΔGL, and ΔL viruses expressing Cre and Flpo, each titered in four different ways, Related to Figure 1.**
Titer values for each sample, followed by statistical comparisons and p-values.

*See following pages.*

**Supplementary File S3. Cell counts and statistics for ∆GL vs ∆L retrograde targeting experiments, Related to Figure 2.**

Cell counts for each individual 50-μm section, followed by the total cell count for each series (of every sixth section) for each mouse, followed by statistical comparisons and p-values.

*See following pages.*

| Condition | Mouse number | Section number | Cell counts |
|---|---|---|---|
| RVΔGL-Flpo: 1 week | 21021505LJ | 10 | 5 |
| RVΔGL-Flpo: 1 week | 21021505LJ | 11 | 8 |
| RVΔGL-Flpo: 1 week | 21021505LJ | 12 | 9 |
| RVΔGL-Flpo: 1 week | 21021505LJ | total: | 22 |
| RVΔGL-Flpo: 1 week | 21021506LJ | 9 | 5 |
| RVΔGL-Flpo: 1 week | 21021506LJ | 10 | 6 |
| RVΔGL-Flpo: 1 week | 21021506LJ | 11 | 4 |
| RVΔGL-Flpo: 1 week | 21021506LJ | 12 | 3 |
| RVΔGL-Flpo: 1 week | 21021506LJ | total: | 18 |
| RVΔGL-Flpo: 1 week | 21021507LJ | 5 | 1 |
| RVΔGL-Flpo: 1 week | 21021507LJ | 6 | 27 |
| RVΔGL-Flpo: 1 week | 21021507LJ | 7 | 29 |
| RVΔGL-Flpo: 1 week | 21021507LJ | 8 | 25 |
| RVΔGL-Flpo: 1 week | 21021507LJ | 9 | 20 |
| RVΔGL-Flpo: 1 week | 21021507LJ | 10 | 10 |
| RVΔGL-Flpo: 1 week | 21021507LJ | 11 | 4 |
| RVΔGL-Flpo: 1 week | 21021507LJ | 12 | 3 |
| RVΔGL-Flpo: 1 week | 21021507LJ | 13 | 2 |
| RVΔGL-Flpo: 1 week | 21021507LJ | total: | 121 |
| RVΔGL-Flpo: 1 week | 21021508LJ | 11 | 1 |
| RVΔGL-Flpo: 1 week | 21021508LJ | 15 | 1 |
| RVΔGL-Flpo: 1 week | 21021508LJ | total: | 2 |
| RVΔGL-Flpo: 1 week | 21052505LJ | 9 | 1 |
| RVΔGL-Flpo: 1 week | 21052505LJ | 10 | 5 |
| RVΔGL-Flpo: 1 week | 21052505LJ | 11 | 1 |
| RVΔGL-Flpo: 1 week | 21052505LJ | total: | 7 |
| RVΔGL-Flpo: 1 week | 21052506LJ | 11 | 4 |
| RVΔGL-Flpo: 1 week | 21052506LJ | 12 | 4 |
| RVΔGL-Flpo: 1 week | 21052506LJ | total: | 8 |
| RVΔGL-Flpo: 1 week | 21052507LJ | 6 | 2 |
| RVΔGL-Flpo: 1 week | 21052507LJ | 7 | 20 |
| RVΔGL-Flpo: 1 week | 21052507LJ | 8 | 45 |
| RVΔGL-Flpo: 1 week | 21052507LJ | 9 | 26 |
| RVΔGL-Flpo: 1 week | 21052507LJ | 10 | 5 |
| RVΔGL-Flpo: 1 week | 21052507LJ | total: | 98 |
| RVΔGL-Flpo: 1 week | 21052508LJ | 8 | 1 |
| RVΔGL-Flpo: 1 week | 21052508LJ | 10 | 1 |
| RVΔGL-Flpo: 1 week | 21052508LJ | 11 | 1 |
| RVΔGL-Flpo: 1 week | 21052508LJ | 12 | 1 |
| RVΔGL-Flpo: 1 week | 21052508LJ | 14 | 2 |
| RVΔGL-Flpo: 1 week | 21052508LJ | total: | 6 |
| RVΔL-Flpo: 1 week | 21021805LJ | 1 | 57 |
| RVΔL-Flpo: 1 week | 21021805LJ | 2 | 20 |
| RVΔL-Flpo: 1 week | 21021805LJ | 3 | 11 |
| RVΔL-Flpo: 1 week | 21021805LJ | 4 | 115 |
| RVΔL-Flpo: 1 week | 21021805LJ | 5 | 270 |
| RVΔL-Flpo: 1 week | 21021805LJ | 6 | 399 |
| RVΔL-Flpo: 1 week | 21021805LJ | 7 | 428 |
| RVΔL-Flpo: 1 week | 21021805LJ | 8 | 542 |
| RVΔL-Flpo: 1 week | 21021805LJ | 9 | 416 |
| RVΔL-Flpo: 1 week | 21021805LJ | 10 | 368 |
| RVΔL-Flpo: 1 week | 21021805LJ | 11 | 240 |
| RVΔL-Flpo: 1 week | 21021805LJ | 12 | 106 |
| RVΔL-Flpo: 1 week | 21021805LJ | 13 | 42 |

| | | | |
|---|---|---|---|
| RVΔL-Flpo: 1 week | 21021805LJ | 14 | 52 |
| RVΔL-Flpo: 1 week | 21021805LJ | 15 | 27 |
| RVΔL-Flpo: 1 week | 21021805LJ | total: | 3093 |
| RVΔL-Flpo: 1 week | 21021806LJ | 1 | 1 |
| RVΔL-Flpo: 1 week | 21021806LJ | 4 | 5 |
| RVΔL-Flpo: 1 week | 21021806LJ | 5 | 3 |
| RVΔL-Flpo: 1 week | 21021806LJ | 6 | 4 |
| RVΔL-Flpo: 1 week | 21021806LJ | 7 | 16 |
| RVΔL-Flpo: 1 week | 21021806LJ | 8 | 35 |
| RVΔL-Flpo: 1 week | 21021806LJ | 9 | 93 |
| RVΔL-Flpo: 1 week | 21021806LJ | 10 | 74 |
| RVΔL-Flpo: 1 week | 21021806LJ | 11 | 40 |
| RVΔL-Flpo: 1 week | 21021806LJ | 12 | 24 |
| RVΔL-Flpo: 1 week | 21021806LJ | 13 | 11 |
| RVΔL-Flpo: 1 week | 21021806LJ | 14 | 7 |
| RVΔL-Flpo: 1 week | 21021806LJ | 15 | 7 |
| RVΔL-Flpo: 1 week | 21021806LJ | total: | 320 |
| RVΔL-Flpo: 1 week | 21021807LJ | 5 | 1 |
| RVΔL-Flpo: 1 week | 21021807LJ | 6 | 2 |
| RVΔL-Flpo: 1 week | 21021807LJ | 7 | 6 |
| RVΔL-Flpo: 1 week | 21021807LJ | 8 | 8 |
| RVΔL-Flpo: 1 week | 21021807LJ | 9 | 35 |
| RVΔL-Flpo: 1 week | 21021807LJ | 10 | 64 |
| RVΔL-Flpo: 1 week | 21021807LJ | 11 | 80 |
| RVΔL-Flpo: 1 week | 21021807LJ | 12 | 55 |
| RVΔL-Flpo: 1 week | 21021807LJ | 13 | 34 |
| RVΔL-Flpo: 1 week | 21021807LJ | 14 | 13 |
| RVΔL-Flpo: 1 week | 21021807LJ | 15 | 13 |
| RVΔL-Flpo: 1 week | 21021807LJ | total: | 311 |
| RVΔL-Flpo: 1 week | 21021808LJ | 6 | 7 |
| RVΔL-Flpo: 1 week | 21021808LJ | 7 | 13 |
| RVΔL-Flpo: 1 week | 21021808LJ | 8 | 48 |
| RVΔL-Flpo: 1 week | 21021808LJ | 9 | 74 |
| RVΔL-Flpo: 1 week | 21021808LJ | 10 | 57 |
| RVΔL-Flpo: 1 week | 21021808LJ | 11 | 20 |
| RVΔL-Flpo: 1 week | 21021808LJ | 12 | 13 |
| RVΔL-Flpo: 1 week | 21021808LJ | 13 | 13 |
| RVΔL-Flpo: 1 week | 21021808LJ | 14 | 14 |
| RVΔL-Flpo: 1 week | 21021808LJ | 15 | 4 |
| RVΔL-Flpo: 1 week | 21021808LJ | total: | 263 |
| RVΔL-Flpo: 1 week | 21052607LJ | 4 | 3 |
| RVΔL-Flpo: 1 week | 21052607LJ | 5 | 2 |
| RVΔL-Flpo: 1 week | 21052607LJ | 6 | 4 |
| RVΔL-Flpo: 1 week | 21052607LJ | 7 | 11 |
| RVΔL-Flpo: 1 week | 21052607LJ | 8 | 10 |
| RVΔL-Flpo: 1 week | 21052607LJ | 9 | 4 |
| RVΔL-Flpo: 1 week | 21052607LJ | 11 | 6 |
| RVΔL-Flpo: 1 week | 21052607LJ | 12 | 7 |
| RVΔL-Flpo: 1 week | 21052607LJ | total: | 47 |
| RVΔL-Flpo: 1 week | 21052608LJ | 1 | 105 |
| RVΔL-Flpo: 1 week | 21052608LJ | 2 | 116 |
| RVΔL-Flpo: 1 week | 21052608LJ | 3 | 110 |
| RVΔL-Flpo: 1 week | 21052608LJ | 4 | 192 |
| RVΔL-Flpo: 1 week | 21052608LJ | 5 | 395 |
| RVΔL-Flpo: 1 week | 21052608LJ | 6 | 389 |

| | | | |
|---|---|---|---|
| RVΔL-Flpo: 1 week | 21052608LJ | 7 | 421 |
| RVΔL-Flpo: 1 week | 21052608LJ | 8 | 340 |
| RVΔL-Flpo: 1 week | 21052608LJ | 9 | 72 |
| RVΔL-Flpo: 1 week | 21052608LJ | 10 | 14 |
| RVΔL-Flpo: 1 week | 21052608LJ | 11 | 5 |
| RVΔL-Flpo: 1 week | 21052608LJ | 12 | 2 |
| RVΔL-Flpo: 1 week | 21052608LJ | total: | 2161 |
| RVΔL-Flpo: 1 week | 21060302LJ | 1 | 8 |
| RVΔL-Flpo: 1 week | 21060302LJ | 2 | 10 |
| RVΔL-Flpo: 1 week | 21060302LJ | 3 | 17 |
| RVΔL-Flpo: 1 week | 21060302LJ | 4 | 24 |
| RVΔL-Flpo: 1 week | 21060302LJ | 5 | 22 |
| RVΔL-Flpo: 1 week | 21060302LJ | 6 | 29 |
| RVΔL-Flpo: 1 week | 21060302LJ | 7 | 34 |
| RVΔL-Flpo: 1 week | 21060302LJ | 8 | 59 |
| RVΔL-Flpo: 1 week | 21060302LJ | 9 | 78 |
| RVΔL-Flpo: 1 week | 21060302LJ | 10 | 46 |
| RVΔL-Flpo: 1 week | 21060302LJ | 11 | 16 |
| RVΔL-Flpo: 1 week | 21060302LJ | 12 | 12 |
| RVΔL-Flpo: 1 week | 21060302LJ | 13 | 6 |
| RVΔL-Flpo: 1 week | 21060302LJ | 14 | 2 |
| RVΔL-Flpo: 1 week | 21060302LJ | 15 | 2 |
| RVΔL-Flpo: 1 week | 21060302LJ | total: | 365 |
| RVΔL-Flpo: 1 week | 21060303LJ | 1 | 7 |
| RVΔL-Flpo: 1 week | 21060303LJ | 2 | 9 |
| RVΔL-Flpo: 1 week | 21060303LJ | 3 | 10 |
| RVΔL-Flpo: 1 week | 21060303LJ | 4 | 16 |
| RVΔL-Flpo: 1 week | 21060303LJ | 5 | 22 |
| RVΔL-Flpo: 1 week | 21060303LJ | 6 | 27 |
| RVΔL-Flpo: 1 week | 21060303LJ | 7 | 25 |
| RVΔL-Flpo: 1 week | 21060303LJ | 8 | 27 |
| RVΔL-Flpo: 1 week | 21060303LJ | 9 | 36 |
| RVΔL-Flpo: 1 week | 21060303LJ | 10 | 30 |
| RVΔL-Flpo: 1 week | 21060303LJ | 11 | 15 |
| RVΔL-Flpo: 1 week | 21060303LJ | 12 | 7 |
| RVΔL-Flpo: 1 week | 21060303LJ | 14 | 6 |
| RVΔL-Flpo: 1 week | 21060303LJ | 15 | 1 |
| RVΔL-Flpo: 1 week | 21060303LJ | total: | 238 |
| RVΔGL-Flpo: 4 weeks | 21021501LJ | 7 | 2 |
| RVΔGL-Flpo: 4 weeks | 21021501LJ | 8 | 12 |
| RVΔGL-Flpo: 4 weeks | 21021501LJ | 9 | 25 |
| RVΔGL-Flpo: 4 weeks | 21021501LJ | 10 | 26 |
| RVΔGL-Flpo: 4 weeks | 21021501LJ | 11 | 7 |
| RVΔGL-Flpo: 4 weeks | 21021501LJ | 12 | 3 |
| RVΔGL-Flpo: 4 weeks | 21021501LJ | 13 | 3 |
| RVΔGL-Flpo: 4 weeks | 21021501LJ | 14 | 1 |
| RVΔGL-Flpo: 4 weeks | 21021501LJ | 15 | 1 |
| RVΔGL-Flpo: 4 weeks | 21021501LJ | total: | 80 |
| RVΔGL-Flpo: 4 weeks | 21021502LJ | 9 | 2 |
| RVΔGL-Flpo: 4 weeks | 21021502LJ | 10 | 25 |
| RVΔGL-Flpo: 4 weeks | 21021502LJ | 11 | 26 |
| RVΔGL-Flpo: 4 weeks | 21021502LJ | 12 | 5 |
| RVΔGL-Flpo: 4 weeks | 21021502LJ | 13 | 4 |
| RVΔGL-Flpo: 4 weeks | 21021502LJ | total: | 62 |
| RVΔGL-Flpo: 4 weeks | 21021503LJ | 1 | 88 |

| | | | |
|---|---|---|---|
| RVΔGL-Flpo: 4 weeks | 21021503LJ | 2 | 129 |
| RVΔGL-Flpo: 4 weeks | 21021503LJ | 3 | 199 |
| RVΔGL-Flpo: 4 weeks | 21021503LJ | 4 | 254 |
| RVΔGL-Flpo: 4 weeks | 21021503LJ | 5 | 312 |
| RVΔGL-Flpo: 4 weeks | 21021503LJ | 6 | 396 |
| RVΔGL-Flpo: 4 weeks | 21021503LJ | 7 | 351 |
| RVΔGL-Flpo: 4 weeks | 21021503LJ | 8 | 391 |
| RVΔGL-Flpo: 4 weeks | 21021503LJ | 9 | 359 |
| RVΔGL-Flpo: 4 weeks | 21021503LJ | 10 | 295 |
| RVΔGL-Flpo: 4 weeks | 21021503LJ | 11 | 133 |
| RVΔGL-Flpo: 4 weeks | 21021503LJ | 12 | 216 |
| RVΔGL-Flpo: 4 weeks | 21021503LJ | 13 | 59 |
| RVΔGL-Flpo: 4 weeks | 21021503LJ | 14 | 40 |
| RVΔGL-Flpo: 4 weeks | 21021503LJ | 15 | 32 |
| RVΔGL-Flpo: 4 weeks | 21021503LJ | total: | 3254 |
| RVΔGL-Flpo: 4 weeks | 21021504LJ | 3 | 1 |
| RVΔGL-Flpo: 4 weeks | 21021504LJ | 4 | 4 |
| RVΔGL-Flpo: 4 weeks | 21021504LJ | 5 | 22 |
| RVΔGL-Flpo: 4 weeks | 21021504LJ | 6 | 24 |
| RVΔGL-Flpo: 4 weeks | 21021504LJ | 7 | 56 |
| RVΔGL-Flpo: 4 weeks | 21021504LJ | 8 | 78 |
| RVΔGL-Flpo: 4 weeks | 21021504LJ | 9 | 32 |
| RVΔGL-Flpo: 4 weeks | 21021504LJ | 10 | 18 |
| RVΔGL-Flpo: 4 weeks | 21021504LJ | 11 | 1 |
| RVΔGL-Flpo: 4 weeks | 21021504LJ | 12 | 1 |
| RVΔGL-Flpo: 4 weeks | 21021504LJ | total: | 237 |
| RVΔGL-Flpo: 4 weeks | 21052501LJ | 2 | 1 |
| RVΔGL-Flpo: 4 weeks | 21052501LJ | 4 | 1 |
| RVΔGL-Flpo: 4 weeks | 21052501LJ | 7 | 2 |
| RVΔGL-Flpo: 4 weeks | 21052501LJ | 8 | 6 |
| RVΔGL-Flpo: 4 weeks | 21052501LJ | 9 | 9 |
| RVΔGL-Flpo: 4 weeks | 21052501LJ | 10 | 9 |
| RVΔGL-Flpo: 4 weeks | 21052501LJ | 11 | 2 |
| RVΔGL-Flpo: 4 weeks | 21052501LJ | 12 | 6 |
| RVΔGL-Flpo: 4 weeks | 21052501LJ | 13 | 4 |
| RVΔGL-Flpo: 4 weeks | 21052501LJ | 14 | 1 |
| RVΔGL-Flpo: 4 weeks | 21052501LJ | 15 | 2 |
| RVΔGL-Flpo: 4 weeks | 21052501LJ | total: | 43 |
| RVΔGL-Flpo: 4 weeks | 21052502LJ | 6 | 2 |
| RVΔGL-Flpo: 4 weeks | 21052502LJ | 7 | 1 |
| RVΔGL-Flpo: 4 weeks | 21052502LJ | 8 | 4 |
| RVΔGL-Flpo: 4 weeks | 21052502LJ | 11 | 1 |
| RVΔGL-Flpo: 4 weeks | 21052502LJ | 14 | 3 |
| RVΔGL-Flpo: 4 weeks | 21052502LJ | 15 | 3 |
| RVΔGL-Flpo: 4 weeks | 21052502LJ | total: | 14 |
| RVΔGL-Flpo: 4 weeks | 21052503LJ | 6 | 2 |
| RVΔGL-Flpo: 4 weeks | 21052503LJ | 8 | 1 |
| RVΔGL-Flpo: 4 weeks | 21052503LJ | 9 | 9 |
| RVΔGL-Flpo: 4 weeks | 21052503LJ | 10 | 45 |
| RVΔGL-Flpo: 4 weeks | 21052503LJ | 11 | 33 |
| RVΔGL-Flpo: 4 weeks | 21052503LJ | 12 | 17 |
| RVΔGL-Flpo: 4 weeks | 21052503LJ | 13 | 10 |
| RVΔGL-Flpo: 4 weeks | 21052503LJ | 14 | 7 |
| RVΔGL-Flpo: 4 weeks | 21052503LJ | 15 | 6 |
| RVΔGL-Flpo: 4 weeks | 21052503LJ | total: | 130 |

| | | | |
|---|---|---|---|
| RVΔGL-Flpo: 4 weeks | 21052504LJ | 7 | 2 |
| RVΔGL-Flpo: 4 weeks | 21052504LJ | 8 | 13 |
| RVΔGL-Flpo: 4 weeks | 21052504LJ | 9 | 18 |
| RVΔGL-Flpo: 4 weeks | 21052504LJ | 10 | 16 |
| RVΔGL-Flpo: 4 weeks | 21052504LJ | 11 | 8 |
| RVΔGL-Flpo: 4 weeks | 21052504LJ | 12 | 3 |
| RVΔGL-Flpo: 4 weeks | 21052504LJ | 13 | 3 |
| RVΔGL-Flpo: 4 weeks | 21052504LJ | 14 | 1 |
| RVΔGL-Flpo: 4 weeks | 21052504LJ | 15 | 1 |
| RVΔGL-Flpo: 4 weeks | 21052504LJ | total: | 65 |
| RVΔL-Flpo: 4 weeks | 21021801LJ | 1 | 36 |
| RVΔL-Flpo: 4 weeks | 21021801LJ | 2 | 18 |
| RVΔL-Flpo: 4 weeks | 21021801LJ | 3 | 2 |
| RVΔL-Flpo: 4 weeks | 21021801LJ | 4 | 28 |
| RVΔL-Flpo: 4 weeks | 21021801LJ | 5 | 180 |
| RVΔL-Flpo: 4 weeks | 21021801LJ | 6 | 415 |
| RVΔL-Flpo: 4 weeks | 21021801LJ | 7 | 622 |
| RVΔL-Flpo: 4 weeks | 21021801LJ | 8 | 649 |
| RVΔL-Flpo: 4 weeks | 21021801LJ | 9 | 739 |
| RVΔL-Flpo: 4 weeks | 21021801LJ | 10 | 577 |
| RVΔL-Flpo: 4 weeks | 21021801LJ | 11 | 431 |
| RVΔL-Flpo: 4 weeks | 21021801LJ | 12 | 211 |
| RVΔL-Flpo: 4 weeks | 21021801LJ | 13 | 166 |
| RVΔL-Flpo: 4 weeks | 21021801LJ | 14 | 170 |
| RVΔL-Flpo: 4 weeks | 21021801LJ | 15 | 370 |
| RVΔL-Flpo: 4 weeks | 21021801LJ | total: | 4614 |
| RVΔL-Flpo: 4 weeks | 21021802LJ | 1 | 3 |
| RVΔL-Flpo: 4 weeks | 21021802LJ | 2 | 6 |
| RVΔL-Flpo: 4 weeks | 21021802LJ | 3 | 8 |
| RVΔL-Flpo: 4 weeks | 21021802LJ | 5 | 32 |
| RVΔL-Flpo: 4 weeks | 21021802LJ | 6 | 224 |
| RVΔL-Flpo: 4 weeks | 21021802LJ | 7 | 279 |
| RVΔL-Flpo: 4 weeks | 21021802LJ | 8 | 287 |
| RVΔL-Flpo: 4 weeks | 21021802LJ | 9 | 457 |
| RVΔL-Flpo: 4 weeks | 21021802LJ | 10 | 501 |
| RVΔL-Flpo: 4 weeks | 21021802LJ | 11 | 491 |
| RVΔL-Flpo: 4 weeks | 21021802LJ | 12 | 211 |
| RVΔL-Flpo: 4 weeks | 21021802LJ | 13 | 287 |
| RVΔL-Flpo: 4 weeks | 21021802LJ | 14 | 220 |
| RVΔL-Flpo: 4 weeks | 21021802LJ | total: | 3006 |
| RVΔL-Flpo: 4 weeks | 21021803LJ | 2 | 6 |
| RVΔL-Flpo: 4 weeks | 21021803LJ | 3 | 9 |
| RVΔL-Flpo: 4 weeks | 21021803LJ | 4 | 48 |
| RVΔL-Flpo: 4 weeks | 21021803LJ | 5 | 177 |
| RVΔL-Flpo: 4 weeks | 21021803LJ | 6 | 247 |
| RVΔL-Flpo: 4 weeks | 21021803LJ | 7 | 339 |
| RVΔL-Flpo: 4 weeks | 21021803LJ | 8 | 316 |
| RVΔL-Flpo: 4 weeks | 21021803LJ | 9 | 322 |
| RVΔL-Flpo: 4 weeks | 21021803LJ | 10 | 205 |
| RVΔL-Flpo: 4 weeks | 21021803LJ | 11 | 78 |
| RVΔL-Flpo: 4 weeks | 21021803LJ | 12 | 46 |
| RVΔL-Flpo: 4 weeks | 21021803LJ | 13 | 41 |
| RVΔL-Flpo: 4 weeks | 21021803LJ | 14 | 13 |
| RVΔL-Flpo: 4 weeks | 21021803LJ | total: | 1847 |
| RVΔL-Flpo: 4 weeks | 21021804LJ | 1 | 6 |

| | | | |
|---|---|---|---|
| RVΔL-Flpo: 4 weeks | 21021804LJ | 2 | 14 |
| RVΔL-Flpo: 4 weeks | 21021804LJ | 3 | 2 |
| RVΔL-Flpo: 4 weeks | 21021804LJ | 4 | 57 |
| RVΔL-Flpo: 4 weeks | 21021804LJ | 5 | 359 |
| RVΔL-Flpo: 4 weeks | 21021804LJ | 6 | 414 |
| RVΔL-Flpo: 4 weeks | 21021804LJ | 7 | 469 |
| RVΔL-Flpo: 4 weeks | 21021804LJ | 8 | 576 |
| RVΔL-Flpo: 4 weeks | 21021804LJ | 9 | 533 |
| RVΔL-Flpo: 4 weeks | 21021804LJ | 10 | 383 |
| RVΔL-Flpo: 4 weeks | 21021804LJ | 11 | 283 |
| RVΔL-Flpo: 4 weeks | 21021804LJ | 12 | 114 |
| RVΔL-Flpo: 4 weeks | 21021804LJ | 13 | 34 |
| RVΔL-Flpo: 4 weeks | 21021804LJ | 14 | 87 |
| RVΔL-Flpo: 4 weeks | 21021804LJ | total: | 3331 |
| RVΔL-Flpo: 4 weeks | 21052601LJ | 2 | 1 |
| RVΔL-Flpo: 4 weeks | 21052601LJ | 3 | 1 |
| RVΔL-Flpo: 4 weeks | 21052601LJ | 5 | 16 |
| RVΔL-Flpo: 4 weeks | 21052601LJ | 6 | 10 |
| RVΔL-Flpo: 4 weeks | 21052601LJ | 7 | 28 |
| RVΔL-Flpo: 4 weeks | 21052601LJ | 8 | 92 |
| RVΔL-Flpo: 4 weeks | 21052601LJ | 9 | 161 |
| RVΔL-Flpo: 4 weeks | 21052601LJ | 10 | 232 |
| RVΔL-Flpo: 4 weeks | 21052601LJ | 11 | 280 |
| RVΔL-Flpo: 4 weeks | 21052601LJ | 12 | 201 |
| RVΔL-Flpo: 4 weeks | 21052601LJ | 13 | 143 |
| RVΔL-Flpo: 4 weeks | 21052601LJ | 14 | 106 |
| RVΔL-Flpo: 4 weeks | 21052601LJ | 15 | 98 |
| RVΔL-Flpo: 4 weeks | 21052601LJ | total: | 1369 |
| RVΔL-Flpo: 4 weeks | 21052602LJ | 1 | 49 |
| RVΔL-Flpo: 4 weeks | 21052602LJ | 2 | 12 |
| RVΔL-Flpo: 4 weeks | 21052602LJ | 3 | 17 |
| RVΔL-Flpo: 4 weeks | 21052602LJ | 4 | 51 |
| RVΔL-Flpo: 4 weeks | 21052602LJ | 5 | 163 |
| RVΔL-Flpo: 4 weeks | 21052602LJ | 6 | 418 |
| RVΔL-Flpo: 4 weeks | 21052602LJ | 7 | 489 |
| RVΔL-Flpo: 4 weeks | 21052602LJ | 8 | 526 |
| RVΔL-Flpo: 4 weeks | 21052602LJ | 9 | 510 |
| RVΔL-Flpo: 4 weeks | 21052602LJ | 10 | 319 |
| RVΔL-Flpo: 4 weeks | 21052602LJ | 11 | 179 |
| RVΔL-Flpo: 4 weeks | 21052602LJ | 12 | 72 |
| RVΔL-Flpo: 4 weeks | 21052602LJ | 13 | 48 |
| RVΔL-Flpo: 4 weeks | 21052602LJ | 14 | 32 |
| RVΔL-Flpo: 4 weeks | 21052602LJ | 15 | 34 |
| RVΔL-Flpo: 4 weeks | 21052602LJ | total: | 2919 |
| RVΔL-Flpo: 4 weeks | 21052603LJ | 1 | 38 |
| RVΔL-Flpo: 4 weeks | 21052603LJ | 2 | 33 |
| RVΔL-Flpo: 4 weeks | 21052603LJ | 3 | 16 |
| RVΔL-Flpo: 4 weeks | 21052603LJ | 4 | 44 |
| RVΔL-Flpo: 4 weeks | 21052603LJ | 5 | 302 |
| RVΔL-Flpo: 4 weeks | 21052603LJ | 6 | 648 |
| RVΔL-Flpo: 4 weeks | 21052603LJ | 7 | 636 |
| RVΔL-Flpo: 4 weeks | 21052603LJ | 8 | 666 |
| RVΔL-Flpo: 4 weeks | 21052603LJ | 9 | 582 |
| RVΔL-Flpo: 4 weeks | 21052603LJ | 10 | 409 |
| RVΔL-Flpo: 4 weeks | 21052603LJ | 11 | 207 |

| | | | |
|---|---|---|---|
| RVΔL-Flpo: 4 weeks | 21052603LJ | 12 | 79 |
| RVΔL-Flpo: 4 weeks | 21052603LJ | 13 | 26 |
| RVΔL-Flpo: 4 weeks | 21052603LJ | 14 | 25 |
| RVΔL-Flpo: 4 weeks | 21052603LJ | 15 | 19 |
| RVΔL-Flpo: 4 weeks | 21052603LJ | total: | 3730 |
| RVΔL-Flpo: 4 weeks | 21052606LJ | 1 | 43 |
| RVΔL-Flpo: 4 weeks | 21052606LJ | 2 | 23 |
| RVΔL-Flpo: 4 weeks | 21052606LJ | 3 | 14 |
| RVΔL-Flpo: 4 weeks | 21052606LJ | 4 | 32 |
| RVΔL-Flpo: 4 weeks | 21052606LJ | 5 | 220 |
| RVΔL-Flpo: 4 weeks | 21052606LJ | 6 | 503 |
| RVΔL-Flpo: 4 weeks | 21052606LJ | 7 | 616 |
| RVΔL-Flpo: 4 weeks | 21052606LJ | 8 | 622 |
| RVΔL-Flpo: 4 weeks | 21052606LJ | 9 | 560 |
| RVΔL-Flpo: 4 weeks | 21052606LJ | 10 | 403 |
| RVΔL-Flpo: 4 weeks | 21052606LJ | 11 | 179 |
| RVΔL-Flpo: 4 weeks | 21052606LJ | 12 | 82 |
| RVΔL-Flpo: 4 weeks | 21052606LJ | 13 | 56 |
| RVΔL-Flpo: 4 weeks | 21052606LJ | 14 | 68 |
| RVΔL-Flpo: 4 weeks | 21052606LJ | 15 | 33 |
| RVΔL-Flpo: 4 weeks | 21052606LJ | total: | 3454 |
| RVΔGL-Cre: 1 week | 21021703LJ | 1 | 422 |
| RVΔGL-Cre: 1 week | 21021703LJ | 2 | 395 |
| RVΔGL-Cre: 1 week | 21021703LJ | 3 | 606 |
| RVΔGL-Cre: 1 week | 21021703LJ | 4 | 543 |
| RVΔGL-Cre: 1 week | 21021703LJ | 5 | 637 |
| RVΔGL-Cre: 1 week | 21021703LJ | 6 | 848 |
| RVΔGL-Cre: 1 week | 21021703LJ | 7 | 1145 |
| RVΔGL-Cre: 1 week | 21021703LJ | 8 | 1183 |
| RVΔGL-Cre: 1 week | 21021703LJ | 9 | 1292 |
| RVΔGL-Cre: 1 week | 21021703LJ | 10 | 1311 |
| RVΔGL-Cre: 1 week | 21021703LJ | 11 | 1377 |
| RVΔGL-Cre: 1 week | 21021703LJ | 12 | 1251 |
| RVΔGL-Cre: 1 week | 21021703LJ | 13 | 998 |
| RVΔGL-Cre: 1 week | 21021703LJ | 14 | 835 |
| RVΔGL-Cre: 1 week | 21021703LJ | 15 | 641 |
| RVΔGL-Cre: 1 week | 21021703LJ | total: | 13484 |
| RVΔGL-Cre: 1 week | 21021704LJ | 1 | 918 |
| RVΔGL-Cre: 1 week | 21021704LJ | 2 | 975 |
| RVΔGL-Cre: 1 week | 21021704LJ | 3 | 1144 |
| RVΔGL-Cre: 1 week | 21021704LJ | 4 | 1404 |
| RVΔGL-Cre: 1 week | 21021704LJ | 5 | 1491 |
| RVΔGL-Cre: 1 week | 21021704LJ | 6 | 1377 |
| RVΔGL-Cre: 1 week | 21021704LJ | 7 | 1231 |
| RVΔGL-Cre: 1 week | 21021704LJ | 8 | 1337 |
| RVΔGL-Cre: 1 week | 21021704LJ | 9 | 1135 |
| RVΔGL-Cre: 1 week | 21021704LJ | 10 | 1058 |
| RVΔGL-Cre: 1 week | 21021704LJ | 11 | 711 |
| RVΔGL-Cre: 1 week | 21021704LJ | 12 | 568 |
| RVΔGL-Cre: 1 week | 21021704LJ | 13 | 443 |
| RVΔGL-Cre: 1 week | 21021704LJ | 14 | 432 |
| RVΔGL-Cre: 1 week | 21021704LJ | 15 | 411 |
| RVΔGL-Cre: 1 week | 21021704LJ | total: | 14635 |
| RVΔGL-Cre: 1 week | 21021707LJ | 1 | 497 |
| RVΔGL-Cre: 1 week | 21021707LJ | 2 | 568 |

| | | | |
|---|---|---|---|
| RVΔGL-Cre: 1 week | 21021707LJ | 3 | 454 |
| RVΔGL-Cre: 1 week | 21021707LJ | 4 | 454 |
| RVΔGL-Cre: 1 week | 21021707LJ | 5 | 583 |
| RVΔGL-Cre: 1 week | 21021707LJ | 6 | 873 |
| RVΔGL-Cre: 1 week | 21021707LJ | 7 | 973 |
| RVΔGL-Cre: 1 week | 21021707LJ | 8 | 1133 |
| RVΔGL-Cre: 1 week | 21021707LJ | 9 | 1220 |
| RVΔGL-Cre: 1 week | 21021707LJ | 10 | 1176 |
| RVΔGL-Cre: 1 week | 21021707LJ | 11 | 994 |
| RVΔGL-Cre: 1 week | 21021707LJ | 12 | 645 |
| RVΔGL-Cre: 1 week | 21021707LJ | 13 | 467 |
| RVΔGL-Cre: 1 week | 21021707LJ | 14 | 330 |
| RVΔGL-Cre: 1 week | 21021707LJ | 15 | 280 |
| RVΔGL-Cre: 1 week | 21021707LJ | total: | 10647 |
| RVΔGL-Cre: 1 week | 21021708LJ | 1 | 271 |
| RVΔGL-Cre: 1 week | 21021708LJ | 2 | 303 |
| RVΔGL-Cre: 1 week | 21021708LJ | 3 | 386 |
| RVΔGL-Cre: 1 week | 21021708LJ | 4 | 634 |
| RVΔGL-Cre: 1 week | 21021708LJ | 5 | 805 |
| RVΔGL-Cre: 1 week | 21021708LJ | 6 | 931 |
| RVΔGL-Cre: 1 week | 21021708LJ | 7 | 1029 |
| RVΔGL-Cre: 1 week | 21021708LJ | 8 | 1186 |
| RVΔGL-Cre: 1 week | 21021708LJ | 9 | 1227 |
| RVΔGL-Cre: 1 week | 21021708LJ | 10 | 1102 |
| RVΔGL-Cre: 1 week | 21021708LJ | 11 | 1019 |
| RVΔGL-Cre: 1 week | 21021708LJ | 12 | 666 |
| RVΔGL-Cre: 1 week | 21021708LJ | 13 | 431 |
| RVΔGL-Cre: 1 week | 21021708LJ | 14 | 369 |
| RVΔGL-Cre: 1 week | 21021708LJ | 15 | 486 |
| RVΔGL-Cre: 1 week | 21021708LJ | total: | 10845 |
| RVΔL-Cre: 1 week | 21021905LJ | 1 | 1161 |
| RVΔL-Cre: 1 week | 21021905LJ | 2 | 1093 |
| RVΔL-Cre: 1 week | 21021905LJ | 3 | 1072 |
| RVΔL-Cre: 1 week | 21021905LJ | 4 | 1079 |
| RVΔL-Cre: 1 week | 21021905LJ | 5 | 1318 |
| RVΔL-Cre: 1 week | 21021905LJ | 6 | 1417 |
| RVΔL-Cre: 1 week | 21021905LJ | 7 | 1541 |
| RVΔL-Cre: 1 week | 21021905LJ | 8 | 1510 |
| RVΔL-Cre: 1 week | 21021905LJ | 9 | 1500 |
| RVΔL-Cre: 1 week | 21021905LJ | 10 | 1549 |
| RVΔL-Cre: 1 week | 21021905LJ | 11 | 1381 |
| RVΔL-Cre: 1 week | 21021905LJ | 12 | 1109 |
| RVΔL-Cre: 1 week | 21021905LJ | 13 | 532 |
| RVΔL-Cre: 1 week | 21021905LJ | 14 | 380 |
| RVΔL-Cre: 1 week | 21021905LJ | 15 | 265 |
| RVΔL-Cre: 1 week | 21021905LJ | total: | 16907 |
| RVΔL-Cre: 1 week | 21021906LJ | 1 | 985 |
| RVΔL-Cre: 1 week | 21021906LJ | 2 | 1012 |
| RVΔL-Cre: 1 week | 21021906LJ | 3 | 1020 |
| RVΔL-Cre: 1 week | 21021906LJ | 4 | 1060 |
| RVΔL-Cre: 1 week | 21021906LJ | 5 | 1082 |
| RVΔL-Cre: 1 week | 21021906LJ | 6 | 1515 |
| RVΔL-Cre: 1 week | 21021906LJ | 7 | 1808 |
| RVΔL-Cre: 1 week | 21021906LJ | 8 | 1675 |
| RVΔL-Cre: 1 week | 21021906LJ | 9 | 1538 |

| | | | |
|---|---|---|---|
| RVΔL-Cre: 1 week | 21021906LJ | 10 | 1728 |
| RVΔL-Cre: 1 week | 21021906LJ | 11 | 955 |
| RVΔL-Cre: 1 week | 21021906LJ | 12 | 1018 |
| RVΔL-Cre: 1 week | 21021906LJ | 13 | 993 |
| RVΔL-Cre: 1 week | 21021906LJ | 14 | 701 |
| RVΔL-Cre: 1 week | 21021906LJ | 15 | 483 |
| RVΔL-Cre: 1 week | 21021906LJ | 16 | 350 |
| RVΔL-Cre: 1 week | 21021906LJ | total: | 17923 |
| RVΔL-Cre: 1 week | 21021907LJ | 1 | 1207 |
| RVΔL-Cre: 1 week | 21021907LJ | 2 | 1083 |
| RVΔL-Cre: 1 week | 21021907LJ | 3 | 1152 |
| RVΔL-Cre: 1 week | 21021907LJ | 4 | 1225 |
| RVΔL-Cre: 1 week | 21021907LJ | 5 | 1336 |
| RVΔL-Cre: 1 week | 21021907LJ | 6 | 1558 |
| RVΔL-Cre: 1 week | 21021907LJ | 7 | 1498 |
| RVΔL-Cre: 1 week | 21021907LJ | 8 | 1709 |
| RVΔL-Cre: 1 week | 21021907LJ | 9 | 1547 |
| RVΔL-Cre: 1 week | 21021907LJ | 10 | 1718 |
| RVΔL-Cre: 1 week | 21021907LJ | 11 | 1503 |
| RVΔL-Cre: 1 week | 21021907LJ | 12 | 1249 |
| RVΔL-Cre: 1 week | 21021907LJ | 13 | 845 |
| RVΔL-Cre: 1 week | 21021907LJ | 14 | 626 |
| RVΔL-Cre: 1 week | 21021907LJ | 15 | 495 |
| RVΔL-Cre: 1 week | 21021907LJ | total: | 18751 |
| RVΔL-Cre: 1 week | 21021908LJ | 1 | 1011 |
| RVΔL-Cre: 1 week | 21021908LJ | 2 | 892 |
| RVΔL-Cre: 1 week | 21021908LJ | 3 | 896 |
| RVΔL-Cre: 1 week | 21021908LJ | 4 | 957 |
| RVΔL-Cre: 1 week | 21021908LJ | 5 | 1243 |
| RVΔL-Cre: 1 week | 21021908LJ | 6 | 1448 |
| RVΔL-Cre: 1 week | 21021908LJ | 7 | 1568 |
| RVΔL-Cre: 1 week | 21021908LJ | 8 | 1459 |
| RVΔL-Cre: 1 week | 21021908LJ | 9 | 1573 |
| RVΔL-Cre: 1 week | 21021908LJ | 10 | 1313 |
| RVΔL-Cre: 1 week | 21021908LJ | 11 | 1146 |
| RVΔL-Cre: 1 week | 21021908LJ | 12 | 1046 |
| RVΔL-Cre: 1 week | 21021908LJ | 13 | 774 |
| RVΔL-Cre: 1 week | 21021908LJ | 14 | 483 |
| RVΔL-Cre: 1 week | 21021908LJ | 15 | 448 |
| RVΔL-Cre: 1 week | 21021908LJ | total: | 16257 |
| RVΔGL-Cre: 4 weeks | 21021701LJ | 1 | 1314 |
| RVΔGL-Cre: 4 weeks | 21021701LJ | 2 | 1246 |
| RVΔGL-Cre: 4 weeks | 21021701LJ | 3 | 1351 |
| RVΔGL-Cre: 4 weeks | 21021701LJ | 4 | 1601 |
| RVΔGL-Cre: 4 weeks | 21021701LJ | 5 | 1668 |
| RVΔGL-Cre: 4 weeks | 21021701LJ | 6 | 1618 |
| RVΔGL-Cre: 4 weeks | 21021701LJ | 7 | 1593 |
| RVΔGL-Cre: 4 weeks | 21021701LJ | 8 | 1757 |
| RVΔGL-Cre: 4 weeks | 21021701LJ | 9 | 1730 |
| RVΔGL-Cre: 4 weeks | 21021701LJ | 10 | 1600 |
| RVΔGL-Cre: 4 weeks | 21021701LJ | 11 | 1268 |
| RVΔGL-Cre: 4 weeks | 21021701LJ | 12 | 931 |
| RVΔGL-Cre: 4 weeks | 21021701LJ | 13 | 652 |
| RVΔGL-Cre: 4 weeks | 21021701LJ | 14 | 491 |
| RVΔGL-Cre: 4 weeks | 21021701LJ | 15 | 336 |

| | | | |
|---|---|---|---|
| RVΔGL-Cre: 4 weeks | 21021701LJ | total: | 19156 |
| RVΔGL-Cre: 4 weeks | 21021702LJ | 1 | 957 |
| RVΔGL-Cre: 4 weeks | 21021702LJ | 2 | 1175 |
| RVΔGL-Cre: 4 weeks | 21021702LJ | 3 | 1414 |
| RVΔGL-Cre: 4 weeks | 21021702LJ | 4 | 1470 |
| RVΔGL-Cre: 4 weeks | 21021702LJ | 5 | 1481 |
| RVΔGL-Cre: 4 weeks | 21021702LJ | 6 | 1654 |
| RVΔGL-Cre: 4 weeks | 21021702LJ | 7 | 1616 |
| RVΔGL-Cre: 4 weeks | 21021702LJ | 8 | 1601 |
| RVΔGL-Cre: 4 weeks | 21021702LJ | 9 | 1717 |
| RVΔGL-Cre: 4 weeks | 21021702LJ | 10 | 1449 |
| RVΔGL-Cre: 4 weeks | 21021702LJ | 11 | 1132 |
| RVΔGL-Cre: 4 weeks | 21021702LJ | 12 | 871 |
| RVΔGL-Cre: 4 weeks | 21021702LJ | 13 | 587 |
| RVΔGL-Cre: 4 weeks | 21021702LJ | 14 | 456 |
| RVΔGL-Cre: 4 weeks | 21021702LJ | 15 | 355 |
| RVΔGL-Cre: 4 weeks | 21021702LJ | total: | 17935 |
| RVΔGL-Cre: 4 weeks | 21021705LJ | 1 | 892 |
| RVΔGL-Cre: 4 weeks | 21021705LJ | 2 | 824 |
| RVΔGL-Cre: 4 weeks | 21021705LJ | 3 | 879 |
| RVΔGL-Cre: 4 weeks | 21021705LJ | 4 | 1094 |
| RVΔGL-Cre: 4 weeks | 21021705LJ | 5 | 1182 |
| RVΔGL-Cre: 4 weeks | 21021705LJ | 6 | 1318 |
| RVΔGL-Cre: 4 weeks | 21021705LJ | 7 | 1482 |
| RVΔGL-Cre: 4 weeks | 21021705LJ | 8 | 1637 |
| RVΔGL-Cre: 4 weeks | 21021705LJ | 9 | 1666 |
| RVΔGL-Cre: 4 weeks | 21021705LJ | 10 | 1796 |
| RVΔGL-Cre: 4 weeks | 21021705LJ | 11 | 1636 |
| RVΔGL-Cre: 4 weeks | 21021705LJ | 12 | 1415 |
| RVΔGL-Cre: 4 weeks | 21021705LJ | 13 | 1121 |
| RVΔGL-Cre: 4 weeks | 21021705LJ | 14 | 894 |
| RVΔGL-Cre: 4 weeks | 21021705LJ | 15 | 703 |
| RVΔGL-Cre: 4 weeks | 21021705LJ | total: | 18539 |
| RVΔGL-Cre: 4 weeks | 21021706LJ | 1 | 756 |
| RVΔGL-Cre: 4 weeks | 21021706LJ | 2 | 817 |
| RVΔGL-Cre: 4 weeks | 21021706LJ | 3 | 750 |
| RVΔGL-Cre: 4 weeks | 21021706LJ | 4 | 798 |
| RVΔGL-Cre: 4 weeks | 21021706LJ | 5 | 920 |
| RVΔGL-Cre: 4 weeks | 21021706LJ | 6 | 1337 |
| RVΔGL-Cre: 4 weeks | 21021706LJ | 7 | 1485 |
| RVΔGL-Cre: 4 weeks | 21021706LJ | 8 | 1490 |
| RVΔGL-Cre: 4 weeks | 21021706LJ | 9 | 1641 |
| RVΔGL-Cre: 4 weeks | 21021706LJ | 10 | 2004 |
| RVΔGL-Cre: 4 weeks | 21021706LJ | 11 | 1955 |
| RVΔGL-Cre: 4 weeks | 21021706LJ | 12 | 1453 |
| RVΔGL-Cre: 4 weeks | 21021706LJ | 13 | 1122 |
| RVΔGL-Cre: 4 weeks | 21021706LJ | 14 | 769 |
| RVΔGL-Cre: 4 weeks | 21021706LJ | 15 | 641 |
| RVΔGL-Cre: 4 weeks | 21021706LJ | total: | 17938 |
| RVΔL-Cre: 4 weeks | 21021901LJ | 1 | 943 |
| RVΔL-Cre: 4 weeks | 21021901LJ | 2 | 1003 |
| RVΔL-Cre: 4 weeks | 21021901LJ | 3 | 1036 |
| RVΔL-Cre: 4 weeks | 21021901LJ | 4 | 1195 |
| RVΔL-Cre: 4 weeks | 21021901LJ | 5 | 1402 |
| RVΔL-Cre: 4 weeks | 21021901LJ | 6 | 1739 |

| | | | |
|---|---|---|---|
| RVΔL-Cre: 4 weeks | 21021901LJ | 7 | 1793 |
| RVΔL-Cre: 4 weeks | 21021901LJ | 8 | 1913 |
| RVΔL-Cre: 4 weeks | 21021901LJ | 9 | 2026 |
| RVΔL-Cre: 4 weeks | 21021901LJ | 10 | 1913 |
| RVΔL-Cre: 4 weeks | 21021901LJ | 11 | 1929 |
| RVΔL-Cre: 4 weeks | 21021901LJ | 12 | 1668 |
| RVΔL-Cre: 4 weeks | 21021901LJ | 13 | 1247 |
| RVΔL-Cre: 4 weeks | 21021901LJ | 14 | 1062 |
| RVΔL-Cre: 4 weeks | 21021901LJ | 15 | 874 |
| RVΔL-Cre: 4 weeks | 21021901LJ | total: | 21743 |
| RVΔL-Cre: 4 weeks | 21021902LJ | 1 | 1357 |
| RVΔL-Cre: 4 weeks | 21021902LJ | 2 | 1416 |
| RVΔL-Cre: 4 weeks | 21021902LJ | 3 | 1287 |
| RVΔL-Cre: 4 weeks | 21021902LJ | 4 | 1308 |
| RVΔL-Cre: 4 weeks | 21021902LJ | 5 | 1583 |
| RVΔL-Cre: 4 weeks | 21021902LJ | 6 | 1850 |
| RVΔL-Cre: 4 weeks | 21021902LJ | 7 | 2020 |
| RVΔL-Cre: 4 weeks | 21021902LJ | 8 | 2238 |
| RVΔL-Cre: 4 weeks | 21021902LJ | 9 | 2092 |
| RVΔL-Cre: 4 weeks | 21021902LJ | 10 | 2321 |
| RVΔL-Cre: 4 weeks | 21021902LJ | 11 | 2241 |
| RVΔL-Cre: 4 weeks | 21021902LJ | 12 | 1858 |
| RVΔL-Cre: 4 weeks | 21021902LJ | 13 | 1326 |
| RVΔL-Cre: 4 weeks | 21021902LJ | 14 | 1083 |
| RVΔL-Cre: 4 weeks | 21021902LJ | 15 | 793 |
| RVΔL-Cre: 4 weeks | 21021902LJ | total: | 24773 |
| RVΔL-Cre: 4 weeks | 21021903LJ | 1 | 1543 |
| RVΔL-Cre: 4 weeks | 21021903LJ | 2 | 1406 |
| RVΔL-Cre: 4 weeks | 21021903LJ | 3 | 1303 |
| RVΔL-Cre: 4 weeks | 21021903LJ | 4 | 1489 |
| RVΔL-Cre: 4 weeks | 21021903LJ | 5 | 1825 |
| RVΔL-Cre: 4 weeks | 21021903LJ | 6 | 1946 |
| RVΔL-Cre: 4 weeks | 21021903LJ | 7 | 1924 |
| RVΔL-Cre: 4 weeks | 21021903LJ | 8 | 2142 |
| RVΔL-Cre: 4 weeks | 21021903LJ | 9 | 2077 |
| RVΔL-Cre: 4 weeks | 21021903LJ | 10 | 1954 |
| RVΔL-Cre: 4 weeks | 21021903LJ | 11 | 1672 |
| RVΔL-Cre: 4 weeks | 21021903LJ | 12 | 1411 |
| RVΔL-Cre: 4 weeks | 21021903LJ | 13 | 1020 |
| RVΔL-Cre: 4 weeks | 21021903LJ | 14 | 687 |
| RVΔL-Cre: 4 weeks | 21021903LJ | 15 | 627 |
| RVΔL-Cre: 4 weeks | 21021903LJ | total: | 23026 |
| RVΔL-Cre: 4 weeks | 21021904LJ | 1 | 1592 |
| RVΔL-Cre: 4 weeks | 21021904LJ | 2 | 1466 |
| RVΔL-Cre: 4 weeks | 21021904LJ | 3 | 1355 |
| RVΔL-Cre: 4 weeks | 21021904LJ | 4 | 1599 |
| RVΔL-Cre: 4 weeks | 21021904LJ | 5 | 1736 |
| RVΔL-Cre: 4 weeks | 21021904LJ | 6 | 1862 |
| RVΔL-Cre: 4 weeks | 21021904LJ | 7 | 1794 |
| RVΔL-Cre: 4 weeks | 21021904LJ | 8 | 1930 |
| RVΔL-Cre: 4 weeks | 21021904LJ | 9 | 2047 |
| RVΔL-Cre: 4 weeks | 21021904LJ | 10 | 1584 |
| RVΔL-Cre: 4 weeks | 21021904LJ | 11 | 1755 |
| RVΔL-Cre: 4 weeks | 21021904LJ | 12 | 1431 |
| RVΔL-Cre: 4 weeks | 21021904LJ | 13 | 741 |

| | | | |
|---|---|---|---|
| RVΔL-Cre: 4 weeks | 21021904LJ | 14 | 755 |
| RVΔL-Cre: 4 weeks | 21021904LJ | 15 | 625 |
| RVΔL-Cre: 4 weeks | 21021904LJ | total: | 22272 |
| RVΔL-5tTA: 1 week | 21042901LJ | 3 | 2 |
| RVΔL-5tTA: 1 week | 21042901LJ | 6 | 9 |
| RVΔL-5tTA: 1 week | 21042901LJ | 7 | 4 |
| RVΔL-5tTA: 1 week | 21042901LJ | 8 | 26 |
| RVΔL-5tTA: 1 week | 21042901LJ | 9 | 32 |
| RVΔL-5tTA: 1 week | 21042901LJ | 10 | 54 |
| RVΔL-5tTA: 1 week | 21042901LJ | 11 | 69 |
| RVΔL-5tTA: 1 week | 21042901LJ | 12 | 56 |
| RVΔL-5tTA: 1 week | 21042901LJ | 13 | 51 |
| RVΔL-5tTA: 1 week | 21042901LJ | 14 | 14 |
| RVΔL-5tTA: 1 week | 21042901LJ | 15 | 22 |
| RVΔL-5tTA: 1 week | 21042901LJ | total: | 339 |
| RVΔL-5tTA: 1 week | 21042902LJ | 1 | 4 |
| RVΔL-5tTA: 1 week | 21042902LJ | 2 | 11 |
| RVΔL-5tTA: 1 week | 21042902LJ | 3 | 14 |
| RVΔL-5tTA: 1 week | 21042902LJ | 4 | 4 |
| RVΔL-5tTA: 1 week | 21042902LJ | 5 | 39 |
| RVΔL-5tTA: 1 week | 21042902LJ | 6 | 108 |
| RVΔL-5tTA: 1 week | 21042902LJ | 7 | 213 |
| RVΔL-5tTA: 1 week | 21042902LJ | 8 | 255 |
| RVΔL-5tTA: 1 week | 21042902LJ | 9 | 296 |
| RVΔL-5tTA: 1 week | 21042902LJ | 10 | 219 |
| RVΔL-5tTA: 1 week | 21042902LJ | 11 | 134 |
| RVΔL-5tTA: 1 week | 21042902LJ | 12 | 64 |
| RVΔL-5tTA: 1 week | 21042902LJ | 13 | 29 |
| RVΔL-5tTA: 1 week | 21042902LJ | 14 | 14 |
| RVΔL-5tTA: 1 week | 21042902LJ | 15 | 7 |
| RVΔL-5tTA: 1 week | 21042902LJ | total: | 1411 |
| RVΔL-5tTA: 1 week | 21042903LJ | 2 | 3 |
| RVΔL-5tTA: 1 week | 21042903LJ | 3 | 7 |
| RVΔL-5tTA: 1 week | 21042903LJ | 4 | 6 |
| RVΔL-5tTA: 1 week | 21042903LJ | 5 | 4 |
| RVΔL-5tTA: 1 week | 21042903LJ | 6 | 27 |
| RVΔL-5tTA: 1 week | 21042903LJ | 7 | 40 |
| RVΔL-5tTA: 1 week | 21042903LJ | 8 | 81 |
| RVΔL-5tTA: 1 week | 21042903LJ | 9 | 115 |
| RVΔL-5tTA: 1 week | 21042903LJ | 10 | 165 |
| RVΔL-5tTA: 1 week | 21042903LJ | 11 | 120 |
| RVΔL-5tTA: 1 week | 21042903LJ | 12 | 84 |
| RVΔL-5tTA: 1 week | 21042903LJ | 13 | 67 |
| RVΔL-5tTA: 1 week | 21042903LJ | 14 | 44 |
| RVΔL-5tTA: 1 week | 21042903LJ | 15 | 33 |
| RVΔL-5tTA: 1 week | 21042903LJ | total: | 796 |
| RVΔL-5tTA: 1 week | 21060301LJ | 1 | 85 |
| RVΔL-5tTA: 1 week | 21060301LJ | 2 | 117 |
| RVΔL-5tTA: 1 week | 21060301LJ | 3 | 154 |
| RVΔL-5tTA: 1 week | 21060301LJ | 4 | 59 |
| RVΔL-5tTA: 1 week | 21060301LJ | 5 | 207 |
| RVΔL-5tTA: 1 week | 21060301LJ | 6 | 325 |
| RVΔL-5tTA: 1 week | 21060301LJ | 7 | 314 |
| RVΔL-5tTA: 1 week | 21060301LJ | 8 | 303 |
| RVΔL-5tTA: 1 week | 21060301LJ | 9 | 288 |

| | | | |
|---|---|---|---|
| RVΔL-5tTA: 1 week | 21060301LJ | 10 | 185 |
| RVΔL-5tTA: 1 week | 21060301LJ | 11 | 141 |
| RVΔL-5tTA: 1 week | 21060301LJ | 12 | 47 |
| RVΔL-5tTA: 1 week | 21060301LJ | 13 | 19 |
| RVΔL-5tTA: 1 week | 21060301LJ | 14 | 14 |
| RVΔL-5tTA: 1 week | 21060301LJ | 15 | 13 |
| RVΔL-5tTA: 1 week | 21060301LJ | total: | 2271 |
| RVΔL-5tTA: 4 weeks | 21042905LJ | 1 | 4 |
| RVΔL-5tTA: 4 weeks | 21042905LJ | 2 | 14 |
| RVΔL-5tTA: 4 weeks | 21042905LJ | 3 | 6 |
| RVΔL-5tTA: 4 weeks | 21042905LJ | 4 | 21 |
| RVΔL-5tTA: 4 weeks | 21042905LJ | 5 | 15 |
| RVΔL-5tTA: 4 weeks | 21042905LJ | 6 | 41 |
| RVΔL-5tTA: 4 weeks | 21042905LJ | 7 | 42 |
| RVΔL-5tTA: 4 weeks | 21042905LJ | 8 | 101 |
| RVΔL-5tTA: 4 weeks | 21042905LJ | 9 | 156 |
| RVΔL-5tTA: 4 weeks | 21042905LJ | 10 | 109 |
| RVΔL-5tTA: 4 weeks | 21042905LJ | 11 | 94 |
| RVΔL-5tTA: 4 weeks | 21042905LJ | 12 | 24 |
| RVΔL-5tTA: 4 weeks | 21042905LJ | 13 | 87 |
| RVΔL-5tTA: 4 weeks | 21042905LJ | 14 | 44 |
| RVΔL-5tTA: 4 weeks | 21042905LJ | 15 | 71 |
| RVΔL-5tTA: 4 weeks | 21042905LJ | total: | 829 |
| RVΔL-5tTA: 4 weeks | 21042906LJ | 1 | 9 |
| RVΔL-5tTA: 4 weeks | 21042906LJ | 2 | 17 |
| RVΔL-5tTA: 4 weeks | 21042906LJ | 3 | 28 |
| RVΔL-5tTA: 4 weeks | 21042906LJ | 4 | 23 |
| RVΔL-5tTA: 4 weeks | 21042906LJ | 5 | 24 |
| RVΔL-5tTA: 4 weeks | 21042906LJ | 6 | 25 |
| RVΔL-5tTA: 4 weeks | 21042906LJ | 7 | 93 |
| RVΔL-5tTA: 4 weeks | 21042906LJ | 8 | 200 |
| RVΔL-5tTA: 4 weeks | 21042906LJ | 9 | 305 |
| RVΔL-5tTA: 4 weeks | 21042906LJ | 10 | 233 |
| RVΔL-5tTA: 4 weeks | 21042906LJ | 11 | 200 |
| RVΔL-5tTA: 4 weeks | 21042906LJ | 12 | 141 |
| RVΔL-5tTA: 4 weeks | 21042906LJ | 13 | 74 |
| RVΔL-5tTA: 4 weeks | 21042906LJ | 14 | 108 |
| RVΔL-5tTA: 4 weeks | 21042906LJ | 15 | 94 |
| RVΔL-5tTA: 4 weeks | 21042906LJ | total: | 1574 |
| RVΔL-5tTA: 4 weeks | 21042907LJ | 2 | 2 |
| RVΔL-5tTA: 4 weeks | 21042907LJ | 4 | 3 |
| RVΔL-5tTA: 4 weeks | 21042907LJ | 5 | 5 |
| RVΔL-5tTA: 4 weeks | 21042907LJ | 6 | 24 |
| RVΔL-5tTA: 4 weeks | 21042907LJ | 7 | 32 |
| RVΔL-5tTA: 4 weeks | 21042907LJ | 8 | 51 |
| RVΔL-5tTA: 4 weeks | 21042907LJ | 9 | 100 |
| RVΔL-5tTA: 4 weeks | 21042907LJ | 10 | 74 |
| RVΔL-5tTA: 4 weeks | 21042907LJ | 11 | 69 |
| RVΔL-5tTA: 4 weeks | 21042907LJ | 12 | 54 |
| RVΔL-5tTA: 4 weeks | 21042907LJ | 13 | 26 |
| RVΔL-5tTA: 4 weeks | 21042907LJ | 14 | 31 |
| RVΔL-5tTA: 4 weeks | 21042907LJ | 15 | 28 |
| RVΔL-5tTA: 4 weeks | 21042907LJ | total: | 499 |
| RVΔL-5tTA: 4 weeks | 21042908LJ | 1 | 64 |
| RVΔL-5tTA: 4 weeks | 21042908LJ | 2 | 67 |

| | | | |
|---|---|---|---|
| RVΔL-5tTA: 4 weeks | 21042908LJ | 3 | 66 |
| RVΔL-5tTA: 4 weeks | 21042908LJ | 4 | 79 |
| RVΔL-5tTA: 4 weeks | 21042908LJ | 5 | 151 |
| RVΔL-5tTA: 4 weeks | 21042908LJ | 6 | 149 |
| RVΔL-5tTA: 4 weeks | 21042908LJ | 7 | 166 |
| RVΔL-5tTA: 4 weeks | 21042908LJ | 8 | 171 |
| RVΔL-5tTA: 4 weeks | 21042908LJ | 9 | 184 |
| RVΔL-5tTA: 4 weeks | 21042908LJ | 10 | 57 |
| RVΔL-5tTA: 4 weeks | 21042908LJ | 11 | 48 |
| RVΔL-5tTA: 4 weeks | 21042908LJ | 12 | 29 |
| RVΔL-5tTA: 4 weeks | 21042908LJ | 13 | 14 |
| RVΔL-5tTA: 4 weeks | 21042908LJ | 14 | 27 |
| RVΔL-5tTA: 4 weeks | 21042908LJ | 15 | 57 |
| RVΔL-5tTA: 4 weeks | 21042908LJ | total: | 1329 |

| Condition | Mouse number | Cell counts |
|---|---|---|
| RVΔGL-Flpo: 1 week | 21021505LJ | 22 |
| RVΔGL-Flpo: 1 week | 21021506LJ | 18 |
| RVΔGL-Flpo: 1 week | 21021507LJ | 121 |
| RVΔGL-Flpo: 1 week | 21021508LJ | 2 |
| RVΔGL-Flpo: 1 week | 21052505LJ | 7 |
| RVΔGL-Flpo: 1 week | 21052506LJ | 8 |
| RVΔGL-Flpo: 1 week | 21052507LJ | 98 |
| RVΔGL-Flpo: 1 week | 21052508LJ | 6 |
| RVΔGL-Flpo: 1 week | Mean: | 35.25 |
| RVΔL-Flpo: 1 week | 21021805LJ | 3093 |
| RVΔL-Flpo: 1 week | 21021806LJ | 320 |
| RVΔL-Flpo: 1 week | 21021807LJ | 311 |
| RVΔL-Flpo: 1 week | 21021808LJ | 263 |
| RVΔL-Flpo: 1 week | 21052607LJ | 47 |
| RVΔL-Flpo: 1 week | 21052608LJ | 2161 |
| RVΔL-Flpo: 1 week | 21060302LJ | 365 |
| RVΔL-Flpo: 1 week | 21060303LJ | 238 |
| RVΔL-Flpo: 1 week | Mean: | 849.75 |
| RVΔGL-Flpo: 4 weeks | 21021501LJ | 80 |
| RVΔGL-Flpo: 4 weeks | 21021502LJ | 62 |
| RVΔGL-Flpo: 4 weeks | 21021503LJ | 3254 |
| RVΔGL-Flpo: 4 weeks | 21021504LJ | 237 |
| RVΔGL-Flpo: 4 weeks | 21052501LJ | 43 |
| RVΔGL-Flpo: 4 weeks | 21052502LJ | 14 |
| RVΔGL-Flpo: 4 weeks | 21052503LJ | 130 |
| RVΔGL-Flpo: 4 weeks | 21052504LJ | 65 |
| RVΔGL-Flpo: 4 weeks | Mean: | 485.625 |
| RVΔL-Flpo: 4 weeks | 21021801LJ | 4614 |
| RVΔL-Flpo: 4 weeks | 21021802LJ | 3006 |
| RVΔL-Flpo: 4 weeks | 21021803LJ | 1847 |
| RVΔL-Flpo: 4 weeks | 21021804LJ | 3331 |
| RVΔL-Flpo: 4 weeks | 21052601LJ | 1369 |
| RVΔL-Flpo: 4 weeks | 21052602LJ | 2919 |
| RVΔL-Flpo: 4 weeks | 21052603LJ | 3730 |
| RVΔL-Flpo: 4 weeks | 21052606LJ | 3454 |
| RVΔL-Flpo: 4 weeks | Mean: | 3033.75 |
| RVΔGL-Cre: 1 week | 21021703LJ | 13484 |
| RVΔGL-Cre: 1 week | 21021704LJ | 14635 |
| RVΔGL-Cre: 1 week | 21021707LJ | 10647 |
| RVΔGL-Cre: 1 week | 21021708LJ | 10845 |
| RVΔGL-Cre: 1 week | Mean: | 12402.75 |
| RVΔL-Cre: 1 week | 21021905LJ | 16907 |
| RVΔL-Cre: 1 week | 21021906LJ | 17923 |
| RVΔL-Cre: 1 week | 21021907LJ | 18751 |
| RVΔL-Cre: 1 week | 21021908LJ | 16257 |
| RVΔL-Cre: 1 week | Mean: | 17459.5 |
| RVΔGL-Cre: 4 weeks | 21021701LJ | 19156 |
| RVΔGL-Cre: 4 weeks | 21021702LJ | 17935 |
| RVΔGL-Cre: 4 weeks | 21021705LJ | 18539 |
| RVΔGL-Cre: 4 weeks | 21021706LJ | 17938 |
| RVΔGL-Cre: 4 weeks | Mean: | 18392 |
| RVΔL-Cre: 4 weeks | 21021901LJ | 21743 |
| RVΔL-Cre: 4 weeks | 21021902LJ | 24773 |
| RVΔL-Cre: 4 weeks | 21021903LJ | 23026 |
| RVΔL-Cre: 4 weeks | 21021904LJ | 22272 |
| RVΔL-Cre: 4 weeks | Mean: | 22953.5 |
| RVΔL-5tTA: 1 week | 21042901LJ | 339 |
| RVΔL-5tTA: 1 week | 21042902LJ | 1411 |
| RVΔL-5tTA: 1 week | 21042903LJ | 796 |
| RVΔL-5tTA: 1 week | 21060301LJ | 2271 |
| RVΔL-5tTA: 1 week | Mean: | 1204.25 |
| RVΔL-5tTA: 4 weeks | 21042905LJ | 829 |
| RVΔL-5tTA: 4 weeks | 21042906LJ | 1574 |
| RVΔL-5tTA: 4 weeks | 21042907LJ | 499 |
| RVΔL-5tTA: 4 weeks | 21042908LJ | 1329 |
| RVΔL-5tTA: 4 weeks | Mean: | 1057.75 |

| RVΔGL-Flpo 4 weeks | RVΔL-Flpo 4 weeks |
|---|---|
| 80 | 4614 |
| 62 | 3006 |
| 3254 | 1847 |
| 237 | 3331 |
| 43 | 1369 |
| 14 | 2919 |
| 130 | 3730 |
| 65 | 3454 |

Anova: Single Factor

SUMMARY

| *Groups* | *Count* | *Sum* | *Average* | *Variance* |
|---|---|---|---|---|
| RVΔGL-Flpo 4 weeks | 8 | 3885 | 485.625 | 1255920.84 |
| RVΔL-Flpo 4 weeks | 8 | 24270 | 3033.75 | 1062946.79 |

ANOVA

| *Source of Variation* | *SS* | *df* | *MS* | *F* | *P-value* | *F crit* |
|---|---|---|---|---|---|---|
| Between Groups | 25971764.06 | 1 | 25971764.1 | 22.4003852 | 0.00032063 | 4.60010994 |
| Within Groups | 16232073.38 | 14 | 1159433.81 | | | |
| Total | 42203837.44 | 15 | | | | |

| RVΔGL-Flpo 1 week | RVΔGL-Flpo 4 weeks |
|---|---|
| 22 | 80 |
| 18 | 62 |
| 121 | 3254 |
| 2 | 237 |
| 7 | 43 |
| 8 | 14 |
| 98 | 130 |
| 6 | 65 |

Anova: Single Factor

SUMMARY

| *Groups* | *Count* | *Sum* | *Average* | *Variance* |
|---|---|---|---|---|
| RVΔGL-Flpo 1 week | 8 | 282 | 35.25 | 2180.78571 |
| RVΔGL-Flpo 4 weeks | 8 | 3885 | 485.625 | 1255920.84 |

ANOVA

| *Source of Variation* | *SS* | *df* | *MS* | *F* | *P-value* | *F crit* |
|---|---|---|---|---|---|---|
| Between Groups | 811350.5625 | 1 | 811350.563 | 1.28980131 | 0.27515341 | 4.60010994 |
| Within Groups | 8806711.375 | 14 | 629050.813 | | | |
| Total | 9618061.938 | 15 | | | | |

| RVΔGL-Flpo 4 weeks | RVΔL-Flpo 1 week |
|---|---|
| 80 | 3093 |
| 62 | 320 |
| 3254 | 311 |
| 237 | 263 |

Anova: Single Factor

SUMMARY

| *Groups* | *Count* | *Sum* | *Average* | *Variance* |
|---|---|---|---|---|
| RVΔGL-Flpo 4 weeks | 4 | 3633 | 908.25 | 2451752.25 |
| RVΔL-Flpo 1 week | 4 | 3987 | 996.75 | 1953632.25 |

ANOVA

| *Source of Variation* | *SS* | *df* | *MS* | *F* | *P-value* | *F crit* |
|---|---|---|---|---|---|---|
| Between Groups | 15664.5 | 1 | 15664.5 | 0.00711152 | 0.93553752 | 5.98737761 |
| Within Groups | 13216153.5 | 6 | 2202692.25 | | | |
| Total | 13231818 | 7 | | | | |

| RVΔL-Flpo 4 weeks | RVΔGL-Flpo 1 week |
|---|---|
| 4614 | 22 |
| 3006 | 18 |
| 1847 | 121 |
| 3331 | 2 |

Anova: Single Factor

SUMMARY

| *Groups* | *Count* | *Sum* | *Average* | *Variance* |
|---|---|---|---|---|
| RVΔL-Flpo 4 weeks | 4 | 12798 | 3199.5 | 1294933.67 |
| RVΔGL-Flpo 1 week | 4 | 163 | 40.75 | 2936.91667 |

ANOVA

| *Source of Variation* | *SS* | *df* | *MS* | *F* | *P-value* | *F crit* |
|---|---|---|---|---|---|---|
| Between Groups | 19955403.13 | 1 | 19955403.1 | 30.7509907 | 0.00145233 | 5.98737761 |
| Within Groups | 3893611.75 | 6 | 648935.292 | | | |
| Total | 23849014.88 | 7 | | | | |

| RVΔL-Flpo 1 week | RVΔL-Flpo 4 weeks |
|---|---|
| 3093 | 4614 |
| 320 | 3006 |
| 311 | 1847 |
| 263 | 3331 |
| 47 | 1369 |
| 2161 | 2919 |
| 365 | 3730 |
| 238 | 3454 |

Anova: Single Factor

SUMMARY

| Groups | Count | Sum | Average | Variance |
|---|---|---|---|---|
| RVΔL-Flpo 1 week | 8 | 6798 | 849.75 | 1274333.93 |
| RVΔL-Flpo 4 weeks | 8 | 24270 | 3033.75 | 1062946.79 |

ANOVA

| Source of Variation | SS | df | MS | F | P-value | F crit |
|---|---|---|---|---|---|---|
| Between Groups | 19079424 | 1 | 19079424 | 16.3261724 | 0.00121549 | 4.60010994 |
| Within Groups | 16360965 | 14 | 1168640.36 | | | |
| Total | 35440389 | 15 | | | | |

| RVΔGL-Flpo 1 week | RVΔL-Flpo 1 week |
|---|---|
| 22 | 3093 |
| 18 | 320 |
| 121 | 311 |
| 2 | 263 |
| 7 | 47 |
| 8 | 2161 |
| 98 | 365 |
| 6 | 238 |

Anova: Single Factor

SUMMARY

| Groups | Count | Sum | Average | Variance |
|---|---|---|---|---|
| RVΔGL-Flpo 1 week | 8 | 282 | 35.25 | 2180.78571 |
| RVΔL-Flpo 1 week | 8 | 6798 | 849.75 | 1274333.93 |

ANOVA

| Source of Variation | SS | df | MS | F | P-value | F crit |
|---|---|---|---|---|---|---|
| Between Groups | 2653641 | 1 | 2653641 | 4.1576348 | 0.06079134 | 4.60010994 |
| Within Groups | 8935603 | 14 | 638257.357 | | | |
| Total | 11589244 | 15 | | | | |

| RVΔGL-Cre 4 weeks | RVΔL-Cre 4 weeks |
|---|---|
| 19156 | 21743 |
| 17935 | 24773 |
| 18539 | 23026 |
| 17938 | 22272 |

Anova: Single Factor

SUMMARY

| Groups | Count | Sum | Average | Variance |
|---|---|---|---|---|
| RVΔGL-Cre 4 weeks | 4 | 73568 | 18392 | 340090 |
| RVΔL-Cre 4 weeks | 4 | 91814 | 22953.5 | 1748529.67 |

ANOVA

| Source of Variation | SS | df | MS | F | P-value | F crit |
|---|---|---|---|---|---|---|
| Between Groups | 41614564.5 | 1 | 41614564.5 | 39.8488678 | 0.00073769 | 5.98737761 |
| Within Groups | 6265859 | 6 | 1044309.83 | | | |
| Total | 47880423.5 | 7 | | | | |

| RVΔGL-Cre 4 weeks | RVΔGL-Cre 1 week |
|---|---|
| 19156 | 13484 |
| 17935 | 14635 |
| 18539 | 10647 |
| 17938 | 10845 |

Anova: Single Factor

SUMMARY

| *Groups* | *Count* | *Sum* | *Average* | *Variance* |
|---|---|---|---|---|
| RVΔGL-Cre 4 weeks | 4 | 73568 | 18392 | 340090 |
| RVΔGL-Cre 1 week | 4 | 49611 | 12402.75 | 3887094.92 |

ANOVA

| *Source of Variation* | *SS* | *df* | *MS* | *F* | *P-value* | *F crit* |
|---|---|---|---|---|---|---|
| Between Groups | 71742231.13 | 1 | 71742231.1 | 33.943266 | 0.00112486 | 5.98737761 |
| Within Groups | 12681554.75 | 6 | 2113592.46 | | | |
| Total | 84423785.88 | 7 | | | | |

| RVΔGL-Cre 4 weeks | RVΔL-Cre 1 week |
|---|---|
| 19156 | 16907 |
| 17935 | 17923 |
| 18539 | 18751 |
| 17938 | 16257 |

Anova: Single Factor

SUMMARY

| *Groups* | *Count* | *Sum* | *Average* | *Variance* |
|---|---|---|---|---|
| RVΔGL-Cre 4 weeks | 4 | 73568 | 18392 | 340090 |
| RVΔL-Cre 1 week | 4 | 69838 | 17459.5 | 1211355.67 |

ANOVA

| *Source of Variation* | *SS* | *df* | *MS* | *F* | *P-value* | *F crit* |
|---|---|---|---|---|---|---|
| Between Groups | 1739112.5 | 1 | 1739112.5 | 2.24192511 | 0.1849584 | 5.98737761 |
| Within Groups | 4654337 | 6 | 775722.833 | | | |
| Total | 6393449.5 | 7 | | | | |

| RVΔL-Cre 4 weeks | RVΔGL-Cre 1 week |
|---|---|
| 21743 | 13484 |
| 24773 | 14635 |
| 23026 | 10647 |
| 22272 | 10845 |

Anova: Single Factor

SUMMARY

| *Groups* | *Count* | *Sum* | *Average* | *Variance* |
|---|---|---|---|---|
| RVΔL-Cre 4 weeks | 4 | 91814 | 22953.5 | 1748529.67 |
| RVΔGL-Cre 1 week | 4 | 49611 | 12402.75 | 3887094.92 |

ANOVA

| *Source of Variation* | *SS* | *df* | *MS* | *F* | *P-value* | *F crit* |
|---|---|---|---|---|---|---|
| Between Groups | 222636651.1 | 1 | 222636651 | 79.010462 | 0.00011291 | 5.98737761 |
| Within Groups | 16906873.75 | 6 | 2817812.29 | | | |
| Total | 239543524.9 | 7 | | | | |

| RVΔL-Cre 4 weeks | RVΔL-Cre 1 week |
|---|---|
| 21743 | 16907 |
| 24773 | 17923 |
| 23026 | 18751 |
| 22272 | 16257 |

Anova: Single Factor

SUMMARY

| *Groups* | *Count* | *Sum* | *Average* | *Variance* |
|---|---|---|---|---|
| RVΔL-Cre 4 weeks | 4 | 91814 | 22953.5 | 1748529.67 |
| RVΔL-Cre 1 week | 4 | 69838 | 17459.5 | 1211355.67 |

ANOVA

| *Source of Variation* | *SS* | *df* | *MS* | *F* | *P-value* | *F crit* |
|---|---|---|---|---|---|---|
| Between Groups | 60368072 | 1 | 60368072 | 40.790818 | 0.00069327 | 5.98737761 |
| Within Groups | 8879656 | 6 | 1479942.67 | | | |
| Total | 69247728 | 7 | | | | |

| RVΔGL-Cre 1 week | RVΔL-Cre 1 week |
|---|---|
| 13484 | 16907 |
| 14635 | 17923 |
| 10647 | 18751 |
| 10845 | 16257 |

Anova: Single Factor

SUMMARY

| Groups | Count | Sum | Average | Variance |
|---|---|---|---|---|
| RVΔGL-Cre 1 week | 4 | 49611 | 12402.75 | 3887094.92 |
| RVΔL-Cre 1 week | 4 | 69838 | 17459.5 | 1211355.67 |

ANOVA

| Source of Variation | SS | df | MS | F | P-value | F crit |
|---|---|---|---|---|---|---|
| Between Groups | 51141441.13 | 1 | 51141441.1 | 20.061562 | 0.00419681 | 5.98737761 |
| Within Groups | 15295351.75 | 6 | 2549225.29 | | | |
| Total | 66436792.88 | 7 | | | | |

| RVΔL-5tTA 1 week | RVΔL-5tTA 4 weeks |
|---|---|
| 339 | 829 |
| 1411 | 1574 |
| 796 | 499 |
| 2271 | 1329 |

Anova: Single Factor

SUMMARY

| Groups | Count | Sum | Average | Variance |
|---|---|---|---|---|
| RVΔL-5tTA 1 week | 4 | 4817 | 1204.25 | 698675.583 |
| RVΔL-5tTA 4 weeks | 4 | 4231 | 1057.75 | 234872.917 |

ANOVA

| Source of Variation | SS | df | MS | F | P-value | F crit |
|---|---|---|---|---|---|---|
| Between Groups | 42924.5 | 1 | 42924.5 | 0.09195987 | 0.77193993 | 5.98737761 |
| Within Groups | 2800645.5 | 6 | 466774.25 | | | |
| Total | 2843570 | 7 | | | | |

| Comparison | p-value |
| --- | --- |
| RVΔGL-Flpo 4 weeks vs RVΔL-Flpo 4 weeks | **0.000320625** |
| RVΔGL-Flpo 4 weeks vs RVΔGL-Flpo 1 week | 0.27515341 |
| RVΔGL-Flpo 4 weeks vs RVΔL-Flpo 1 week | 0.93553752 |
| RVΔL-Flpo 4 weeks vs RVΔGL-Flpo 1 week | **0.001452333** |
| RVΔL-Flpo 4 weeks vs RVΔL-Flpo 1 week | **0.001215493** |
| RVΔGL-Flpo 1 week vs RVΔL-Flpo 1 week | 0.060791343 |
| RVΔGL-Cre 4 weeks vs RVΔL-Cre 4 weeks | **0.000737695** |
| RVΔGL-Cre 4 weeks vs RVΔGL-Cre 1 week | **0.001124862** |
| RVΔGL-Cre 4 weeks vs RVΔL-Cre 1 weeks | 0.184958403 |
| RVΔL-Cre 4 weeks vs RVΔGL-Cre 1 week | **0.000112909** |
| RVΔL-Cre 4 weeks vs RVΔL-Cre 1 week | **0.000693266** |
| RVΔGL-Cre 1 week vs RVΔL-Cre 1 week | **0.004196815** |
| RVΔL-5tTA 1 week vs RVΔL-5tTA 4 weeks | 0.77193993 |

**Supplementary File S4. Counts and statistics for spine density analysis, Related to Figure 2.** Spine numbers, lengths, and densities, followed by statistical comparisons.

*See following page.*

| Survival time | Mouse number | Section number | FOV number | Spine number | Length of dendrite (µm) | Spine density (number/µm) |
|---|---|---|---|---|---|---|
| 4 weeks | 21021901LJ | sec-8 | 1 | 48 | 63.93 | 0.75 |
| 4 weeks | 21021901LJ | sec-8 | 2 | 38 | 54.78 | 0.69 |
| 4 weeks | 21021901LJ | sec-8 | 3 | 39 | 44.15 | 0.88 |
| 4 weeks | 21021901LJ | sec-9 | 1 | 60 | 72.68 | 0.83 |
| 4 weeks | 21021901LJ | sec-9 | 2 | 32 | 45.99 | 0.70 |
| 4 weeks | 21021901LJ | sec-9 | 3 | 46 | 56.99 | 0.81 |
| 4 weeks | 21021901LJ | sec-9 | 4 | 42 | 44.60 | 0.94 |
| 4 weeks | 21021901LJ | sec-11 | 1 | 42 | 48.80 | 0.86 |
| 4 weeks | 21021901LJ | sec-11 | 2 | 27 | 41.14 | 0.66 |
| 4 weeks | 21021902LJ | sec-10 | 1 | 39 | 53.81 | 0.72 |
| 4 weeks | 21021902LJ | sec-10 | 2 | 34 | 47.88 | 0.71 |
| 4 weeks | 21021902LJ | sec-10 | 3 | 43 | 49.02 | 0.88 |
| 4 weeks | 21021902LJ | sec-10 | 4 | 45 | 58.15 | 0.77 |
| 4 weeks | 21021902LJ | sec-10 | 5 | 48 | 54.85 | 0.88 |
| 4 weeks | Mean: | | | | | 0.79 |
| 6 months | 18080701NL | sec-08 | 1 | 35 | 54.28 | 0.64 |
| 6 months | 18080701NL | sec-08 | 2 | 33 | 48.03 | 0.69 |
| 6 months | 18080701NL | sec-11 | 1 | 29 | 33.23 | 0.87 |
| 6 months | 18080701NL | sec-11 | 2 | 24 | 27.65 | 0.87 |
| 6 months | 18080701NL | sec-11 | 3 | 10 | 13.87 | 0.72 |
| 6 months | 18080701NL | sec-11 | 4 | 29 | 37.45 | 0.77 |
| 6 months | 18080702NL | sec-10 | 1 | 23 | 27.09 | 0.85 |
| 6 months | 18080702NL | sec-10 | 2 | 39 | 45.03 | 0.87 |
| 6 months | 18080702NL | sec-10 | 3 | 33 | 47.20 | 0.70 |
| 6 months | 18080702NL | sec-11 | 1 | 22 | 37.11 | 0.59 |
| 6 months | 18080702NL | sec-11 | 2 | 9 | 13.48 | 0.67 |
| 6 months | 18080702NL | sec-11 | 3 | 27 | 33.20 | 0.81 |
| 6 months | 18080702NL | sec-11 | 4 | 12 | 18.81 | 0.64 |
| 6 months | 18080702NL | sec-11 | 5 | 20 | 28.03 | 0.71 |
| 6 months | 18080702NL | sec-11 | 6 | 25 | 28.91 | 0.86 |
| 6 months | Mean: | | | | | 0.75 |

Anova: Single Factor

SUMMARY

| *Groups* | *Count* | *Sum* | *Average* | *Variance* |
|---|---|---|---|---|
| 4 weeks | 14 | 11.07599868 | 0.791142763 | 0.007811633 |
| 6 months | 15 | 11.27228792 | 0.751485861 | 0.009561113 |

ANOVA

| *Source of Variation* | *SS* | *df* | *MS* | *F* | *P-value* | *F crit* |
|---|---|---|---|---|---|---|
| Between Groups | 0.011388299 | 1 | 0.011388299 | 1.306181705 | **0.263120437** | 4.210008468 |
| Within Groups | 0.235406815 | 27 | 0.008718771 | | | |
| Total | 0.246795114 | 28 | | | | |

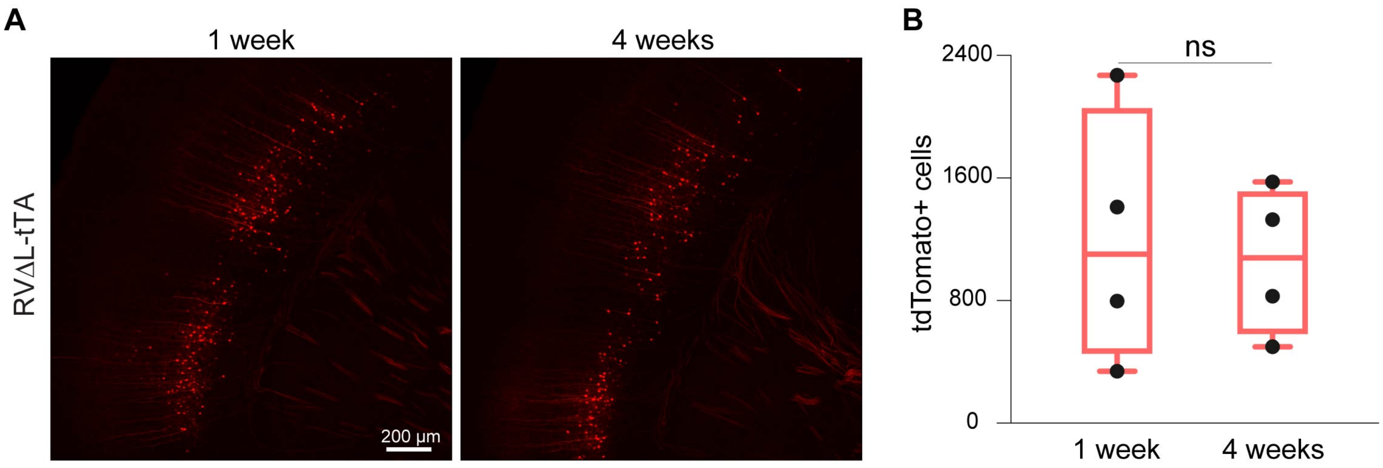

**Supplementary Figure S4. Retrograde targeting with third-generation (ΔL) rabies virus expressing the tetracycline transactivator, Related to Figure 2.**
(A) Corticothalamic neurons retrogradely labeled by a ΔL virus expressing tTA injected in the somatosensory thalamus of Ai63 reporter mice (tdTomato driven by TRE-tight) 1 week (left image) or 4 weeks (right image) prior to perfusion. Scale bar: 200 µm, applies to both images.
(B) Counts of labeled cortical neurons, with each data point being the total number found in one series consisting of every sixth 50 µm section from a given brain - see Methods). Numbers are not significantly different between the two time points (single factor ANOVA, $p = 0.772$, $n = 4$ mice per group). See Supplementary File S3 for counts and statistics).

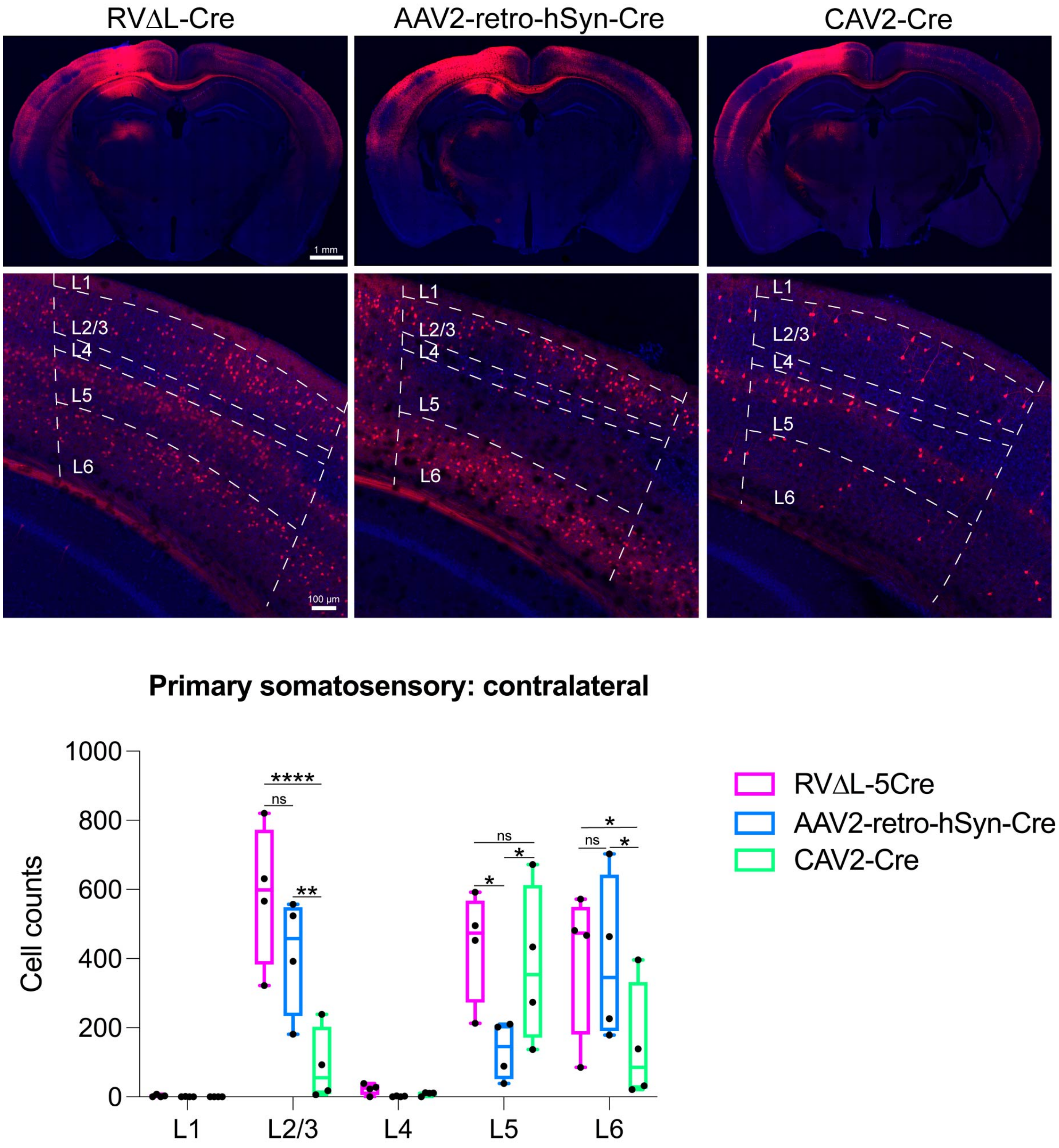

**Supplementary Figure S5. Retrograde labeling by ΔL rabies virus compared to rAAV2-retro and CAV-2: injections in anteromedial visual cortex (AM), Related to Figure 3.**
RVΔL-Cre, rAAV2-retro-hSyn-Cre (from Addgene), or CAV-Cre (from the Plateforme de Vectorologie de Montpellier) was injected undiluted into the cortical anteromedial area (AM) of reporter mice, with injections being of equal volumes (200 µl); after a 4-week survival time, brain sections were imaged and labeled neurons in several brain regions were counted. Note that each data point is the total number in one series consisting of every sixth 50 µm section from a given brain (see Methods) so that the total number of labeled S1 neurons in each brain would be approximately six times the corresponding number shown here. In contralateral primary somatosensory cortex, RVΔL labeled more cells in layers 2/3, 5, and 6 than did either of the other two viruses, although the difference between RVΔL and the “runner up” in each case (rAAV2-retro in layers 2/3 and 6, CAV-2 in layer 5) was not statistically significant. Few neurons were labeled in

layers 1 and 4, and the differences in these layers were not significant. See Supplementary File S5 for all counts and statistical comparisons. See also Supplementary Figures S6 – S11 for sets of high-resolution confocal images of series of coronal sections from mice labeled with each of the three viruses for each of the two injection sites.

**Supplementary Figures S6a-t. High-resolution whole-section confocal images of retrograde labeling by viruses injected in anterior cingulate area (ACA): rAAV2-retro-hSyn-Cre, Related to Figure 3.**

*See external file.*

**Supplementary Figures S7a-t. High-resolution whole-section confocal images of retrograde labeling by viruses injected in anterior cingulate area (ACA): CAV2-Cre, Related to Figure 3.**

*See external file.*

**Supplementary Figures S8a-t. High-resolution whole-section confocal images of retrograde labeling by viruses injected in anterior cingulate area (ACA): RV∆L-Cre, Related to Figure 3.**

*See external file.*

**Supplementary Figures S9a-t. High-resolution whole-section confocal images of retrograde labeling by viruses injected in anteromedial visual cortex (AM): rAAV2-retro-hSyn-Cre, Related to Figure 3.**

*See external file.*

**Supplementary Figures S10a-t. High-resolution whole-section confocal images of retrograde labeling by viruses injected in anteromedial visual cortex (AM): CAV2-Cre, Related to Figure 3.**

*See external file.*

**Supplementary Figures S11a-t. High-resolution whole-section confocal images of retrograde labeling by viruses injected in anteromedial visual cortex (AM): RV∆L-Cre, Related to Figure 3.**

*See external file.*

**Supplementary File S5. Cell counts and statistics for RV$\Delta$L vs rAAV2-retro vs CAV-Cre retrograde targeting experiments, Related to Figure 3.**

Cell counts in individual sections, followed by totals for each mouse, followed by statistical comparisons, for both ACA and AM injections.

*See following pages.*

**ACA injections**

Basolateral amygdala (BLA)

| Virus group | Animal ID | Section Nr: | Cell counts |
|---|---|---|---|
| CAV2-Cre | 22073101LJ | sec-14 | 3 |
| | 22073101LJ | sec-15 | 8 |
| | 22073101LJ | sec-16 | 18 |
| | 22073101LJ | sec-17 | 4 |
| | 22073101LJ | Sum: | 33 |
| | 22073102LJ | sec-16 | 7 |
| | 22073102LJ | sec-17 | 3 |
| | 22073102LJ | Sum: | 10 |
| | 22073103LJ | sec-14 | 1 |
| | 22073103LJ | sec-15 | 2 |
| | 22073103LJ | sec-16 | 14 |
| | 22073103LJ | sec-17 | 7 |
| | 22073103LJ | Sum: | 24 |
| | 22073104LJ | sec-14 | 2 |
| | 22073104LJ | sec-15 | 10 |
| | 22073104LJ | sec-16 | 4 |
| | 22073104LJ | Sum: | 16 |
| AAV2-retro-hSyn-Cre | 22072201LJ | sec-14 | 36 |
| | 22072201LJ | sec-15 | 64 |
| | 22072201LJ | sec-16 | 25 |
| | 22072201LJ | Sum: | 125 |
| | 22072202LJ | sec-14 | 34 |
| | 22072202LJ | sec-15 | 80 |
| | 22072202LJ | sec-16 | 64 |
| | 22072202LJ | sec-17 | 9 |
| | 22072202LJ | Sum: | 187 |
| | 22072203LJ | sec-14 | 15 |
| | 22072203LJ | sec-15 | 51 |
| | 22072203LJ | sec-16 | 52 |
| | 22072203LJ | sec-17 | 3 |
| | 22072203LJ | Sum: | 121 |
| | 22072204LJ | sec-13 | 8 |
| | 22072204LJ | sec-14 | 83 |
| | 22072204LJ | sec-15 | 73 |
| | 22072204LJ | sec-16 | 19 |
| | 22072204LJ | Sum: | 183 |
| RVΔL-5Cre | 22072101LJ | sec-15 | 23 |
| | 22072101LJ | sec-16 | 30 |
| | 22072101LJ | sec-17 | 5 |
| | 22072101LJ | Sum: | 58 |
| | 22072102LJ | sec-12 | 8 |
| | 22072102LJ | sec-13 | 26 |
| | 22072102LJ | sec-14 | 28 |
| | 22072102LJ | sec-15 | 1 |
| | 22072102LJ | Sum: | 63 |
| | 22072103LJ | sec-13 | 6 |
| | 22072103LJ | sec-14 | 34 |
| | 22072103LJ | sec-15 | 18 |
| | 22072103LJ | sec-16 | 4 |
| | 22072103LJ | Sum: | 62 |
| | 22072104LJ | sec-14 | 4 |
| | 22072104LJ | sec-15 | 48 |
| | 22072104LJ | sec-16 | 33 |
| | 22072104LJ | sec-17 | 12 |
| | 22072104LJ | Sum: | 97 |

Hippocampus

| Virus group | Animal ID | Section Nr: | Cell counts |
|---|---|---|---|
| CAV2-Cre | 22073101LJ | sec-19 | 1 |
| | 22073101LJ | Sum: | 1 |
| | 22073102LJ | sec-18 | 1 |
| | 22073102LJ | sec-20 | 4 |
| | 22073102LJ | Sum: | 5 |

| | | | |
|---|---|---|---|
| | 22073103LJ | sec-17 | 1 |
| | 22073103LJ | sec-20 | 5 |
| | 22073103LJ | Sum: | 6 |
| | 22073104LJ | sec-17 | 4 |
| | 22073104LJ | sec-19 | 9 |
| | 22073104LJ | sec-20 | 6 |
| | 22073104LJ | Sum: | 19 |
| AAV2-retro-hSyn-Cre | 22072201LJ | sec-15 | 51 |
| | 22072201LJ | sec-16 | 19 |
| | 22072201LJ | sec-17 | 5 |
| | 22072201LJ | sec-18 | 10 |
| | 22072201LJ | sec-19 | 12 |
| | 22072201LJ | sec-20 | 40 |
| | 22072201LJ | Sum: | 137 |
| | 22072202LJ | sec-14 | 7 |
| | 22072202LJ | sec-16 | 39 |
| | 22072202LJ | sec-17 | 40 |
| | 22072202LJ | sec-18 | 35 |
| | 22072202LJ | sec-19 | 69 |
| | 22072202LJ | sec-20 | 92 |
| | 22072202LJ | Sum: | 282 |
| | 22072203LJ | sec-14 | 1 |
| | 22072203LJ | sec-15 | 56 |
| | 22072203LJ | sec-16 | 13 |
| | 22072203LJ | sec-17 | 10 |
| | 22072203LJ | sec-18 | 12 |
| | 22072203LJ | sec-19 | 29 |
| | 22072203LJ | sec-20 | 49 |
| | 22072203LJ | Sum: | 170 |
| | 22072204LJ | sec-14 | 6 |
| | 22072204LJ | sec-15 | 7 |
| | 22072204LJ | sec-16 | 6 |
| | 22072204LJ | sec-17 | 5 |
| | 22072204LJ | sec-19 | 11 |
| | 22072204LJ | sec-20 | 2 |
| | 22072204LJ | Sum: | 37 |
| RVΔL-5Cre | 22072101LJ | sec-11 | 1 |
| | 22072101LJ | sec-12 | 30 |
| | 22072101LJ | sec-13 | 52 |
| | 22072101LJ | sec-14 | 34 |
| | 22072101LJ | sec-15 | 39 |
| | 22072101LJ | sec-16 | 54 |
| | 22072101LJ | sec-17 | 111 |
| | 22072101LJ | sec-18 | 92 |
| | 22072101LJ | sec-19 | 88 |
| | 22072101LJ | sec-20 | 82 |
| | 22072101LJ | Sum: | 583 |
| | 22072102LJ | sec-11 | 48 |
| | 22072102LJ | sec-12 | 117 |
| | 22072102LJ | sec-13 | 107 |
| | 22072102LJ | sec-14 | 111 |
| | 22072102LJ | sec-15 | 140 |
| | 22072102LJ | sec-16 | 177 |
| | 22072102LJ | sec-17 | 269 |
| | 22072102LJ | sec-18 | 209 |
| | 22072102LJ | sec-19 | 178 |
| | 22072102LJ | Sum: | 1356 |
| | 22072103LJ | sec-13 | 20 |
| | 22072103LJ | sec-14 | 30 |
| | 22072103LJ | sec-15 | 33 |
| | 22072103LJ | sec-16 | 32 |
| | 22072103LJ | sec-17 | 20 |
| | 22072103LJ | sec-18 | 64 |
| | 22072103LJ | sec-19 | 127 |
| | 22072103LJ | sec-20 | 59 |
| | 22072103LJ | Sum: | 385 |
| | 22072104LJ | sec-14 | 22 |
| | 22072104LJ | sec-15 | 32 |

| | | | |
|---|---|---|---|
| | 22072104LJ | sec-16 | 29 |
| | 22072104LJ | sec-17 | 33 |
| | 22072104LJ | sec-18 | 27 |
| | 22072104LJ | sec-19 | 92 |
| | 22072104LJ | sec-20 | 161 |
| | 22072104LJ | Sum: | 396 |

Primary motor area

| Virus group | Animal ID | Section Nr: | L1 | L2/3 | L4 | L5 | L6 |
|---|---|---|---|---|---|---|---|
| CAV2-Cre | 22073101LJ | sec-1 | 0 | 2 | 3 | 5 | 1 |
| | 22073101LJ | sec-3 | 0 | 7 | 0 | 4 | 10 |
| | 22073101LJ | sec-5 | 0 | 9 | 0 | 1 | 3 |
| | 22073101LJ | sec-6 | 0 | 3 | 0 | 22 | 4 |
| | 22073101LJ | sec-7 | 3 | 2 | 0 | 53 | 26 |
| | 22073101LJ | sec-8 | 5 | 2 | 0 | 56 | 30 |
| | 22073101LJ | sec-9 | 1 | 2 | 0 | 42 | 40 |
| | 22073101LJ | sec-10 | 1 | 2 | 0 | 14 | 32 |
| | 22073101LJ | sec-11 | 1 | 2 | 0 | 7 | 26 |
| | 22073101LJ | sec-12 | 1 | 3 | 0 | 12 | 12 |
| | 22073101LJ | sec-13 | 0 | 7 | 0 | 27 | 20 |
| | 22073101LJ | sec-14 | 0 | 8 | 0 | 39 | 40 |
| | 22073101LJ | sec-15 | 0 | 11 | 0 | 55 | 45 |
| | 22073101LJ | Sum: | 12 | 58 | 0 | 332 | 288 |
| | 22073102LJ | sec-2 | 0 | 0 | 0 | 1 | 0 |
| | 22073102LJ | sec-4 | 1 | 0 | 0 | 1 | 0 |
| | 22073102LJ | sec-5 | 0 | 3 | 0 | 0 | 5 |
| | 22073102LJ | sec-6 | 0 | 6 | 0 | 1 | 4 |
| | 22073102LJ | sec-7 | 0 | 11 | 0 | 96 | 20 |
| | 22073102LJ | sec-8 | 4 | 18 | 0 | 92 | 129 |
| | 22073102LJ | sec-9 | 2 | 1 | 0 | 98 | 98 |
| | 22073102LJ | sec-10 | 0 | 5 | 0 | 2 | 27 |
| | 22073102LJ | sec-11 | 2 | 3 | 0 | 10 | 8 |
| | 22073102LJ | sec-12 | 0 | 6 | 0 | 5 | 17 |
| | 22073102LJ | sec-13 | 0 | 10 | 0 | 42 | 40 |
| | 22073102LJ | sec-14 | 0 | 7 | 0 | 80 | 39 |
| | 22073102LJ | sec-15 | 0 | 11 | 0 | 84 | 48 |
| | 22073102LJ | Sum: | 9 | 81 | 0 | 512 | 435 |
| | 22073103LJ | sec-1 | 0 | 0 | 0 | 14 | 0 |
| | 22073103LJ | sec-2 | 0 | 0 | 0 | 7 | 0 |
| | 22073103LJ | sec-3 | 0 | 0 | 0 | 2 | 0 |
| | 22073103LJ | sec-4 | 0 | 2 | 0 | 5 | 5 |
| | 22073103LJ | sec-5 | 2 | 8 | 0 | 16 | 5 |
| | 22073103LJ | sec-6 | 4 | 4 | 0 | 54 | 25 |
| | 22073103LJ | sec-7 | 3 | 9 | 0 | 66 | 42 |
| | 22073103LJ | sec-8 | 4 | 4 | 0 | 66 | 45 |
| | 22073103LJ | sec-9 | 0 | 3 | 0 | 28 | 46 |
| | 22073103LJ | sec-10 | 3 | 4 | 0 | 33 | 43 |
| | 22073103LJ | sec-11 | 2 | 3 | 0 | 37 | 49 |
| | 22073103LJ | sec-12 | 0 | 5 | 0 | 25 | 26 |
| | 22073103LJ | sec-13 | 0 | 4 | 0 | 49 | 40 |
| | 22073103LJ | sec-14 | 0 | 9 | 0 | 67 | 43 |
| | 22073103LJ | sec-15 | 0 | 9 | 0 | 66 | 34 |
| | 22073103LJ | Sum: | 19 | 64 | 0 | 535 | 403 |
| | 22073104LJ | sec-4 | 0 | 0 | 0 | 0 | 3 |
| | 22073104LJ | sec-5 | 0 | 1 | 0 | 0 | 1 |
| | 22073104LJ | sec-6 | 0 | 3 | 0 | 14 | 6 |
| | 22073104LJ | sec-7 | 2 | 4 | 0 | 19 | 18 |
| | 22073104LJ | sec-8 | 2 | 1 | 0 | 29 | 7 |
| | 22073104LJ | sec-9 | 3 | 0 | 0 | 6 | 8 |
| | 22073104LJ | sec-10 | 2 | 2 | 0 | 12 | 13 |
| | 22073104LJ | sec-11 | 0 | 7 | 0 | 33 | 33 |
| | 22073104LJ | sec-12 | 0 | 5 | 0 | 51 | 30 |
| | 22073104LJ | sec-13 | 0 | 5 | 0 | 62 | 33 |
| | 22073104LJ | sec-14 | 0 | 14 | 0 | 68 | 37 |
| | 22073104LJ | sec-15 | 0 | 9 | 0 | 60 | 34 |
| | 22073104LJ | Sum: | 9 | 51 | 0 | 354 | 223 |
| AAV2-retro-hSyn-Cre | 22072201LJ | sec-3 | 0 | 0 | 0 | 6 | 0 |

| | | | | | | | |
|---|---|---|---|---|---|---|---|
| | 22072201LJ | sec-4 | 0 | 1 | 0 | 3 | 0 |
| | 22072201LJ | sec-5 | 0 | 2 | 0 | 6 | 0 |
| | 22072201LJ | sec-6 | 0 | 24 | 0 | 14 | 5 |
| | 22072201LJ | sec-7 | 0 | 38 | 0 | 12 | 15 |
| | 22072201LJ | sec-8 | 0 | 43 | 0 | 9 | 18 |
| | 22072201LJ | sec-9 | 0 | 42 | 0 | 2 | 4 |
| | 22072201LJ | sec-10 | 0 | 29 | 0 | 4 | 28 |
| | 22072201LJ | sec-11 | 0 | 45 | 0 | 12 | 22 |
| | 22072201LJ | sec-12 | 0 | 50 | 0 | 80 | 65 |
| | 22072201LJ | sec-13 | 0 | 55 | 0 | 58 | 59 |
| | 22072201LJ | sec-14 | 0 | 66 | 0 | 78 | 35 |
| | 22072201LJ | Sum: | 0 | 395 | 0 | 284 | 251 |
| | 22072202LJ | sec-4 | 0 | 1 | 0 | 0 | 9 |
| | 22072202LJ | sec-5 | 0 | 16 | 0 | 3 | 6 |
| | 22072202LJ | sec-6 | 1 | 23 | 0 | 16 | 19 |
| | 22072202LJ | sec-7 | 0 | 49 | 0 | 21 | 24 |
| | 22072202LJ | sec-8 | 0 | 64 | 0 | 28 | 57 |
| | 22072202LJ | sec-9 | 0 | 45 | 0 | 23 | 33 |
| | 22072202LJ | sec-10 | 0 | 30 | 0 | 12 | 32 |
| | 22072202LJ | sec-11 | 0 | 34 | 0 | 14 | 35 |
| | 22072202LJ | sec-12 | 0 | 36 | 0 | 18 | 29 |
| | 22072202LJ | sec-13 | 0 | 47 | 0 | 28 | 41 |
| | 22072202LJ | sec-14 | 0 | 76 | 0 | 72 | 70 |
| | 22072202LJ | sec-15 | 0 | 40 | 0 | 56 | 42 |
| | 22072202LJ | Sum: | 1 | 461 | 0 | 291 | 397 |
| | 22072203LJ | sec-1 | 0 | 4 | 0 | 2 | 0 |
| | 22072203LJ | sec-5 | 0 | 1 | 0 | 3 | 3 |
| | 22072203LJ | sec-6 | 0 | 9 | 0 | 3 | 3 |
| | 22072203LJ | sec-7 | 1 | 32 | 0 | 23 | 32 |
| | 22072203LJ | sec-8 | 0 | 32 | 0 | 10 | 57 |
| | 22072203LJ | sec-9 | 0 | 43 | 0 | 17 | 26 |
| | 22072203LJ | sec-10 | 0 | 51 | 0 | 16 | 64 |
| | 22072203LJ | sec-11 | 0 | 62 | 0 | 26 | 35 |
| | 22072203LJ | sec-12 | 0 | 74 | 0 | 48 | 64 |
| | 22072203LJ | sec-13 | 0 | 101 | 0 | 90 | 72 |
| | 22072203LJ | Sum: | 1 | 409 | 0 | 238 | 356 |
| | 22072204LJ | sec-2 | 0 | 3 | 0 | 0 | 0 |
| | 22072204LJ | sec-3 | 0 | 51 | 0 | 22 | 0 |
| | 22072204LJ | sec-4 | 0 | 70 | 0 | 22 | 50 |
| | 22072204LJ | sec-5 | 0 | 110 | 0 | 8 | 30 |
| | 22072204LJ | sec-6 | 0 | 100 | 0 | 44 | 35 |
| | 22072204LJ | sec-7 | 0 | 50 | 0 | 35 | 52 |
| | 22072204LJ | sec-8 | 0 | 40 | 0 | 15 | 28 |
| | 22072204LJ | sec-9 | 9 | 15 | 0 | 10 | 28 |
| | 22072204LJ | sec-10 | 13 | 40 | 0 | 20 | 54 |
| | 22072204LJ | sec-11 | 13 | 34 | 0 | 12 | 40 |
| | 22072204LJ | sec-12 | 3 | 30 | 0 | 25 | 35 |
| | 22072204LJ | sec-13 | 0 | 20 | 0 | 28 | 43 |
| | 22072204LJ | Sum: | 38 | 563 | 0 | 241 | 395 |
| RVΔL-5Cre | 22072101LJ | sec-1 | 0 | 0 | 0 | 21 | 0 |
| | 22072101LJ | sec-3 | 0 | 1 | 0 | 38 | 0 |
| | 22072101LJ | sec-4 | 0 | 27 | 0 | 5 | 0 |
| | 22072101LJ | sec-5 | 0 | 20 | 0 | 13 | 22 |
| | 22072101LJ | sec-6 | 5 | 105 | 0 | 62 | 34 |
| | 22072101LJ | sec-7 | 2 | 154 | 0 | 242 | 70 |
| | 22072101LJ | sec-8 | 0 | 135 | 0 | 174 | 164 |
| | 22072101LJ | sec-9 | 2 | 45 | 0 | 149 | 166 |
| | 22072101LJ | sec-10 | 0 | 28 | 0 | 20 | 34 |
| | 22072101LJ | sec-11 | 0 | 27 | 0 | 52 | 53 |
| | 22072101LJ | sec-12 | 0 | 93 | 0 | 47 | 120 |
| | 22072101LJ | sec-13 | 0 | 169 | 0 | 90 | 102 |
| | 22072101LJ | Sum: | 9 | 804 | 0 | 913 | 765 |
| | 22072102LJ | sec-1 | 0 | 0 | 0 | 19 | 0 |
| | 22072102LJ | sec-2 | 0 | 0 | 0 | 9 | 0 |
| | 22072102LJ | sec-3 | 0 | 0 | 0 | 3 | 0 |
| | 22072102LJ | sec-4 | 0 | 14 | 0 | 2 | 0 |
| | 22072102LJ | sec-5 | 0 | 110 | 0 | 69 | 39 |
| | 22072102LJ | sec-6 | 5 | 142 | 0 | 140 | 99 |

| | | | | | | |
|---|---|---|---|---|---|---|
| 22072102LJ | sec-7 | 0 | 180 | 0 | 105 | 134 |
| 22072102LJ | sec-8 | 0 | 136 | 0 | 190 | 234 |
| 22072102LJ | sec-9 | 1 | 265 | 0 | 210 | 267 |
| 22072102LJ | sec-10 | 0 | 94 | 0 | 88 | 136 |
| 22072102LJ | sec-11 | 0 | 160 | 0 | 153 | 230 |
| 22072102LJ | sec-12 | 0 | 95 | 0 | 108 | 123 |
| 22072102LJ | sec-13 | 0 | 70 | 0 | 138 | 89 |
| 22072102LJ | Sum: | 6 | 1266 | 0 | 1234 | 1351 |
| 22072103LJ | sec-1 | 0 | 0 | 0 | 15 | 0 |
| 22072103LJ | sec-2 | 0 | 0 | 0 | 12 | 0 |
| 22072103LJ | sec-3 | 0 | 0 | 0 | 10 | 0 |
| 22072103LJ | sec-4 | 0 | 1 | 0 | 6 | 0 |
| 22072103LJ | sec-5 | 0 | 9 | 0 | 5 | 0 |
| 22072103LJ | sec-6 | 0 | 25 | 0 | 19 | 18 |
| 22072103LJ | sec-7 | 0 | 117 | 0 | 51 | 40 |
| 22072103LJ | sec-8 | 1 | 133 | 0 | 101 | 129 |
| 22072103LJ | sec-9 | 3 | 28 | 0 | 60 | 94 |
| 22072103LJ | sec-10 | 0 | 66 | 0 | 47 | 80 |
| 22072103LJ | sec-11 | 1 | 82 | 0 | 112 | 93 |
| 22072103LJ | sec-12 | 0 | 104 | 0 | 120 | 115 |
| 22072103LJ | sec-13 | 0 | 104 | 0 | 129 | 93 |
| 22072103LJ | sec-14 | 0 | 113 | 0 | 110 | 90 |
| 22072103LJ | Sum: | 5 | 782 | 0 | 797 | 752 |
| 22072104LJ | sec-1 | 0 | 0 | 0 | 20 | 0 |
| 22072104LJ | sec-2 | 0 | 5 | 0 | 14 | 0 |
| 22072104LJ | sec-3 | 2 | 6 | 0 | 3 | 6 |
| 22072104LJ | sec-4 | 0 | 12 | 0 | 5 | 8 |
| 22072104LJ | sec-5 | 2 | 30 | 0 | 1 | 8 |
| 22072104LJ | sec-6 | 0 | 42 | 0 | 46 | 14 |
| 22072104LJ | sec-7 | 0 | 57 | 0 | 27 | 37 |
| 22072104LJ | sec-8 | 0 | 79 | 0 | 40 | 22 |
| 22072104LJ | sec-9 | 0 | 99 | 0 | 64 | 82 |
| 22072104LJ | sec-10 | 0 | 59 | 0 | 90 | 138 |
| 22072104LJ | sec-11 | 0 | 71 | 0 | 60 | 110 |
| 22072104LJ | sec-12 | 0 | 53 | 0 | 51 | 105 |
| 22072104LJ | sec-13 | 0 | 69 | 0 | 71 | 86 |
| 22072104LJ | sec-14 | 0 | 76 | 0 | 113 | 76 |
| 22072104LJ | Sum: | 4 | 658 | 0 | 605 | 692 |

**Summary for ACA injections**

Basolateral amygdala (BLA)

| Virus group | Animal ID | Sum of cell counts |
|---|---|---|
| CAV2-Cre | 22073101LJ | 33 |
| CAV2-Cre | 22073102LJ | 10 |
| CAV2-Cre | 22073103LJ | 24 |
| CAV2-Cre | 22073104LJ | 16 |
| AAV2-retro-hSyn-Cre | 22072201LJ | 125 |
| AAV2-retro-hSyn-Cre | 22072202LJ | 187 |
| AAV2-retro-hSyn-Cre | 22072203LJ | 121 |
| AAV2-retro-hSyn-Cre | 22072204LJ | 183 |
| RVΔL-5Cre | 22072101LJ | 58 |
| RVΔL-5Cre | 22072102LJ | 63 |
| RVΔL-5Cre | 22072103LJ | 62 |
| RVΔL-5Cre | 22072104LJ | 97 |

Hippocampus

| Virus group | Animal ID | Sum of cell counts |
|---|---|---|
| CAV2-Cre | 22073101LJ | 1 |
| CAV2-Cre | 22073102LJ | 5 |
| CAV2-Cre | 22073103LJ | 6 |
| CAV2-Cre | 22073104LJ | 19 |
| AAV2-retro-hSyn-Cre | 22072201LJ | 137 |
| AAV2-retro-hSyn-Cre | 22072202LJ | 282 |
| AAV2-retro-hSyn-Cre | 22072203LJ | 170 |
| AAV2-retro-hSyn-Cre | 22072204LJ | 37 |
| RVΔL-5Cre | 22072101LJ | 583 |
| RVΔL-5Cre | 22072102LJ | 1356 |
| RVΔL-5Cre | 22072103LJ | 385 |
| RVΔL-5Cre | 22072104LJ | 396 |

Primary motor area

| Virus group | Animal ID | L1 | L2/3 | L4 | L5 | L6 | Sum of all layers |
|---|---|---|---|---|---|---|---|
| CAV2-Cre | 22073101LJ | 12 | 58 | 0 | 332 | 288 | 690 |
| CAV2-Cre | 22073102LJ | 9 | 81 | 0 | 512 | 435 | 1037 |
| CAV2-Cre | 22073103LJ | 19 | 64 | 0 | 535 | 403 | 1021 |
| CAV2-Cre | 22073104LJ | 9 | 51 | 0 | 354 | 223 | 637 |
| AAV2-retro-hSyn-Cre | 22072201LJ | 0 | 395 | 0 | 284 | 251 | 930 |
| AAV2-retro-hSyn-Cre | 22072202LJ | 1 | 461 | 0 | 291 | 397 | 1150 |
| AAV2-retro-hSyn-Cre | 22072203LJ | 1 | 409 | 0 | 238 | 356 | 1004 |
| AAV2-retro-hSyn-Cre | 22072204LJ | 38 | 563 | 0 | 241 | 395 | 1237 |
| RVΔL-5Cre | 22072101LJ | 9 | 804 | 0 | 913 | 765 | 2491 |
| RVΔL-5Cre | 22072102LJ | 6 | 1266 | 0 | 1234 | 1351 | 3857 |
| RVΔL-5Cre | 22072103LJ | 5 | 782 | 0 | 797 | 752 | 2336 |
| RVΔL-5Cre | 22072104LJ | 4 | 658 | 0 | 605 | 692 | 1959 |

**Ordinary one-way ANOVA of BLA**

| | |
|---|---|
| Number of families | 1 |
| Number of comparisons per family | 3 |
| Alpha | 0.05 |

| Tukey's multiple comparisons test | Mean Diff. | 95.00% CI of diff. | Below threshold? | Summary | Adjusted P Value | |
|---|---|---|---|---|---|---|
| CAV2-Cre vs. AAV2-retro-hSyn-Cre | -133.3 | -180.5 to -86.05 | Yes | **** | <0.0001 | A-B |
| CAV2-Cre vs. RV∆L-5Cre | -49.25 | -96.45 to -2.048 | Yes | * | 0.0414 | A-C |
| AAV2-retro-hSyn-Cre vs. RV∆L-5Cre | 84 | 36.80 to 131.2 | Yes | ** | 0.002 | B-C |

| Test details | Mean 1 | Mean 2 | Mean Diff. | SE of diff. | n1 | n2 | q | DF |
|---|---|---|---|---|---|---|---|---|
| CAV2-Cre vs. AAV2-retro-hSyn-Cre | 20.75 | 154 | -133.3 | 16.91 | 4 | 4 | 11.15 | 9 |
| CAV2-Cre vs. RV∆L-5Cre | 20.75 | 70 | -49.25 | 16.91 | 4 | 4 | 4.12 | 9 |
| AAV2-retro-hSyn-Cre vs. RV∆L-5Cre | 154 | 70 | 84 | 16.91 | 4 | 4 | 7.027 | 9 |

**Ordinary one-way ANOVA of Hippocampus**

| | |
|---|---|
| Number of families | 1 |
| Number of comparisons per family | 3 |
| Alpha | 0.05 |

| Tukey's multiple comparisons test | Mean Diff. | 95.00% CI of diff. | Below threshold? | Summary | Adjusted P Value | |
|---|---|---|---|---|---|---|
| CAV2-Cre vs. AAV2-retro-hSyn-Cre | -148.8 | -685.3 to 387.8 | No | ns | 0.7274 | A-B |
| CAV2-Cre vs. RVΔL-5Cre | -672.3 | -1209 to -135.7 | Yes | * | 0.0167 | A-C |
| AAV2-retro-hSyn-Cre vs. RVΔL-5Cre | -523.5 | -1060 to 13.09 | No | ns | 0.0556 | B-C |

| Test details | Mean 1 | Mean 2 | Mean Diff. | SE of diff. | n1 | n2 |
|---|---|---|---|---|---|---|
| CAV2-Cre vs. AAV2-retro-hSyn-Cre | 7.75 | 156.5 | -148.8 | 192.2 | 4 | 4 |
| CAV2-Cre vs. RVΔL-5Cre | 7.75 | 680 | -672.3 | 192.2 | 4 | 4 |
| AAV2-retro-hSyn-Cre vs. RVΔL-5Cre | 156.5 | 680 | -523.5 | 192.2 | 4 | 4 |

**2 way ANOVA of Primary motor area**
Within each row, compare columns (simple effects within rows)

| | |
|---|---|
| Number of families | 4 |
| Number of comparisons per family | 3 |
| Alpha | 0.05 |

| Tukey's multiple comparisons test | Mean Diff. | 95.00% CI of diff. | Below thres | Summary | Adjusted P Value |
|---|---|---|---|---|---|
| L1 | | | | | |
| CAV2-Cre vs. AAV2-retro-hSyn-Cre | 2.25 | -256.5 to 261.0 | No | ns | 0.9998 |
| CAV2-Cre vs. RVΔL-5Cre | 6.25 | -252.5 to 265.0 | No | ns | 0.9981 |
| AAV2-retro-hSyn-Cre vs. RVΔL-5Cre | 4 | -254.7 to 262.7 | No | ns | 0.9992 |
| L2/3 | | | | | |
| CAV2-Cre vs. AAV2-retro-hSyn-Cre | -393.5 | -652.2 to -134.8 | Yes | ** | 0.0019 |
| CAV2-Cre vs. RVΔL-5Cre | -814 | -1073 to -555.3 | Yes | **** | <0.0001 |
| AAV2-retro-hSyn-Cre vs. RVΔL-5Cre | -420.5 | -679.2 to -161.8 | Yes | *** | 0.0009 |
| L5 | | | | | |
| CAV2-Cre vs. AAV2-retro-hSyn-Cre | 169.8 | -88.96 to 428.5 | No | ns | 0.2571 |
| CAV2-Cre vs. RVΔL-5Cre | -454 | -712.7 to -195.3 | Yes | *** | 0.0004 |
| AAV2-retro-hSyn-Cre vs. RVΔL-5Cre | -623.8 | -882.5 to -365.0 | Yes | **** | <0.0001 |
| L6 | | | | | |
| CAV2-Cre vs. AAV2-retro-hSyn-Cre | -12.5 | -271.2 to 246.2 | No | ns | 0.9923 |
| CAV2-Cre vs. RVΔL-5Cre | -552.8 | -811.5 to -294.0 | Yes | **** | <0.0001 |
| AAV2-retro-hSyn-Cre vs. RVΔL-5Cre | -540.3 | -799.0 to -281.5 | Yes | **** | <0.0001 |

| Test details | Mean 1 | Mean 2 | Mean Diff. | SE of diff. | N1 | N2 | q | DF |
|---|---|---|---|---|---|---|---|---|
| L1 | | | | | | | | |
| CAV2-Cre vs. AAV2-retro-hSyn-Cre | 12.25 | 10 | 2.25 | 105.8 | 4 | 4 | 0.03006 | 36 |
| CAV2-Cre vs. RVΔL-5Cre | 12.25 | 6 | 6.25 | 105.8 | 4 | 4 | 0.08351 | 36 |
| AAV2-retro-hSyn-Cre vs. RVΔL-5Cre | 10 | 6 | 4 | 105.8 | 4 | 4 | 0.05345 | 36 |
| L2/3 | | | | | | | | |
| CAV2-Cre vs. AAV2-retro-hSyn-Cre | 63.5 | 457 | -393.5 | 105.8 | 4 | 4 | 5.258 | 36 |
| CAV2-Cre vs. RVΔL-5Cre | 63.5 | 877.5 | -814 | 105.8 | 4 | 4 | 10.88 | 36 |
| AAV2-retro-hSyn-Cre vs. RVΔL-5Cre | 457 | 877.5 | -420.5 | 105.8 | 4 | 4 | 5.618 | 36 |
| L5 | | | | | | | | |
| CAV2-Cre vs. AAV2-retro-hSyn-Cre | 433.3 | 263.5 | 169.8 | 105.8 | 4 | 4 | 2.268 | 36 |
| CAV2-Cre vs. RVΔL-5Cre | 433.3 | 887.3 | -454 | 105.8 | 4 | 4 | 6.066 | 36 |
| AAV2-retro-hSyn-Cre vs. RVΔL-5Cre | 263.5 | 887.3 | -623.8 | 105.8 | 4 | 4 | 8.334 | 36 |
| L6 | | | | | | | | |
| CAV2-Cre vs. AAV2-retro-hSyn-Cre | 337.3 | 349.8 | -12.5 | 105.8 | 4 | 4 | 0.167 | 36 |
| CAV2-Cre vs. RVΔL-5Cre | 337.3 | 890 | -552.8 | 105.8 | 4 | 4 | 7.385 | 36 |
| AAV2-retro-hSyn-Cre vs. RVΔL-5Cre | 349.8 | 890 | -540.3 | 105.8 | 4 | 4 | 7.218 | 36 |

**Ordinary one-way ANOVA of Primary motor area all layers**

| | |
|---|---|
| Number of families | 1 |
| Number of comparisons per family | 3 |
| Alpha | 0.05 |

| Tukey's multiple comparisons test | Mean Diff. | 95.00% CI of diff. | Below threshold? | Summary | Adjusted P Value | |
|---|---|---|---|---|---|---|
| CAV2-Cre vs. AAV2-retro-hSyn-Cre | -234 | -1221 to 753.3 | No | ns | 0.7906 | A-B |
| CAV2-Cre vs. RVΔL-5Cre | -1815 | -2802 to -827.2 | Yes | ** | 0.0016 | A-C |
| AAV2-retro-hSyn-Cre vs. RVΔL-5Cre | -1581 | -2568 to -593.2 | Yes | ** | 0.004 | B-C |

| Test details | Mean 1 | Mean 2 | Mean Diff. | SE of diff. | n1 | n2 | q | DF |
|---|---|---|---|---|---|---|---|---|
| CAV2-Cre vs. AAV2-retro-hSyn-Cre | 846.3 | 1080 | -234 | 353.6 | 4 | 4 | 0.9359 | 9 |
| CAV2-Cre vs. RVΔL-5Cre | 846.3 | 2661 | -1815 | 353.6 | 4 | 4 | 7.257 | 9 |
| AAV2-retro-hSyn-Cre vs. RVΔL-5Cre | 1080 | 2661 | -1581 | 353.6 | 4 | 4 | 6.321 | 9 |

**AM injections**

| primary somatosensory: contralateral | Animal ID | Section ID | L1 | L2/3 | L4 | L5 | L6 |
|---|---|---|---|---|---|---|---|
| CAV2-Cre | 22073105LJ | sec-4 | 0 | 0 | 0 | 7 | 0 |
| | 22073105LJ | sec-5 | 0 | 1 | 0 | 3 | 0 |
| | 22073105LJ | sec-6 | 0 | 4 | 3 | 79 | 0 |
| | 22073105LJ | sec-7 | 0 | 4 | 4 | 52 | 8 |
| | 22073105LJ | sec-8 | 0 | 4 | 2 | 54 | 4 |
| | 22073105LJ | sec-9 | 0 | 4 | 0 | 45 | 5 |
| | 22073105LJ | sec-10 | 0 | 1 | 2 | 34 | 4 |
| | 22073105LJ | Sum: | 0 | 18 | 11 | 274 | 21 |
| | 22073106LJ | sec-2 | 0 | 1 | 0 | 3 | 0 |
| | 22073106LJ | sec-3 | 0 | 0 | 0 | 2 | 0 |
| | 22073106LJ | sec-4 | 0 | 0 | 0 | 3 | 1 |
| | 22073106LJ | sec-5 | 0 | 3 | 0 | 26 | 10 |
| | 22073106LJ | sec-6 | 0 | 2 | 0 | 50 | 15 |
| | 22073106LJ | sec-7 | 0 | 19 | 0 | 98 | 36 |
| | 22073106LJ | sec-8 | 0 | 40 | 0 | 93 | 41 |
| | 22073106LJ | sec-9 | 0 | 16 | 8 | 104 | 24 |
| | 22073106LJ | sec-10 | 0 | 12 | 2 | 55 | 12 |
| | 22073106LJ | Sum: | 0 | 93 | 10 | 434 | 139 |
| | 22073107LJ | sec-3 | 0 | 0 | 0 | 1 | 3 |
| | 22073107LJ | sec-4 | 0 | 2 | 0 | 2 | 3 |
| | 22073107LJ | sec-5 | 0 | 6 | 1 | 9 | 13 |
| | 22073107LJ | sec-6 | 0 | 17 | 0 | 38 | 42 |
| | 22073107LJ | sec-7 | 0 | 34 | 4 | 133 | 80 |
| | 22073107LJ | sec-8 | 0 | 88 | 4 | 224 | 117 |
| | 22073107LJ | sec-9 | 0 | 66 | 3 | 199 | 96 |
| | 22073107LJ | sec-10 | 0 | 26 | 0 | 66 | 42 |
| | 22073107LJ | Sum: | 0 | 239 | 12 | 672 | 396 |
| | 22073108LJ | sec-3 | 0 | 0 | 0 | 2 | 1 |
| | 22073108LJ | sec-4 | 0 | 1 | 0 | 11 | 0 |
| | 22073108LJ | sec-5 | 0 | 0 | 0 | 6 | 0 |
| | 22073108LJ | sec-6 | 0 | 4 | 0 | 45 | 12 |
| | 22073108LJ | sec-7 | 0 | 0 | 0 | 49 | 7 |
| | 22073108LJ | sec-8 | 0 | 0 | 0 | 1 | 0 |
| | 22073108LJ | sec-9 | 0 | 1 | 0 | 23 | 12 |
| | 22073108LJ | Sum: | 0 | 6 | 0 | 137 | 32 |
| AAV2-retro-hSyn-Cre | 22072205LJ | sec-4 | 0 | 0 | 0 | 3 | 1 |
| | 22072205LJ | sec-5 | 1 | 8 | 2 | 0 | 18 |
| | 22072205LJ | sec-6 | 0 | 66 | 0 | 33 | 64 |
| | 22072205LJ | sec-7 | 0 | 142 | 0 | 71 | 144 |
| | 22072205LJ | sec-8 | 0 | 96 | 0 | 53 | 156 |
| | 22072205LJ | sec-9 | 0 | 160 | 0 | 32 | 190 |
| | 22072205LJ | sec-10 | 0 | 85 | 0 | 18 | 130 |
| | 22072205LJ | Sum: | 1 | 557 | 2 | 210 | 703 |
| | 22072206LJ | sec-4 | 0 | 3 | 0 | 2 | 2 |
| | 22072206LJ | sec-5 | 0 | 47 | 0 | 17 | 5 |
| | 22072206LJ | sec-6 | 0 | 82 | 0 | 30 | 11 |
| | 22072206LJ | sec-7 | 0 | 135 | 0 | 89 | 46 |
| | 22072206LJ | sec-8 | 0 | 61 | 0 | 56 | 55 |
| | 22072206LJ | sec-9 | 0 | 77 | 0 | 2 | 52 |
| | 22072206LJ | sec-10 | 0 | 119 | 0 | 6 | 55 |
| | 22072206LJ | Sum: | 0 | 524 | 0 | 202 | 226 |
| | 22072208LJ | sec-2 | 0 | 1 | 0 | 0 | 6 |
| | 22072208LJ | sec-3 | 0 | 25 | 1 | 0 | 42 |
| | 22072208LJ | sec-4 | 0 | 77 | 2 | 1 | 91 |
| | 22072208LJ | sec-5 | 0 | 144 | 0 | 1 | 136 |
| | 22072208LJ | sec-6 | 0 | 66 | 0 | 24 | 70 |
| | 22072208LJ | sec-7 | 0 | 37 | 0 | 4 | 73 |
| | 22072208LJ | sec-8 | 0 | 42 | 0 | 9 | 46 |
| | 22072208LJ | Sum: | 0 | 392 | 3 | 39 | 464 |
| | 22073109LJ | sec-4 | 0 | 5 | 0 | 0 | 3 |
| | 22073109LJ | sec-5 | 0 | 12 | 0 | 1 | 3 |
| | 22073109LJ | sec-6 | 0 | 40 | 0 | 10 | 5 |
| | 22073109LJ | sec-7 | 0 | 56 | 0 | 44 | 19 |
| | 22073109LJ | sec-8 | 0 | 20 | 0 | 14 | 48 |
| | 22073109LJ | sec-9 | 0 | 37 | 0 | 15 | 78 |
| | 22073109LJ | sec-10 | 0 | 11 | 0 | 5 | 23 |

| | | | | | | | |
|---|---|---|---|---|---|---|---|
| | 22073109LJ | Sum: | 0 | 181 | 0 | 89 | 179 |
| RVΔL-5Cre | 22072105LJ | sec-2 | 0 | 0 | 0 | 0 | 2 |
| | 22072105LJ | sec-3 | 0 | 8 | 0 | 0 | 4 |
| | 22072105LJ | sec-4 | 0 | 47 | 1 | 1 | 18 |
| | 22072105LJ | sec-5 | 0 | 64 | 0 | 44 | 69 |
| | 22072105LJ | sec-6 | 1 | 151 | 0 | 102 | 78 |
| | 22072105LJ | sec-7 | 0 | 190 | 0 | 139 | 149 |
| | 22072105LJ | sec-8 | 0 | 205 | 0 | 109 | 149 |
| | 22072105LJ | sec-9 | 1 | 50 | 0 | 82 | 53 |
| | 22072105LJ | sec-10 | 1 | 105 | 22 | 115 | 50 |
| | 22072105LJ | Sum: | 3 | 820 | 23 | 592 | 572 |
| | 22072106LJ | sec-4 | 0 | 5 | 0 | 4 | 0 |
| | 22072106LJ | sec-5 | 0 | 18 | 0 | 10 | 0 |
| | 22072106LJ | sec-6 | 0 | 33 | 0 | 18 | 6 |
| | 22072106LJ | sec-7 | 0 | 75 | 0 | 46 | 15 |
| | 22072106LJ | sec-8 | 0 | 52 | 0 | 27 | 10 |
| | 22072106LJ | sec-9 | 0 | 82 | 0 | 54 | 27 |
| | 22072106LJ | sec-10 | 0 | 44 | 0 | 38 | 19 |
| | 22072106LJ | sec-11 | 0 | 13 | 0 | 16 | 8 |
| | 22072106LJ | Sum: | 0 | 322 | 0 | 213 | 85 |
| | 22072107LJ | sec-1 | 0 | 0 | 0 | 0 | 1 |
| | 22072107LJ | sec-2 | 0 | 0 | 0 | 0 | 3 |
| | 22072107LJ | sec-3 | 0 | 0 | 0 | 0 | 3 |
| | 22072107LJ | sec-4 | 0 | 2 | 0 | 1 | 3 |
| | 22072107LJ | sec-5 | 0 | 9 | 0 | 17 | 22 |
| | 22072107LJ | sec-6 | 3 | 74 | 0 | 69 | 79 |
| | 22072107LJ | sec-7 | 0 | 149 | 0 | 78 | 100 |
| | 22072107LJ | sec-8 | 0 | 175 | 5 | 105 | 136 |
| | 22072107LJ | sec-9 | 5 | 92 | 22 | 93 | 80 |
| | 22072107LJ | sec-10 | 0 | 65 | 12 | 90 | 54 |
| | 22072107LJ | Sum: | 8 | 566 | 39 | 453 | 481 |
| | 22072108LJ | sec-2 | 0 | 3 | 0 | 0 | 5 |
| | 22072108LJ | sec-3 | 0 | 16 | 0 | 6 | 6 |
| | 22072108LJ | sec-4 | 0 | 33 | 0 | 28 | 24 |
| | 22072108LJ | sec-5 | 0 | 75 | 0 | 47 | 49 |
| | 22072108LJ | sec-6 | 0 | 142 | 0 | 80 | 82 |
| | 22072108LJ | sec-7 | 0 | 157 | 4 | 102 | 162 |
| | 22072108LJ | sec-8 | 0 | 88 | 16 | 76 | 53 |
| | 22072108LJ | sec-9 | 0 | 63 | 4 | 86 | 42 |
| | 22072108LJ | sec-10 | 0 | 54 | 4 | 70 | 44 |
| | 22072108LJ | Sum: | 0 | 631 | 28 | 495 | 467 |

**Summary for AM injections**

| primary somatosensory: contralateral | Animal ID | L1 | L2/3 | L4 | L5 | L6 | Sum of all layers |
|---|---|---|---|---|---|---|---|
| CAV2-Cre | 22073105LJ | 0 | 18 | 11 | 274 | 21 | 324 |
| | 22073106LJ | 0 | 93 | 10 | 434 | 139 | 676 |
| | 22073107LJ | 0 | 239 | 12 | 672 | 396 | 1319 |
| | 22073108LJ | 0 | 6 | 0 | 137 | 32 | 175 |
| AAV2-retro-hSyn-Cre | 22072205LJ | 1 | 557 | 2 | 210 | 703 | 1473 |
| | 22072206LJ | 0 | 524 | 0 | 202 | 226 | 952 |
| | 22072208LJ | 0 | 392 | 3 | 39 | 464 | 898 |
| | 22073109LJ | 0 | 181 | 0 | 89 | 179 | 449 |
| RVΔL-5Cre | 22072105LJ | 3 | 820 | 23 | 592 | 572 | 2010 |
| | 22072106LJ | 0 | 322 | 0 | 213 | 85 | 620 |
| | 22072107LJ | 8 | 566 | 39 | 453 | 481 | 1547 |
| | 22072108LJ | 0 | 631 | 28 | 495 | 467 | 1621 |

**ANOVA of Primary somatosensory: contralateral**

Within each row, compare columns (simple effects within rows)

| | |
|---|---|
| Number of families | 5 |
| Number of comparisons per family | 3 |
| Alpha | 0.05 |

| Tukey's multiple comparisons test | Mean Diff. | 95.00% CI of diff. | Below threshold? | Summary | Adjusted P Value |
|---|---|---|---|---|---|
| L1 | | | | | |
| CAV2-Cre vs. AAV2-retro-hSyn-Cre | -0.25 | -244.4 to 243.9 | No | ns | >0.9999 |
| CAV2-Cre vs. RVΔL-5Cre | -2.75 | -246.9 to 241.4 | No | ns | 0.9996 |
| AAV2-retro-hSyn-Cre vs. RVΔL-5Cre | -2.5 | -246.6 to 241.6 | No | ns | 0.9997 |
| L2/3 | | | | | |
| CAV2-Cre vs. AAV2-retro-hSyn-Cre | -324.5 | -568.6 to -80.37 | Yes | ** | 0.0066 |
| CAV2-Cre vs. RVΔL-5Cre | -495.8 | -739.9 to -251.6 | Yes | **** | <0.0001 |
| AAV2-retro-hSyn-Cre vs. RVΔL-5Cre | -171.3 | -415.4 to 72.88 | No | ns | 0.2163 |
| L4 | | | | | |
| CAV2-Cre vs. AAV2-retro-hSyn-Cre | 7 | -237.1 to 251.1 | No | ns | 0.9973 |
| CAV2-Cre vs. RVΔL-5Cre | -14.25 | -258.4 to 229.9 | No | ns | 0.989 |
| AAV2-retro-hSyn-Cre vs. RVΔL-5Cre | -21.25 | -265.4 to 222.9 | No | ns | 0.9758 |
| L5 | | | | | |
| CAV2-Cre vs. AAV2-retro-hSyn-Cre | 244.3 | 0.1234 to 488.4 | Yes | * | 0.0499 |
| CAV2-Cre vs. RVΔL-5Cre | -59 | -303.1 to 185.1 | No | ns | 0.8284 |
| AAV2-retro-hSyn-Cre vs. RVΔL-5Cre | -303.3 | -547.4 to -59.12 | Yes | * | 0.0117 |
| L6 | | | | | |
| CAV2-Cre vs. AAV2-retro-hSyn-Cre | -246 | -490.1 to -1.873 | Yes | * | 0.0479 |
| CAV2-Cre vs. RVΔL-5Cre | -254.3 | -498.4 to -10.12 | Yes | * | 0.0396 |
| AAV2-retro-hSyn-Cre vs. RVΔL-5Cre | -8.25 | -252.4 to 235.9 | No | ns | 0.9963 |

| Test details | Mean 1 | Mean 2 | Mean Diff. | SE of diff. | N1 | N2 | q | DF |
|---|---|---|---|---|---|---|---|---|
| L1 | | | | | | | | |
| CAV2-Cre vs. AAV2-retro-hSyn-Cre | 0 | 0.25 | -0.25 | 100.7 | 4 | 4 | 0.00351 | 45 |
| CAV2-Cre vs. RVΔL-5Cre | 0 | 2.75 | -2.75 | 100.7 | 4 | 4 | 0.03861 | 45 |
| AAV2-retro-hSyn-Cre vs. RVΔL-5Cre | 0.25 | 2.75 | -2.5 | 100.7 | 4 | 4 | 0.0351 | 45 |
| L2/3 | | | | | | | | |
| CAV2-Cre vs. AAV2-retro-hSyn-Cre | 89 | 413.5 | -324.5 | 100.7 | 4 | 4 | 4.556 | 45 |
| CAV2-Cre vs. RVΔL-5Cre | 89 | 584.8 | -495.8 | 100.7 | 4 | 4 | 6.96 | 45 |
| AAV2-retro-hSyn-Cre vs. RVΔL-5Cre | 413.5 | 584.8 | -171.3 | 100.7 | 4 | 4 | 2.404 | 45 |
| L4 | | | | | | | | |
| CAV2-Cre vs. AAV2-retro-hSyn-Cre | 8.25 | 1.25 | 7 | 100.7 | 4 | 4 | 0.09828 | 45 |
| CAV2-Cre vs. RVΔL-5Cre | 8.25 | 22.5 | -14.25 | 100.7 | 4 | 4 | 0.2001 | 45 |
| AAV2-retro-hSyn-Cre vs. RVΔL-5Cre | 1.25 | 22.5 | -21.25 | 100.7 | 4 | 4 | 0.2983 | 45 |
| L5 | | | | | | | | |
| CAV2-Cre vs. AAV2-retro-hSyn-Cre | 379.3 | 135 | 244.3 | 100.7 | 4 | 4 | 3.429 | 45 |
| CAV2-Cre vs. RVΔL-5Cre | 379.3 | 438.3 | -59 | 100.7 | 4 | 4 | 0.8284 | 45 |
| AAV2-retro-hSyn-Cre vs. RVΔL-5Cre | 135 | 438.3 | -303.3 | 100.7 | 4 | 4 | 4.258 | 45 |
| L6 | | | | | | | | |
| CAV2-Cre vs. AAV2-retro-hSyn-Cre | 147 | 393 | -246 | 100.7 | 4 | 4 | 3.454 | 45 |
| CAV2-Cre vs. RVΔL-5Cre | 147 | 401.3 | -254.3 | 100.7 | 4 | 4 | 3.57 | 45 |
| AAV2-retro-hSyn-Cre vs. RVΔL-5Cre | 393 | 401.3 | -8.25 | 100.7 | 4 | 4 | 0.1158 | 45 |

**Ordinary one-way ANOVA of Primary somatosensory: contralateral (all layers)**

| | |
|---|---|
| Number of families | 1 |
| Number of comparisons per family | 3 |
| Alpha | 0.05 |

| Tukey's multiple comparisons test | Mean Diff. | 95.00% CI of diff. | Below threshold? | Summary | Adjusted P Value | |
|---|---|---|---|---|---|---|
| CAV2-Cre vs. AAV2-retro-hSyn-Cre | -319.5 | -1327 to 688.4 | No | ns | 0.6625 | A-B |
| CAV2-Cre vs. RVΔL-5Cre | -826 | -1834 to 181.9 | No | ns | 0.109 | A-C |
| AAV2-retro-hSyn-Cre vs. RVΔL-5Cre | -506.5 | -1514 to 501.4 | No | ns | 0.3795 | B-C |

| Test details | Mean 1 | Mean 2 | Mean Diff. | SE of diff. | n1 | n2 | q | DF |
|---|---|---|---|---|---|---|---|---|
| CAV2-Cre vs. AAV2-retro-hSyn-Cre | 623.5 | 943 | -319.5 | 361 | 4 | 4 | 1.252 | 9 |
| CAV2-Cre vs. RVΔL-5Cre | 623.5 | 1450 | -826 | 361 | 4 | 4 | 3.236 | 9 |
| AAV2-retro-hSyn-Cre vs. RVΔL-5Cre | 943 | 1450 | -506.5 | 361 | 4 | 4 | 1.984 | 9 |

**Supplementary Video S1. RV∆L-Cre-labeled cortical neurons at 2 weeks vs 10 weeks postinjection, Related to Figure 4.**
3-dimensional rendering of the same cortical volume shown in Figure 4, panel B.

*See external file.*

**Supplementary File S6. Cell counts and statistics for structural two-photon imaging experiments, Related to Figure 4.**
Cell counts and statistical comparisons of neurons labeled by either ∆GL or ∆L RV.

*See following pages.*

| RVΔGL-Cre: cell counts | Animal-1 FOV-1 | Animal-1 FOV-2 | Animal-2 FOV-1 | Animal-2 FOV-2 | Animal-3 FOV-1 | Animal-3 FOV-2 | Animal-4 FOV-1 | Animal-4 FOV-2 |
|---|---|---|---|---|---|---|---|---|
| Week-1 | 300 | 315 | 219 | 318 | 435 | 288 | 203 | 220 |
| Week-2 | 519 | 359 | 269 | 546 | 488 | 364 | 321 | 354 |
| Week-3 | 537 | 390 | 305 | 569 | 505 | 443 | 369 | 427 |
| Week-4 | 538 | 403 | 317 | 572 | 518 | 445 | 372 | 426 |
| Week-6 | 537 | 403 | 319 | 571 | 515 | 451 | 370 | 424 |
| Week-12 | 535 | 400 | 319 | 567 | 514 | 443 | 369 | 424 |
| Week-14 | | | 319 | 567 | 513 | 442 | 368 | 421 |
| Week-16 | | | 319 | | | | | |

| RVΔGL-Cre: percentage to week-1 | Animal-1 FOV-1 | Animal-1 FOV-2 | Animal-2 FOV-1 | Animal-2 FOV-2 | Animal-3 FOV-1 | Animal-3 FOV-2 | Animal-4 FOV-1 | Animal-4 FOV-2 |
|---|---|---|---|---|---|---|---|---|
| Week-1 | 1.00 | 1.00 | 1.00 | 1.00 | 1.00 | 1.00 | 1.00 | 1.00 |
| Week-2 | 1.73 | 1.14 | 1.23 | 1.72 | 1.12 | 1.26 | 1.58 | 1.61 |
| Week-3 | 1.79 | 1.24 | 1.39 | 1.79 | 1.16 | 1.54 | 1.82 | 1.94 |
| Week-4 | 1.79 | 1.28 | 1.45 | 1.80 | 1.19 | 1.55 | 1.83 | 1.94 |
| Week-6 | 1.79 | 1.28 | 1.46 | 1.80 | 1.18 | 1.57 | 1.82 | 1.93 |
| Week-12 | 1.78 | 1.27 | 1.46 | 1.78 | 1.18 | 1.54 | 1.82 | 1.93 |
| Week-14 | | | 1.46 | 1.78 | 1.18 | 1.53 | 1.81 | 1.91 |
| Week-16 | | | 1.46 | | | | | |

| RVΔGL-Cre: comparison between week-1 and week-4 | Animal-1 FOV-1 | Animal-1 FOV-2 | Animal-2 FOV-1 | Animal-2 FOV-2 | Animal-3 FOV-1 | Animal-3 FOV-2 | Animal-4 FOV-1 | Animal-4 FOV-2 |
|---|---|---|---|---|---|---|---|---|
| Week-1 | 300 | 315 | 219 | 318 | 435 | 288 | 203 | 220 |
| Week-4 | 538 | 403 | 317 | 572 | 518 | 445 | 372 | 426 |
| Increase (Week-4 - Week-1) | 238 | 88 | 98 | 254 | 83 | 157 | 169 | 206 |
| % increase (Week-4 - Week-1) | 79.33% | 27.94% | 44.75% | 79.87% | 19.08% | 54.51% | 83.25% | 93.64% |
| Increase (Week-12 - Week-4) | -3 | -3 | 2 | -5 | -4 | -2 | -3 | -2 |
| % increase (Week-12 - Week-4) | -0.56% | -0.74% | 0.63% | -0.87% | -0.77% | -0.45% | -0.81% | -0.47% |

| t-Test: Paired Two Sample for Means | | |
|---|---|---|
| | Week-1 | Week-4 |
| Mean | 287.25 | 448.875 |
| Variance | 5712.5 | 7689.267857 |
| Observations | 8 | 8 |
| Pearson Correlation | 0.665513457 | |
| Hypothesized Mean Difference | 0 | |
| df | 7 | |
| t Stat | -6.75473154 | |
| P(T<=t) one-tail | **0.000131934** | |
| t Critical one-tail | 1.894578605 | |
| P(T<=t) two-tail | 0.000263868 | |
| t Critical two-tail | 2.364624252 | |

| t-Test: Paired Two Sample for Means | | |
|---|---|---|
| | Week-4 | Week-12 |
| Mean | 448.875 | 446.375 |
| Variance | 7689.267857 | 7407.410714 |
| Observations | 8 | 8 |
| Pearson Correlation | 0.999890399 | |
| Hypothesized Mean Difference | 0 | |
| df | 7 | |
| t Stat | 3.415650255 | |
| P(T<=t) one-tail | **0.005600716** | |
| t Critical one-tail | 1.894578605 | |
| P(T<=t) two-tail | 0.011201433 | |
| t Critical two-tail | 2.364624252 | |

| % increase from 1 week to 4 weeks: | | | | | | | | |
|---|---|---|---|---|---|---|---|---|
| **RVΔGL-Cre** | 79.33% | 27.94% | 44.75% | 79.87% | 19.08% | 54.51% | 83.25% | 93.64% |
| **RVΔL-Cre** | 84.38% | 84.41% | 49.57% | 70.25% | 54.88% | 67.66% | 40.66% | 93.26% |

| Unpaired t test | |
|---|---|
| Table Analyzed | % increase |
| P value | **0.5187** |
| P value summary | ns |
| Significantly different (P < 0.05)? | No |
| One- or two-tailed P value? | Two-tailed |
| t, df | t=0.6621, df=14 |
| **How big is the difference?** | |
| Mean of RVΔGL-Cre | 0.603 |
| Mean of RVΔL-Cre | 0.6813 |
| Difference between means (B - A) ± SEM | 0.07836 ± 0.1184 |
| 95% confidence interval | -0.1755 to 0.3322 |
| R squared (eta squared) | 0.03036 |
| **F test to compare variances** | |
| F, DFn, Dfd | 2.216, 7, 7 |
| P value | 0.3158 |
| P value summary | ns |
| Significantly different (P < 0.05)? | No |
| **Data analyzed** | |
| Sample size, RVΔGL-Cre | 8 |
| Sample size, RVΔL-Cre | 8 |

| % increase from 4 weeks to 12 weeks: | | | | | | | | |
|---|---|---|---|---|---|---|---|---|
| **RVΔGL-Cre** | -0.56% | -0.74% | 0.63% | -0.87% | -0.77% | -0.45% | -0.81% | -0.47% |
| **RVΔL-Cre** | 0.48% | -0.29% | 2.31% | -1.46% | 3.30% | 0.00% | 0.78% | 0.58% |

| Unpaired t test | |
|---|---|
| P value | **0.0451** |
| P value summary | * |
| Significantly different (P < 0.05)? | Yes |
| One- or two-tailed P value? | Two-tailed |
| t, df | t=2.199, df=14 |
| **How big is the difference?** | |
| Mean of RVΔGL-Cre | -0.005054 |
| Mean of RVΔL-Cre | 0.007135 |
| Difference between means (B - A) ± SEM | .01219 ± 0.005542 |
| 95% confidence interval | 0003030 to 0.02407 |
| R squared (eta squared) | 0.2568 |
| **F test to compare variances** | |
| F, DFn, Dfd | 9.402, 7, 7 |
| P value | 0.0085 |
| P value summary | ** |
| Significantly different (P < 0.05)? | Yes |
| Data analyzed | |
| Sample size, RVΔGL-Cre | 8 |
| Sample size, RVΔL-Cre | 8 |

| RV∆L-Cre: cell counts | Animal-1 FOV-1 | Animal-1 FOV-2 | Animal-2 FOV-1 | Animal-2 FOV-2 | Animal-3 FOV-1 | Animal-3 FOV-2 | Animal-4 FOV-1 | Animal-4 FOV-2 |
|---|---|---|---|---|---|---|---|---|
| Week-1 | 224 | 186 | 232 | 242 | 215 | 167 | 182 | 178 |
| Week-2 | 331 | 307 | 300 | 298 | 270 | 237 | 200 | 228 |
| Week-3 | 396 | 340 | 341 | 378 | 289 | 279 | 253 | 304 |
| Week-4 | 413 | 343 | 347 | 412 | 333 | 280 | 256 | 344 |
| Week-6 | 409 | 342 | 347 | 413 | 342 | 283 | 255 | 345 |
| Week-12 | 415 | 342 | 355 | 406 | 344 | 280 | 258 | 346 |
| Week 14 | 415 | 342 | 354 | 406 | 344 | 280 | | |
| Week-16 | | | | 406 | 344 | 280 | | |

| RV∆L-Cre: percentage to week-1 | Animal-1 FOV-1 | Animal-1 FOV-2 | Animal-2 FOV-1 | Animal-2 FOV-2 | Animal-3 FOV-1 | Animal-3 FOV-2 | Animal-4 FOV-1 | Animal-4 FOV-2 |
|---|---|---|---|---|---|---|---|---|
| Week-1 | 1.00 | 1.00 | 1.00 | 1.00 | 1.00 | 1.00 | 1.00 | 1.00 |
| Week-2 | 1.48 | 1.65 | 1.29 | 1.23 | 1.26 | 1.42 | 1.10 | 1.28 |
| Week-3 | 1.77 | 1.83 | 1.47 | 1.56 | 1.34 | 1.67 | 1.39 | 1.71 |
| Week-4 | 1.84 | 1.84 | 1.50 | 1.70 | 1.55 | 1.68 | 1.41 | 1.93 |
| Week-6 | 1.83 | 1.84 | 1.50 | 1.71 | 1.59 | 1.69 | 1.40 | 1.94 |
| Week-12 | 1.85 | 1.84 | 1.53 | 1.68 | 1.60 | 1.68 | 1.42 | 1.94 |
| Week 14 | 1.85 | 1.84 | 1.53 | 1.68 | 1.60 | 1.68 | | |
| Week-16 | | | | 1.68 | 1.60 | 1.68 | | |

| RV∆L-Cre: comparison between week-1 and week-4 | Animal-1 FOV-1 | Animal-1 FOV-2 | Animal-2 FOV-1 | Animal-2 FOV-2 | Animal-3 FOV-1 | Animal-3 FOV-2 | Animal-4 FOV-1 | Animal-4 FOV-2 |
|---|---|---|---|---|---|---|---|---|
| Week-1 | 224 | 186 | 232 | 242 | 215 | 167 | 182 | 178 |
| Week-4 | 413 | 343 | 347 | 412 | 333 | 280 | 256 | 344 |
| Increase (Week-4 - Week-1) | 189 | 157 | 115 | 170 | 118 | 113 | 74 | 166 |
| % increase (Week-4 - Week-1) | 84.38% | 84.41% | 49.57% | 70.25% | 54.88% | 67.66% | 40.66% | 93.26% |
| Increase (Week-12 - Week-4) | 2 | -1 | 8 | -6 | 11 | 0 | 2 | 2 |
| % increase (Week-12 - Week-4) | 0.48% | -0.29% | 2.31% | -1.46% | 3.30% | 0.00% | 0.78% | 0.58% |

| t-Test: Paired Two Sample for Means | | |
|---|---|---|
| | **Week-1** | **Week-4** |
| Mean | 203.25 | 341 |
| Variance | 799.6428571 | 3040.571429 |
| Observations | 8 | 8 |
| Pearson Correlation | 0.754099973 | |
| Hypothesized Mean Difference | 0 | |
| df | 7 | |
| t Stat | -10.09862351 | |
| P(T<=t) one-tail | **1.00266E-05** | |
| t Critical one-tail | 1.894578605 | |
| P(T<=t) two-tail | 2.00531E-05 | |
| t Critical two-tail | 2.364624252 | |

| t-Test: Paired Two Sample for Means | | |
|---|---|---|
| | **Week-4** | **Week-12** |
| Mean | 341 | 343.25 |
| Variance | 3040.571429 | 2928.785714 |
| Observations | 8 | 8 |
| Pearson Correlation | 0.995543784 | |
| Hypothesized Mean Difference | 0 | |
| df | 7 | |
| t Stat | -1.210419877 | |
| P(T<=t) one-tail | **0.13269902** | |
| t Critical one-tail | 1.894578605 | |
| P(T<=t) two-tail | 0.265398039 | |
| t Critical two-tail | 2.364624252 | |

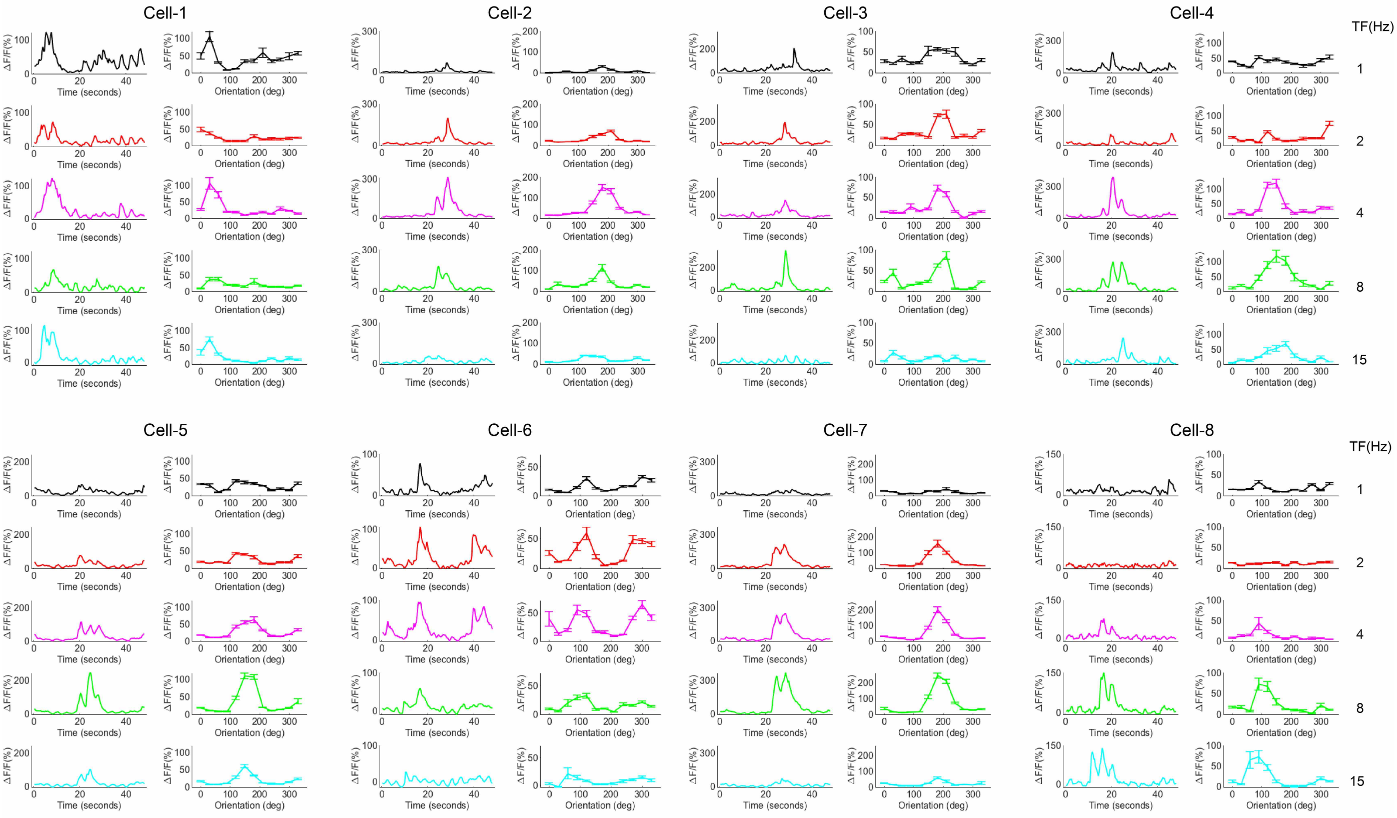

**Supplementary Figure S12. GCaMP6s signals and tuning curves of 8 example V1 neurons at 16 weeks postinjection, Related to Figure 6.**

These data were obtained with drifting gratings presented at 12 directions of motion and 5 temporal frequencies, repeated 10 times (tuning curve: mean ΔF/F ± s.e.m; GCaMP6s signals: mean ΔF/F) at two different FOVs at the 16-week timepoint.

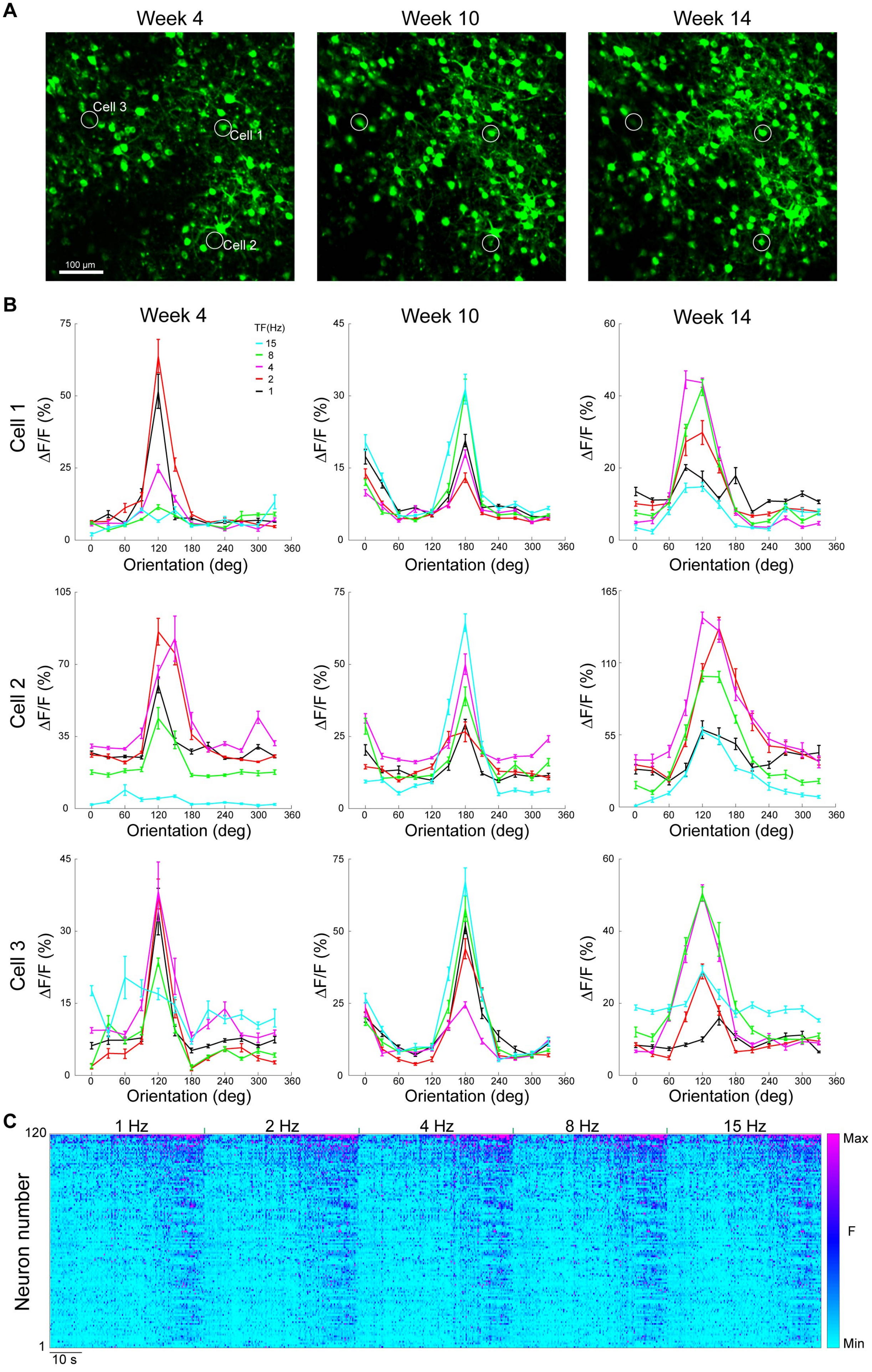
A
Week 4
Week 10
Week 14
Cell 3
Cell 1
Cell 2
100 µm
B
Week 4
Week 10
Week 14
TF(Hz)
15
8
4
2
1
Cell 1
Cell 2
Cell 3
ΔF/F (%)
Orientation (deg)
C
1 Hz
2 Hz
4 Hz
8 Hz
15 Hz
Neuron number
10 s
Max
F
Min

**Supplementary Figure S13. More examples showing long-term stability of orientation and temporal frequency tuning in RV∆L-Cre labeled neurons, Related to Figure 6.**
(A) Maximum intensity projections of the same FOV at 4 weeks, 10 weeks, and 14 weeks postinjection. Scale bar: 100 μm, applies to all images.
(B) Visual responses of the three circled cells in panel **a** measured at the three different timepoints. Data were obtained with drifting gratings presented at 12 directions of motion and 5 temporal frequencies (mean ΔF/F ± s.e.m., averaged over 10 repeats).
(C) Single-cell fluorescence time courses for 120 cells at the 12-week timepoint, showing activity over all five temporal frequencies (mean ΔF/F, averaged over 10 repeats). Cells are ranked in descending order of total activity. Scale bar: 10 s.

**Supplementary Video S2. GCaMP6s signals of visual cortical neurons 16 weeks after injection of RV∆L-Cre, Related to Figure 6**.
Video shows responses to 10 repeats of drifting gratings at 12 directions of motion at one temporal frequency (1 Hz) over a total of 480 seconds.

*See external file.*

**Supplementary File S7. Tuned cell counts and statistics for functional two-photon imaging experiments, Related to Figure 6.**
Counts, percentages of tuned cells at weeks 2 to 14, statistical comparison of percentage tuned between weeks 2 and 14, and counts and percentages of cells tuned at 4 weeks that were identifiable at 14 weeks.

*See following pages.*

| | Animals ID | FOVs | Total cell numbers | Total tuning cells numbers | Percentage |
|---|---|---|---|---|---|
| Week-2 | 18121201LJ | FOV-1 | 39 | 11 | 28.21% |
| | | FOV-2 | 43 | 14 | 32.56% |
| | 18121204LJ | FOV-1 | 97 | 62 | 63.92% |
| | | FOV-2 | 92 | 47 | 51.09% |
| | 18122802LJ | FOV-1 | 59 | 21 | 35.59% |
| | | FOV-2 | 149 | 75 | 50.34% |
| Week-3 | 18121201LJ | FOV-1 | 61 | 21 | 34.43% |
| | | FOV-2 | 44 | 14 | 31.82% |
| | 18121204LJ | FOV-1 | 67 | 42 | 62.69% |
| | | FOV-2 | 86 | 51 | 59.30% |
| | 18122802LJ | FOV-1 | 53 | 25 | 47.17% |
| | | FOV-2 | 115 | 59 | 51.30% |
| Week-4 | 18121201LJ | FOV-1 | 40 | 15 | 37.50% |
| | | FOV-2 | 41 | 16 | 39.02% |
| | 18121204LJ | FOV-1 | 84 | 50 | 59.52% |
| | | FOV-2 | 87 | 57 | 65.52% |
| | 18122802LJ | FOV-1 | 51 | 28 | 54.90% |
| | | FOV-2 | 145 | 82 | 56.55% |
| Week-10 | 18121201LJ | FOV-1 | 68 | 26 | 38.24% |
| | | FOV-2 | 40 | 18 | 45.00% |
| | 18121204LJ | FOV-1 | 70 | 43 | 61.43% |
| | | FOV-2 | 67 | 45 | 67.16% |
| | 18122802LJ | FOV-1 | 74 | 44 | 59.46% |
| | | FOV-2 | 199 | 108 | 54.27% |
| Week-12 | 18121201LJ | FOV-1 | 64 | 28 | 43.75% |
| | | FOV-2 | 59 | 26 | 44.07% |
| | 18121204LJ | FOV-1 | 69 | 44 | 63.77% |
| | | FOV-2 | 100 | 69 | 69.00% |
| | 18122802LJ | FOV-1 | 77 | 49 | 63.64% |
| | | FOV-2 | 214 | 117 | 54.67% |
| Week-14 | 18121201LJ | FOV-1 | 44 | 19 | 43.18% |
| | | FOV-2 | 50 | 22 | 44.00% |
| | 18121204LJ | FOV-1 | 83 | 51 | 61.45% |
| | | FOV-2 | 83 | 60 | 72.29% |
| | 18122802LJ | FOV-1 | 81 | 52 | 64.20% |
| | | FOV-2 | 153 | 84 | 54.90% |

| Animals ID | FOVs | Week-2: Percentage | Week-14: Percentage |
|---|---|---|---|
| 18121201LJ | FOV-1 | 28.21% | 43.18% |
| | FOV-2 | 32.56% | 44.00% |
| 18121204LJ | FOV-1 | 63.92% | 61.45% |
| | FOV-2 | 51.09% | 72.29% |
| 18122802LJ | FOV-1 | 35.59% | 64.20% |
| | FOV-2 | 50.34% | 54.90% |

| t-Test: Paired Two Sample for Means | | |
|---|---|---|
| | Week-2 | Week-14 |
| Mean | 0.436160901 | 0.566693749 |
| Variance | 0.01874512 | 0.013380316 |
| Observations | 6 | 6 |
| Pearson Correlation | 0.617965951 | |
| Hypothesized Mean Difference | 0 | |
| df | 5 | |
| t Stat | -2.853925894 | |
| P(T<=t) one-tail | **0.017829635** | |
| t Critical one-tail | 2.015048373 | |
| P(T<=t) two-tail | 0.03565927 | |
| t Critical two-tail | 2.570581836 | |

| Mouse ID | FOV number | Cells at week-4 | Same cells still tuned at week-14 | Percentage |
|---|---|---|---|---|
| 18121201LJ | FOV-1 | 16 | 15 | 0.9375 |
| 18121201LJ | FOV-2 | 18 | 17 | 0.944444444 |
| 18121204LJ | FOV-1 | 20 | 19 | 0.95 |
| 18121204LJ | FOV-2 | 24 | 21 | 0.875 |
| 18122802LJ | FOV-1 | 31 | 30 | 0.967741935 |
| 18122802LJ | FOV-2 | 55 | 52 | 0.945454545 |
| | | | Mean: | 0.936690154 |

**Supplementary File S8. Information on mice used for experiments.**

Information on all individual mice used in this study: ID #, strain, genotype, age, weight, health status, whether involved in previous procedures, housing (single vs group).

*See following page.*

| Animal Number | Experiment | Strain | Genotype | Sex | Involvement in previous procedures | Age at time of injection (weeks) | Weight at time of injection (g) | Health status | Housing after surgery | Age at time of perfusion (weeks) |
|---|---|---|---|---|---|---|---|---|---|---|
| 21021505LJ | Figure 2 | C57BL6/J background | Ai65F het | Male | None | 16 | 26.5 | Good | Group | 17 |
| 21021506LJ | Figure 2 | C57BL6/J background | Ai65F het | Female | None | 16 | 26.5 | Good | Single | 17 |
| 21021507LJ | Figure 2 | C57BL6/J background | Ai65F het | Male | None | 16 | 27.3 | Good | Group | 17 |
| 21021508LJ | Figure 2 | C57BL6/J background | Ai65F het | Female | None | 14 | 25.7 | Good | Single | 15 |
| 21052505LJ | Figure 2 | C57BL6/J background | Ai65F het | Female | None | 16 | 21.1 | Good | Group | 17 |
| 21052506LJ | Figure 2 | C57BL6/J background | Ai65F het | Female | None | 11 | 23.1 | Good | Group | 12 |
| 21052507LJ | Figure 2 | C57BL6/J background | Ai65F het | Female | None | 11 | 19.8 | Good | Group | 12 |
| 21052508LJ | Figure 2 | C57BL6/J background | Ai65F het | Female | None | 11 | 20.7 | Good | Group | 12 |
| 21021805LJ | Figure 2 | C57BL6/J background | Ai65F het | Male | None | 17 | 35.5 | Good | Single | 18 |
| 21021806LJ | Figure 2 | C57BL6/J background | Ai65F het | Female | None | 13 | 25.8 | Good | Group | 14 |
| 21021807LJ | Figure 2 | C57BL6/J background | Ai65F het | Female | None | 13 | 20.4 | Good | Group | 14 |
| 21021808LJ | Figure 2 | C57BL6/J background | Ai65F het | Male | None | 10 | 28.6 | Good | Single | 11 |
| 21052607LJ | Figure 2 | C57BL6/J background | Ai65F het | Male | None | 11 | 29.3 | Good | Group | 12 |
| 21052608LJ | Figure 2 | C57BL6/J background | Ai65F het | Male | None | 11 | 29.3 | Good | Group | 12 |
| 21060302LJ | Figure 2 | C57BL6/J background | Ai65F het | Male | None | 8 | 24 | Good | Single | 9 |
| 21060303LJ | Figure 2 | C57BL6/J background | Ai65F het | Female | None | 31 | 24.2 | Good | Single | 32 |
| 21021501LJ | Figure 2 | C57BL6/J background | Ai65F het | Female | None | 14 | 21.8 | Good | Group | 18 |
| 21021502LJ | Figure 2 | C57BL6/J background | Ai65F het | Female | None | 14 | 31.5 | Good | Group | 18 |
| 21021503LJ | Figure 2 | C57BL6/J background | Ai65F het | Male | None | 13 | 24.5 | Good | Group | 17 |
| 21021504LJ | Figure 2 | C57BL6/J background | Ai65F het | Male | None | 13 | 29.4 | Good | Group | 17 |
| 21052501LJ | Figure 2 | C57BL6/J background | Ai65F het | Female | None | 11 | 20.4 | Good | Group | 15 |
| 21052502LJ | Figure 2 | C57BL6/J background | Ai65F het | Female | None | 11 | 21.2 | Good | Group | 15 |
| 21052503LJ | Figure 2 | C57BL6/J background | Ai65F het | Female | None | 11 | 20.9 | Good | Group | 15 |
| 21052504LJ | Figure 2 | C57BL6/J background | Ai65F het | Female | None | 16 | 20.4 | Good | Group | 20 |
| 21021801LJ | Figure 2 | C57BL6/J background | Ai65F het | Male | None | 7 | 23.8 | Good | Group | 11 |
| 21021802LJ | Figure 2 | C57BL6/J background | Ai65F het | Male | None | 7 | 23.6 | Good | Group | 11 |
| 21021803LJ | Figure 2 | C57BL6/J background | Ai65F het | Male | None | 7 | 25.1 | Good | Group | 11 |
| 21021804LJ | Figure 2 | C57BL6/J background | Ai65F het | Male | None | 7 | 25.3 | Good | Group | 11 |
| 21052601LJ | Figure 2 | C57BL6/J background | Ai65F het | Male | None | 11 | 28.6 | Good | Group | 15 |
| 21052602LJ | Figure 2 | C57BL6/J background | Ai65F het | Male | None | 11 | 28.5 | Good | Group | 15 |
| 21052603LJ | Figure 2 | C57BL6/J background | Ai65F het | Male | None | 11 | 24.5 | Good | Group | 15 |
| 21052606LJ | Figure 2 | C57BL6/J background | Ai65F het | Male | None | 11 | 25.7 | Good | Single | 15 |
| 21021703LJ | Figure 2 | C57BL6/J background | Ai14 het | Female | None | 13 | 21.4 | Good | Group | 14 |
| 21021704LJ | Figure 2 | C57BL6/J background | Ai14 het | Female | None | 13 | 20.6 | Good | Group | 14 |
| 21021707LJ | Figure 2 | C57BL6/J background | Ai14 het | Male | None | 12 | 27 | Good | Group | 13 |
| 21021708LJ | Figure 2 | C57BL6/J background | Ai14 het | Male | None | 12 | 24.9 | Good | Group | 13 |
| 21021905LJ | Figure 2 | C57BL6/J background | Ai14 het | Female | None | 7 | 17.7 | Good | Group | 8 |
| 21021906LJ | Figure 2 | C57BL6/J background | Ai14 het | Female | None | 7 | 18.2 | Good | Group | 8 |
| 21021907LJ | Figure 2 | C57BL6/J background | Ai14 het | Female | None | 7 | 18.6 | Good | Group | 8 |
| 21021908LJ | Figure 2 | C57BL6/J background | Ai14 het | Male | None | 12 | 27 | Good | Single | 13 |
| 21021701LJ | Figure 2 | C57BL6/J background | Ai14 het | Female | None | 12 | 19.8 | Good | Single | 16 |
| 21021702LJ | Figure 2 | C57BL6/J background | Ai14 het | Female | None | 13 | 22.8 | Good | Group | 17 |
| 21021705LJ | Figure 2 | C57BL6/J background | Ai14 het | Male | None | 12 | 28.6 | Good | Group | 16 |
| 21021706LJ | Figure 2 | C57BL6/J background | Ai14 het | Male | None | 12 | 25.5 | Good | Group | 16 |
| 21021901LJ | Figure 2 | C57BL6/J background | Ai14 het | Female | None | 8 | 19.6 | Good | Group | 12 |
| 21021902LJ | Figure 2 | C57BL6/J background | Ai14 het | Female | None | 8 | 17.9 | Good | Group | 12 |
| 21021903LJ | Figure 2 | C57BL6/J background | Ai14 het | Male | None | 8 | 26 | Good | Group | 12 |
| 21021904LJ | Figure 2 | C57BL6/J background | Ai14 het | Male | None | 8 | 25.1 | Good | Group | 12 |
| 18080801NL | Figure 2 | C57BL6/J background | Ai14 het | Male | None | 6 | 16.9 | Good | Single | 23 |
| 18080701NL | Figure 2 | C57BL6/J background | Ai14 het | Female | None | 6 | 16.5 | Good | Group | 32 |
| 18080702NL | Figure 2 | C57BL6/J background | Ai14 het | Female | None | 6 | 15.6 | Good | Group | 32 |
| 21042901LJ | Figure S4 | C57BL6/J background | Ai63 het | Female | None | 14 | 23.1 | Good | Group | 15 |
| 21042902LJ | Figure S4 | C57BL6/J background | Ai63 het | Female | None | 14 | 21.6 | Good | Group | 15 |
| 21042903LJ | Figure S4 | C57BL6/J background | Ai63 het | Female | None | 14 | 23.8 | Good | Group | 15 |
| 21060301LJ | Figure S4 | C57BL6/J background | Ai63 het | Male | None | 19 | 35 | Good | Single | 20 |
| 21042905LJ | Figure S4 | C57BL6/J background | Ai63 het | Female | None | 8 | 20.3 | Good | Group | 12 |
| 21042906LJ | Figure S4 | C57BL6/J background | Ai63 het | Female | None | 8 | 19 | Good | Group | 12 |
| 21042907LJ | Figure S4 | C57BL6/J background | Ai63 het | Male | None | 8 | 21.5 | Good | Single | 12 |
| 21042908LJ | Figure S4 | C57BL6/J background | Ai63 het | Male | None | 12 | 26.7 | Good | Single | 16 |
| 22073101LJ | Figure 3 | C57BL6/J background | Ai14 het | Female | None | 11 | 22.8 | Good | Group | 15 |
| 22073102LJ | Figure 3 | C57BL6/J background | Ai14 het | Female | None | 11 | 23.1 | Good | Group | 15 |
| 22073103LJ | Figure 3 | C57BL6/J background | Ai14 het | Female | None | 11 | 20.8 | Good | Group | 15 |
| 22073104LJ | Figure 3 | C57BL6/J background | Ai14 het | Female | None | 11 | 22.2 | Good | Group | 15 |
| 22072201LJ | Figure 3 | C57BL6/J background | Ai14 het | Female | None | 23 | 21.8 | Good | Group | 27 |
| 22072202LJ | Figure 3 | C57BL6/J background | Ai14 het | Female | None | 23 | 21.1 | Good | Group | 27 |
| 22072203LJ | Figure 3 | C57BL6/J background | Ai14 het | Female | None | 23 | 19.2 | Good | Group | 27 |
| 22072204LJ | Figure 3 | C57BL6/J background | Ai14 het | Female | None | 23 | 24.2 | Good | Group | 27 |
| 22072101LJ | Figure 3 | C57BL6/J background | Ai14 het | Female | None | 18 | 21.4 | Good | Group | 22 |
| 22072102LJ | Figure 3 | C57BL6/J background | Ai14 het | Female | None | 18 | 22.4 | Good | Group | 22 |
| 22072103LJ | Figure 3 | C57BL6/J background | Ai14 het | Female | None | 18 | 20.1 | Good | Group | 22 |
| 22072104LJ | Figure 3 | C57BL6/J background | Ai14 het | Female | None | 18 | 22.8 | Good | Group | 22 |
| 22073105LJ | Figure S5 | C57BL6/J background | Ai14 het | Female | None | 11 | 21 | Good | Group | 15 |
| 22073106LJ | Figure S5 | C57BL6/J background | Ai14 het | Female | None | 11 | 21.5 | Good | Group | 15 |
| 22073107LJ | Figure S5 | C57BL6/J background | Ai14 het | Female | None | 11 | 22.7 | Good | Group | 15 |
| 22073108LJ | Figure S5 | C57BL6/J background | Ai14 het | Female | None | 10 | 22.3 | Good | Single | 14 |
| 22072205LJ | Figure S5 | C57BL6/J background | Ai14 het | Female | None | 22 | 20.4 | Good | Group | 26 |
| 22072206LJ | Figure S5 | C57BL6/J background | Ai14 het | Female | None | 22 | 20.6 | Good | Group | 26 |
| 22072208LJ | Figure S5 | C57BL6/J background | Ai14 het | Male | None | 24 | 27.3 | Good | Single | 28 |
| 22073109LJ | Figure S5 | C57BL6/J background | Ai14 het | Female | None | 10 | 20.5 | Good | Single | 14 |
| 22072105LJ | Figure S5 | C57BL6/J background | Ai14 het | Female | None | 18 | 24.1 | Good | Group | 22 |
| 22072106LJ | Figure S5 | C57BL6/J background | Ai14 het | Female | None | 18 | 23.4 | Good | Group | 22 |
| 22072107LJ | Figure S5 | C57BL6/J background | Ai14 het | Female | None | 18 | 24.2 | Good | Group | 22 |
| 22072108LJ | Figure S5 | C57BL6/J background | Ai14 het | Female | None | 18 | 20.3 | Good | Group | 22 |
| 18080601LJ | Figure 4 | C57BL6/J background | Ai14 het | Male | None | 9 | 21.8 | Good | Single | (Longitudinal structural two-photon imaging) |
| 18080602LJ | Figure 4 | C57BL6/J background | Ai14 het | Female | None | 8 | 22.8 | Good | Single | (Longitudinal structural two-photon imaging) |
| 18080603LJ | Figure 4 | C57BL6/J background | Ai14 het | Female | None | 8 | 19.1 | Good | Single | (Longitudinal structural two-photon imaging) |
| 18080604LJ | Figure 4 | C57BL6/J background | Ai14 het | Female | None | 8 | 20.4 | Good | Single | (Longitudinal structural two-photon imaging) |
| 1808701LJ | Figure 4 | C57BL6/J background | Ai14 het | Female | None | 8 | 19.1 | Good | Single | (Longitudinal structural two-photon imaging) |
| 18080802LJ | Figure 4 | C57BL6/J background | Ai14 het | Male | None | 6 | 22.4 | Good | Single | (Longitudinal structural two-photon imaging) |
| 18080803LJ | Figure 4 | C57BL6/J background | Ai14 het | Male | None | 6 | 17.2 | Good | Single | (Longitudinal structural two-photon imaging) |
| 18080802TL | Figure 4 | C57BL6/J background | Ai14 het | Male | None | 6 | 20.2 | Good | Single | (Longitudinal structural two-photon imaging) |
| 22060301LJ | Figure 5 | C57BL6/J background | Ai14 het | Female | None | 10 | 22 | Good | Group | 22 |
| 22060302LJ | Figure 5 | C57BL6/J background | Ai14 het | Female | None | 10 | 22.6 | Good | Group | 22 |
| 22060303LJ | Figure 5 | C57BL6/J background | Ai14 het | Female | None | 10 | 22.6 | Good | Group | 22 |
| 22060304LJ | Figure 5 | C57BL6/J background | Ai14 het | Female | None | 10 | 21 | Good | Group | 22 |
| 22060305LJ | Figure 5 | C57BL6/J background | Ai14 het | Female | None | 11 | 22.1 | Good | Group | 23 |
| 22060306LJ | Figure 5 | C57BL6/J background | Ai14 het | Female | None | 11 | 21.6 | Good | Group | 23 |
| 22060307LJ | Figure 5 | C57BL6/J background | Ai14 het | Female | None | 8 | 21.6 | Good | Single | 20 |
| 22060308LJ | Figure 5 | C57BL6/J background | Ai14 het | Female | None | 8 | 19 | Good | Single | 20 |
| 22071701LJ | Figure 5 | C57BL6/J background | Ai14 het | Female | None | 17 | 21.8 | Good | Group | 21 |
| 22071702LJ | Figure 5 | C57BL6/J background | Ai14 het | Female | None | 17 | 23.2 | Good | Group | 21 |
| 22071703LJ | Figure 5 | C57BL6/J background | Ai14 het | Female | None | 17 | 29.4 | Good | Group | 21 |
| 22071704LJ | Figure 5 | C57BL6/J background | Ai14 het | Female | None | 17 | 20.9 | Good | Group | 21 |
| 22071705LJ | Figure 5 | C57BL6/J background | Ai14 het | Female | None | 16 | 25.7 | Good | Group | 20 |
| 22071706LJ | Figure 5 | C57BL6/J background | Ai14 het | Female | None | 16 | 19.2 | Good | Group | 20 |
| 22071707LJ | Figure 5 | C57BL6/J background | Ai14 het | Female | None | 16 | 23.2 | Good | Group | 20 |
| 22072109LJ | Figure 5 | C57BL6/J background | Ai14 het | Female | None | 18 | 24.9 | Good | Single | 22 |
| 18121201LJ | Figure 6, S12 and S13 | C57BL6/J background | Ai94D x ROSA:LNL:tTA homo/homo | Male | None | 8 | 24.9 | Good | Single | (Longitudinal functional two-photon imaging) |
| 18121204LJ | Figure 6, S12 and S13 | C57BL6/J background | Ai94D x ROSA:LNL:tTA homo/homo | Male | None | 15 | 24 | Good | Single | (Longitudinal functional two-photon imaging) |
| 18122802LJ | Figure 6, S12 and S13 | C57BL6/J background | Ai94D x ROSA:LNL:tTA homo/homo | Female | None | 20 | 22.6 | Good | Single | (Longitudinal functional two-photon imaging) |
